# *Insplico*: Effective computational tool for studying intron splicing order genome-wide with short and long RNA-seq reads

**DOI:** 10.1101/2022.08.15.503947

**Authors:** Andre Gohr, Antonio Torres-Méndez, Sophie Bonnal, Manuel Irimia

**Affiliations:** Centre for Genomic Regulation (CRG), The Barcelona Institute of Science and Technology, Barcelona, Spain; Universitat Pompeu Fabra (UPF), Barcelona, Spain; ICREA, Barcelona, Spain

## Abstract

Although splicing occurs largely co-transcriptionally, the order by which introns are removed does not necessarily follow the order in which they are transcribed. Whereas several genomic features are known to influence whether or not an intron is spliced before its downstream neighbour, multiple questions related to intron splicing order (ISO) remain unanswered. Here, we present *Insplico*, the first standalone tool to quantify ISO that works with both short and long read sequencing technologies. We first demonstrate its applicability and effectiveness recapitulating previously reported patterns, while unveiling overlooked biases associated with long read sequencing. We next show that ISO around individual exons is remarkably constant across cell and tissue types and even upon major spliceosomal disruption, and it is evolutionarily conserved between human and mouse brains. We also establish a set of universal features associated with ISO patterns across various animal and plant species. Finally, we used *Insplico* to investigate ISO in the context of tissue-specific exons, particularly focusing on SRRM4-dependent microexons. We found that the majority of such microexons have non-canonical ISO, in which the downstream intron is spliced first, and we revealed two potential modes of SRRM4 regulation of microexons related to their ISO and various splicing-related features. *Insplico* is available on gitlab.com/aghr/insplico.

## INTRODUCTION

Precursor mRNA (pre-mRNA) splicing is the processing step in the gene expression pathway that involves the removal of intronic sequences and ligation of exonic sequences to form mature RNAs (mRNAs). This process is carried out by the spliceosome, a complex molecular machinery that needs to be re-assembled *de novo* for every splicing reaction. By differentially selecting competing splice sites in each pre-mRNA molecule, the spliceosome can give rise to multiple mRNA isoforms per gene, a process referred to as alternative splicing. These alternative choices of splice sites are determined by intronic and exonic *cis*-acting elements and auxiliary proteins known as RNA binding proteins (RBPs). Some *trans*-acting factors have a tissue-specific expression and, therefore, contribute to the establishment of splicing regulatory networks responsible for cell specialization, particularly in the nervous system (1). The different steps of the gene expression pathway, including transcription itself, are interconnected and can form additional layers of (alternative) splicing regulation (2-4).

It is now well established in different species that the vast majority of genes undergo cotranscriptional splicing, i.e., splicing takes place as transcription is still occurring or shortly after (5-8). However, this does not necessarily imply that splicing always follows the order in which introns are transcribed, as initially thought (9). Single-gene studies as well as more recent genome-wide analyses have confirmed that intron splicing order (ISO) does not always occur linearly following the order of transcription (10-16). In fact, when considering the average internal exon as reference, the downstream intron is spliced before the upstream one nearly as often as the reverse (15).

Considering that early spliceosomal components are recruited at the time of transcription, multiple variables may influence ISO. As in alternative splicing regulation, both *cis*-acting elements – including strength of the splice sites and polypyrimidine tract, distance from branch point to the 3′ splice site (3′ ss), and other genomic features such as the size of the introns/exons and GC content (15) – and *trans*-acting factors (15), as well transcription kinetics (17-19) have been reported to affect ISO. Importantly, it is known that ISO can impact splicing decisions. For instance, mutations in an acceptor splice site in the *COL5A1* gene leads to changes in ISO, which affect the inclusion rate of the neighbouring exons (20). Moreover, the exon junction complex (EJC), deposited upon splicing completion, has been shown to impact the splice site selection in subsequent splicing events (21,22), providing one possible mechanism by which ISO can modulate splicing decisions. However, the converse, i.e., whether (alternative) splicing regulation across tissues or conditions affects ISO, remains unknown. Similarly, how alternative splicing regulation through specific *trans*-factors relates to ISO is poorly understood.

To investigate ISO across different cell and tissue types as well as regulatory conditions, and to facilitate further research on this topic, we developed *Insplico*, the first standalone tool to investigate ISO applicable to both short and long RNA sequencing reads. We demonstrate its robustness and effectiveness by comparing it to previous studies on ISO (15,16), recapitulating previous observations while revealing unappreciated biases introduced by long read sequencing. Next, we compared ISO in different cell and tissue types from multiple species (mammals, non-vertebrate model organisms and plants), identifying universal genomic features associated with different modes of ISO and showing that ISO is highly constant across cell and tissue types within a given species. Finally, we explored ISO for introns flanking microexons whose inclusion is dependent on *SRRM4* expression, and found two subsets of microexons with opposite ISO patterns and distinct features, suggesting two distinct modes of regulation.

## RESULTS

### Algorithm overview and definitions

*Insplico* works with both short Illumina reads (ideally paired-end) and long reads (Oxford Nanopore Technology [ONT] or PacBio), using as input BAM files of mapped reads together with a set of user-specified exons with flanking introns. Since splicing takes place in the nucleus during or shortly after transcription, chromatin-associated or nuclear RNA, or at least total ribo-depleted RNA, should preferably be used for studying ISO. *Insplico* automatically detects the read type (single-end or paired-end and strandness), and extracts counts of fragments mapping locally to each exonic region in different configurations (Fig. 1A). Specifically, for a given exon, all fragments mapping to the junction of that exon with any other upstream exon and to the unspliced downstream intron are labelled as upstream-first (*upfi*). Conversely, downstream-first (*dofi*) counts represent fragments that map to the unspliced upstream intron and the junction joining the exon with any other downstream exon. Besides these two types, *Insplico* quantifies fragments that map to both-unspliced (*bus*) flanking introns, as well as fragments supporting fully processed mRNA, i.e., mapping to both the upstream and downstream splice junctions (both-spliced, *bos*). Additionally, it obtains counts of fragments supporting exon inclusion (from upstream or downstream exon-exon junctions) or exon exclusion, containing a splice junction that skips the exon (Fig. 1A). With these counts, *Insplico* estimates different measures for each provided exon (Fig. 1B). The most relevant for ISO analysis of a given exon is F*upfi*, the fraction of *upfi* fragments in relation to the sum of *upfi* and *dofi* fragments. A F*upfi* close to 1 implies that the ISO of this exon is predominantly *upfi*, while F*upfi* close to 0 means it is predominantly *dofi*. In this study, we summarize the distributions of F*upfi* values for sets of exons of interest by empirical histograms (Fig 1C). In addition, for each input exon, *Insplico* provides: percent of exon inclusion (PSI) and percent of intron retention (PIR) for both flanking introns. These measures can be used to identify alternative vs. constitutive exons or exons that are specifically (mis-)regulated in a given condition (see below). Moreover, by additionally utilizing matched cytoplasmic and/or polyA-selected RNA-seq, PIR values of flanking introns can be estimated and used to remove stably retained introns, which can complicate the analysis of ISO.

**Figure 1.**
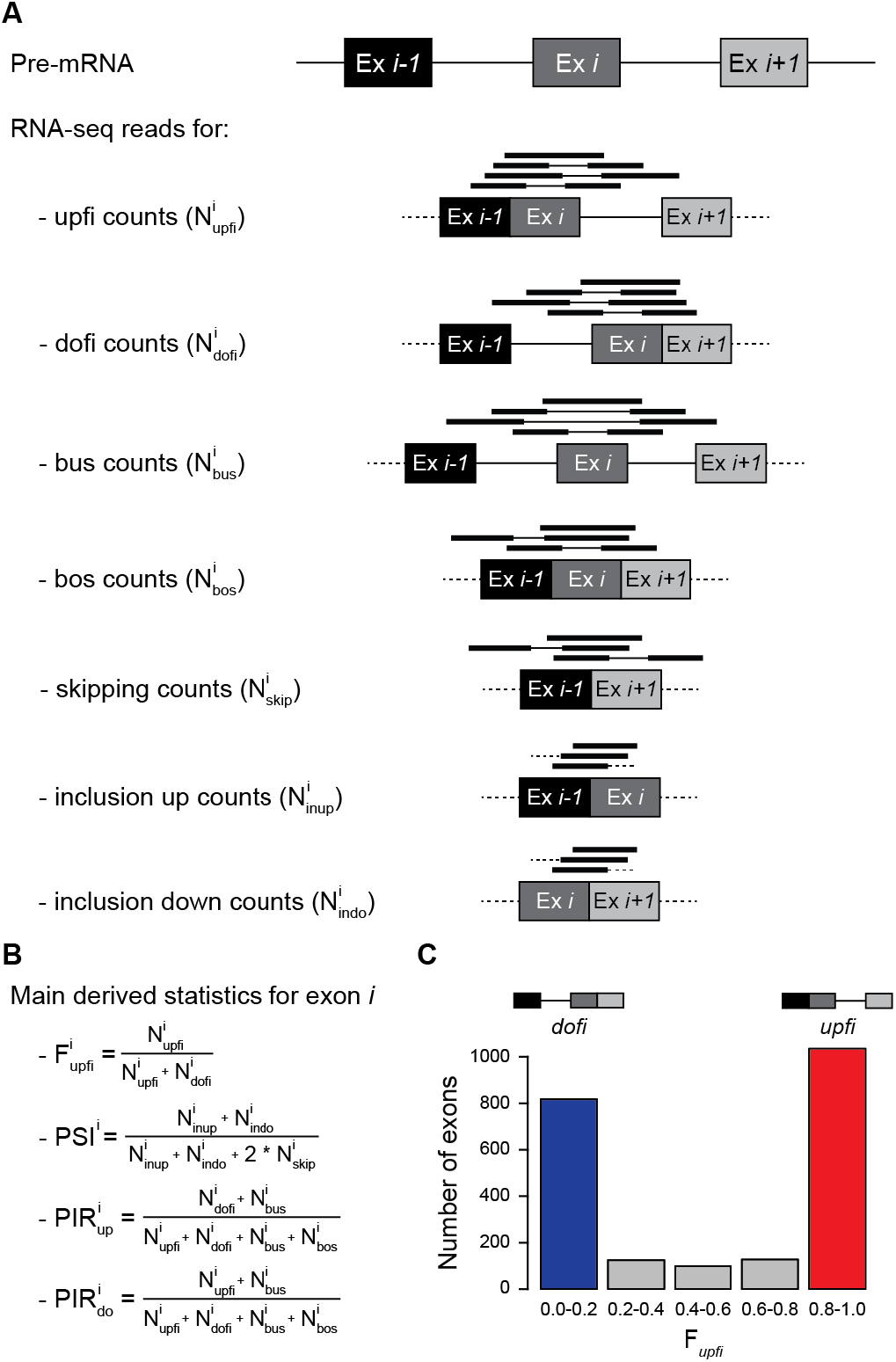
Summary of ISO and splicing related statistics provided by *Insplico*. **A**) Schematic representation of mapped short reads that are informative for each type of processing state for a specific exon (*Ex i*). These include counts for exons in which either the upstream or downstream intron has been spliced first (*upfi* and *dofi*, respectively), and for which none or both of the neighbouring introns have been spliced (*bus* and *bos*, respectively). It also includes exon-exon junction counts, for skipping or inclusion, used to derive exon inclusion levels (*skip, inup* and *indo*). **B**) Main statistics used in this study, as provided by *Insplico*. **C**) Histogram showing the distribution of a representative set of exons based on their F*upfi* values. Throughout the study, exons with F*upfi* ≥ 0.8 (red) and ≤ 0.2 (blue) are considered *upfi* and *dofi* exons, respectively.

### *Insplico* analyses of published datasets reproduce previous results

First, we tested the effectiveness of *Insplico* by applying it to the publicly available RNA-seq datasets used in Kim *et al*. (15) (Fig. 2). We downloaded and mapped the same 57 human RNA-seq samples from GEO, summing up to 7.3 billion 72-76 nt paired-end reads for non-polyA-selected total or nuclear RNA fractions from 16 different human cell lines (Table S1). We then extracted all second-first, truly internal and second-last exons from the complete Ensembl hg19 gene annotation (see Methods), and applied *Insplico* to the mapped reads to extract ISO count statistics for exons with at least 10 (*upfi*+*dofi*) counts (N(*upfi*+*dofi*) ≥ 10). In agreement with Kim *et al*., we found that, genome-wide, the majority of truly internal exons have either a clear *upfi* or *dofi* splicing pattern (F*upfi* ≥ 0.8 or ≤ 0.2, respectively), with a slight excess of *upfi* exons (Fig. 2A,B; centre plots). Second-last exons showed a stronger enrichment of *upfi* splicing patterns, again in agreement with Kim *et al*. (Fig. 2A,B; right plots). However, for second-first exons, we found a similar excess of *upfi* splicing, similar to the trend observed for second-last exons, while Kim *et al*. reported an excess of *dofi* splicing (Fig. 2A,B; left plots). To clarify this discrepancy, we further investigated the set of second-first exons as defined by Kim *et al*. using our exon-type classification derived from the Ensembl hg19 gene annotation. Remarkably, only a minority of these exons are truly second-first in all the transcripts in which they appear; the majority fell in the “diverse” exon category (Fig. 2C), which was the most common one in the annotation (Fig. 2D). Restricting the analysis to the subset of exons used by Kim *et al*. that were strictly annotated as second-first by our annotation in combination with F*upfi* values estimated by Kim *et al*. gave a pattern of ISO highly similar to that obtained with *Insplico* (Fig. 2E).

**Figure 2.**
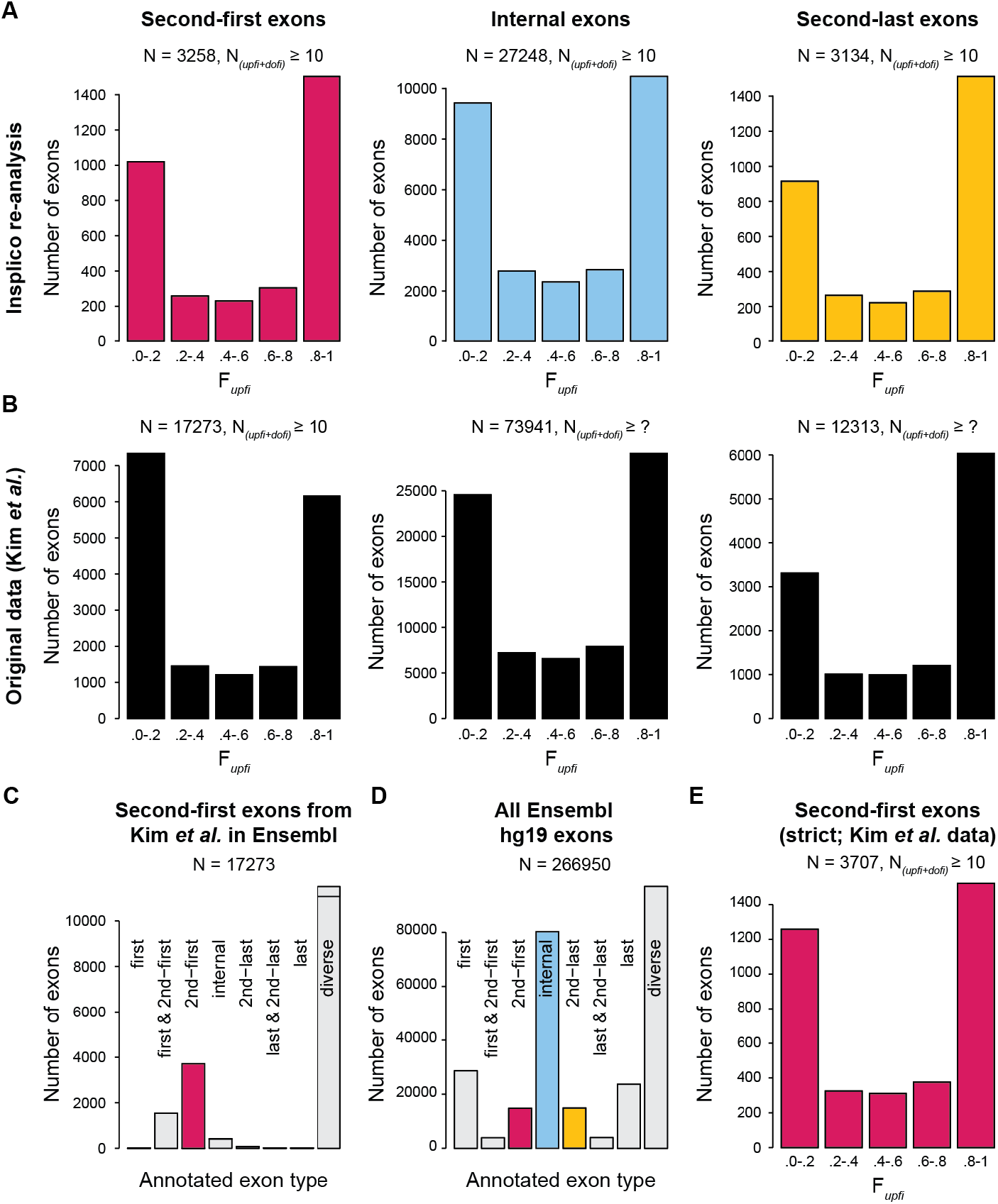
Reproduction of major ISO patterns from Kim *et al*. 2017 using *Insplico*. **A**) Distribution of exons according to F*upfi* values generated by *Insplico* for second-first (left), truly internal (middle) and second-last (right) exons as defined by *extract_exons_from_gtf*.*pl*. **B**) Distribution of exons according to F*upfi* values for second-first (left), truly internal (middle) and second-last (right) exons as provided by Kim *et al*. **C**) Distribution of second-first exons as annotated by Kim *et al*. according to exon types extracted by *extract_exons_from_gtf*.*pl*. “Diverse” correspond to exons with more than one type of annotation across transcripts. **D**) Distribution of exon types as extracted by *extract_exons_from_gtf*.*pl* for all exons in the Ensembl hg19 annotation. **E**) Distribution of strictly defined second-first exons from Kim *et al*.’s original set according to F*upfi* values as provided by Kim *et al*. (15).

### Correction of long read biases recovers short read splicing patterns

We next used RNA-seq data of nascent chromatin-associated RNA from human K562 cells published by Drexler *et al*. (16) (Fig. 3). These data consist of ∼2 million long ONT reads, which we mapped to the human genome with *Minimap2*. Applying *Insplico* to these mapped reads showed an excess of *dofi* splicing (Fig. 3A), in agreement with Drexler *et al*. (16). However, this pattern disagrees with the slight excess of *upfi* splicing described above obtained using short reads (Fig. 2; (15)). To shed light into this discrepancy, we processed with *Insplico* ∼160 million 80 nt paired-end Illumina reads from 4sU-labelled RNA also from K562 cells generated by Drexler *et al*. We observed the same slight excess of *upfi* splicing (Fig. 3B), suggesting that the nature of the sequencing data (long vs. short read sequencing) may have a considerable impact on the results. We reasoned that this discrepancy could be explained by biases introduced by long reads. First, partial pre-mRNAs could generate a bias towards *upfi* splicing for the 3′-most exon identified in the sequenced molecule, since the upstream exon will already be transcribed (and potentially spliced to the test exon) but not the downstream one, making *dofi* counts null by definition (Fig. 3C). Second, for broken RNA molecules, the 5′-most exon identified could have a bias towards *dofi* counts, since its upstream exon is not present (Fig. 3D). To reduce these potential biases in the quantification, we implemented a long read correction module (Fig. 3C,D). Correcting for each of these biases separately strongly shifted the F*upfi* distributions in the expected direction (Fig. 3C,D). Importantly, correction of both biases together retrieved a very similar distribution to the one obtained by short reads, i.e., with a slight excess of *upfi* splicing (Fig. 3C). This suggests that the discrepancies may be due to differences in the sequencing technology (i.e., long vs. short read sequencing) and that *Insplico* can correctly account for such differences through its long read correction module (option --biascorr).

**Figure 3.**
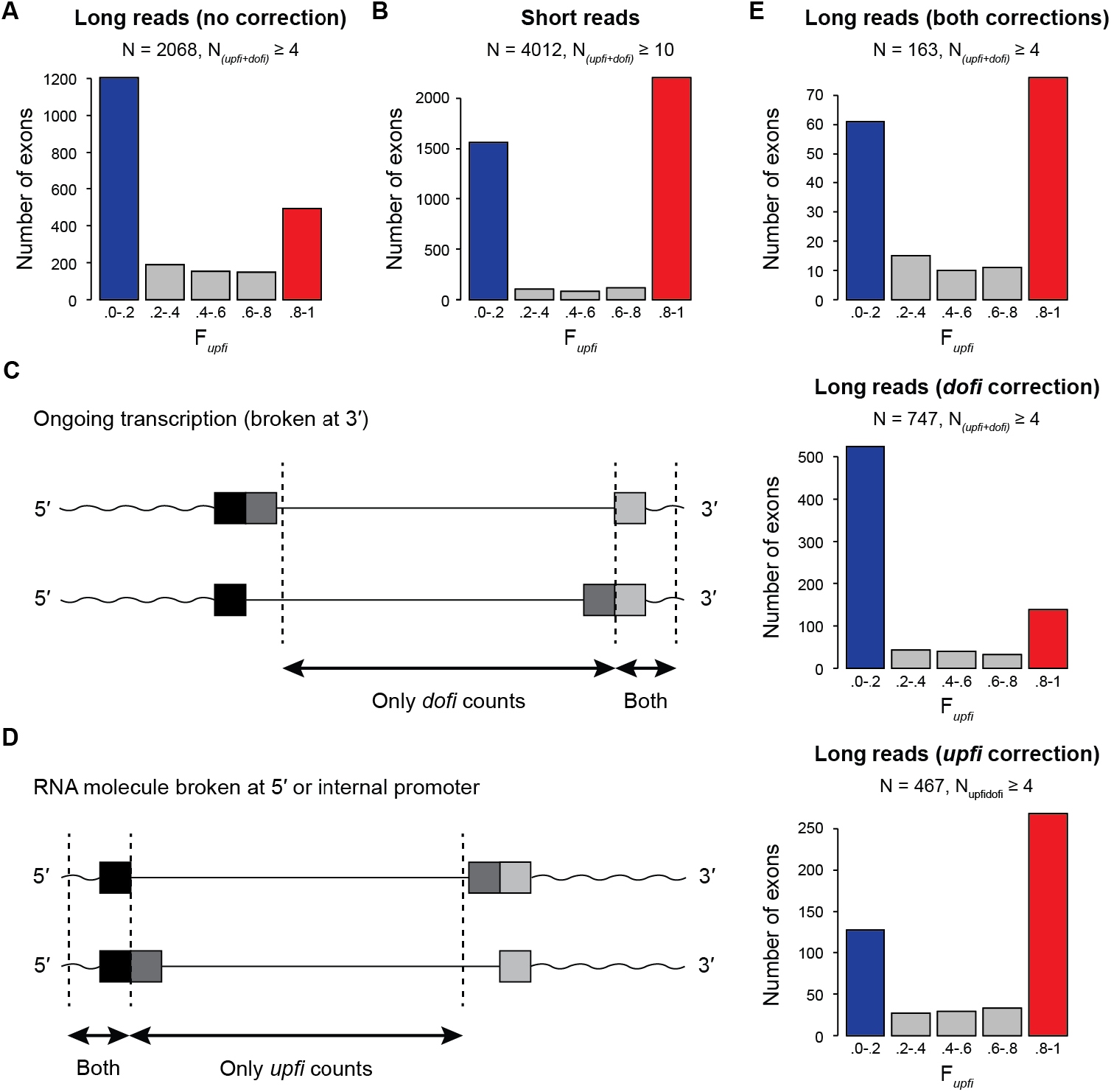
Reanalysis of ISO patterns using Drexler *et al*. 2020 ONT data. **A**) Distribution of internal exons according to F*upfi* values generated by *Insplico* using ONT reads from Drexler *et al*. (16) without correction reproduces the excess of *upfi* exons (blue) reported by the original study. **B**) Distribution of internal exons according to F*upfi* values generated by *Insplico* using Illumina short reads for the same cell type as in (A). **C**,**D**) *Insplico* corrections for ONT reads for *dofi* (C) and *upfi* (D) biases and impact of each of these corrections in the ISO profiles (right). **E**) Distribution of internal exons according to F*upfi* values generated by *Insplico* using ONT reads from Drexler *et al*. (16) with full correction reproduces the ISO profile obtained by short reads.

### Intron splicing order is highly stable across different biological conditions

Despite these consistent genome-wide patterns, the ISO of a given exon may vary across different cell/tissue types or conditions, a possibility that has not been investigated yet. To address this, we compared exons with strong *upfi* and *dofi* splicing (F*upfi* ≥ 0.8 or ≤ 0.2, respectively), in various pairs of deep chromatin-associated RNA-seq samples. We reasoned that if ISO is widely maintained between the two samples, the sets of *upfi* (red) and *dofi* (blue) exons in the first sample will tend to be *upfi* and *dofi*, respectively, in the second sample (“consistent”, Fig. 4A). Otherwise, the background distribution will be observed in the second sample for each ISO type in the first sample (“random”, Fig. 4A). First, we investigated the consistency of ISO patterns between replicates of the same studies for exons with sufficient informative reads (N*(upfi+dofi)* ≥ 10) in the two compared samples. Biological replicates of either human K562 (8) or HeLa (23) cells showed extremely consistent ISO patterns (Fig. 4B,C). Second, we compared the patterns in K562 and HeLa cells, which also exhibited very consistent ISO distributions (Fig. 4D). Third, we aimed at comparing cells of very different origin and function. For this purpose, we generated deep chromatin-bound RNA-seq data from mouse brain (cortex and cerebellum) and liver (Table S1), as well as paired cytoplasmic polyA-selected RNA-seq to identify and discard potential intron retention events. Remarkably, *upfi* and *dofi* exons in the brain showed nearly identical patterns in the liver (Fig. 4E), suggesting that ISO patterns are highly consistent even across very divergent cell types. Moreover, we assessed if these patterns were conserved between species by comparing ISO between human and mouse brains (Fig. 4F). Although the consistency was not as strong as within each species, we found significant conservation of ISO patterns: of 522 *upfi* exons in human, 347 are also *upfi* in mouse (67%; *P* = 2.1e-14, one-sided Binomial test), and of 397 *dofi* human exons, 287 conserved their ISO in mouse (72%; *P* = 1.3e-19, one-sided Binomial test). Finally, we investigated whether ISO consistency may be disrupted upon major spliceosomal interference. We used RNA-seq of 5-Bromouridine (BrU)-labelled RNA from HeLa cells treated with two SF3B1-targeting drugs, spliceostatin (SSA) and sudemycin C1 (SudC1), which cause widespread intron retention and exon skipping (24). Even under these conditions, ISO was largely maintained with respect to DMSO-treated control cells (Fig. 4G,H), including the ISO patterns around exons whose immediately upstream and downstream introns were both substantially affected by the treatments (ΔPIR (Treatment – DMSO) ≥ 0.25; Fig. 4I).

**Figure 4.**
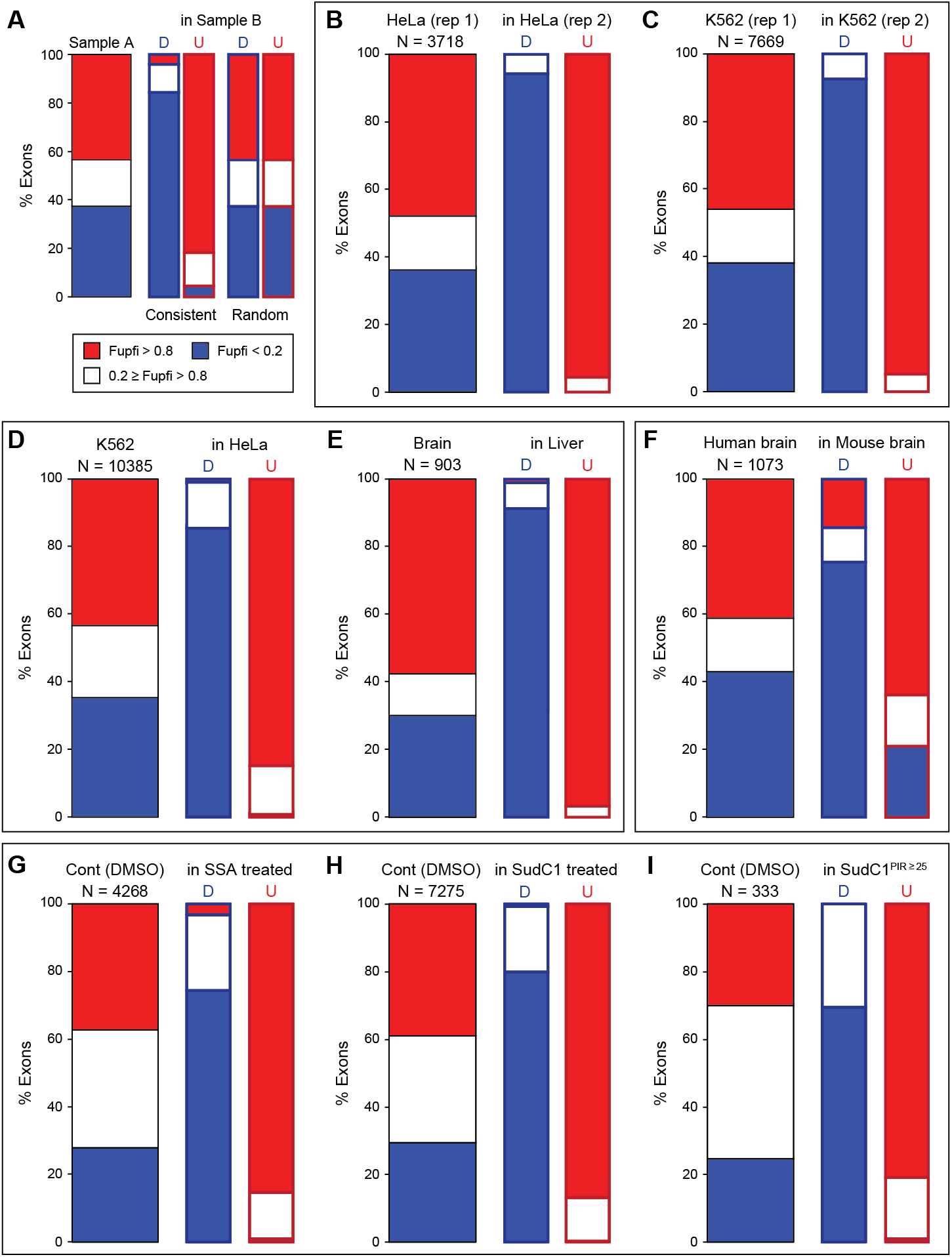
ISO patterns are largely constant across cell and tissue types and experimental conditions. **A**) Schematic summary of the analysis. For a given Sample A, its ISO profile is derived showing the proportion of exons that are strongly *upfi* (blue), *dofi* (red) or intermediate (white). Then, *upfi* and *dofi* exons are separately interrogated in a different Sample B, and their ISO profiles derived in the same manner. Two extreme patterns can be envisioned: “Consistent”, in which most *upfi* exons in Sample A will also be *upfi* in Sample B (and the same for *dofi* exons), and “Random”, in which *upfi* and *dofi* exons separately will reproduce the background distribution of Sample A in Sample B. **B**,**C**) Consistent ISO patterns across replicates in HeLa cells (B, data from (23)) and in K562 cells (C, data from (8)). **D**,**E**) Consistent ISO patterns between human HeLa and K562 cells (D) and mouse brain and liver tissues (E, data from this study). **F**) Significant evolutionary conservation of ISO patterns among orthologous exons in human and mouse brain (data from (39) and this study). **G**,**H**,**I**) Consistent ISO patterns in HeLa cells treated with DMSO (control) or spliceostatin (SSA; G) or sudemycin C1 (SudC1; H). ISO patterns were largely consistent even for exons for which both neighbouring introns were highly affected by drug treatment (ΔPIR > 0.25; I). Data from (24).

### Universal genome-wide features of intron splicing order

Given the stability of ISO across different biological conditions, it is likely that its main determinants are to a large extent hardcoded in the genome. Consistently, different sequence features have been previously shown to influence this process (see Introduction). To further identify universal features across conditions and species, we separately processed with *Insplico* 13 short read RNA-seq datasets from nuclear or chromatin-associated RNA fractions from four animals (human, mouse, fruitfly and round worm) and two plants (Arabidopsis and rice) (Table S1), which have very different intron densities, genome architectures and alternative splicing patterns (25,26). From each dataset, we considered all truly internal exons with sufficient informative reads, computed their ISO pattern, and extracted 55 splicing-related genomic features using *Matt* (27) (see Methods). To assess the possible contribution of each of these features on ISO, we partitioned for each feature the exons into five equal-sized subsets according to increasing feature values, and plotted the average F*upfi* value per bin (Fig. 5A and Supplementary File 1). We identified several genomic features that systematically, linearly or nonlinearly, correlate with ISO with the same functional relationship across all datasets from all species (Fig. 5B,C). Specifically, exons whose upstream flanking intron is removed first (*upfi* exons) have well-defined upstream introns with: (i) strong 5′ ss, (ii) strong 3′ ss, and (iii) branch points (BPs) close to their 3′ ss (AG). On the other hand, their downstream introns are (iv) long, (v) particularly respect to the upstream intron, and (vi) their BP is far from their 3′ ss. Altogether, these results highlight the importance of a strongly defined upstream intron as well as the length and the BP-AG distance of both flanking introns relative to other sequence features.

**Figure 5.**
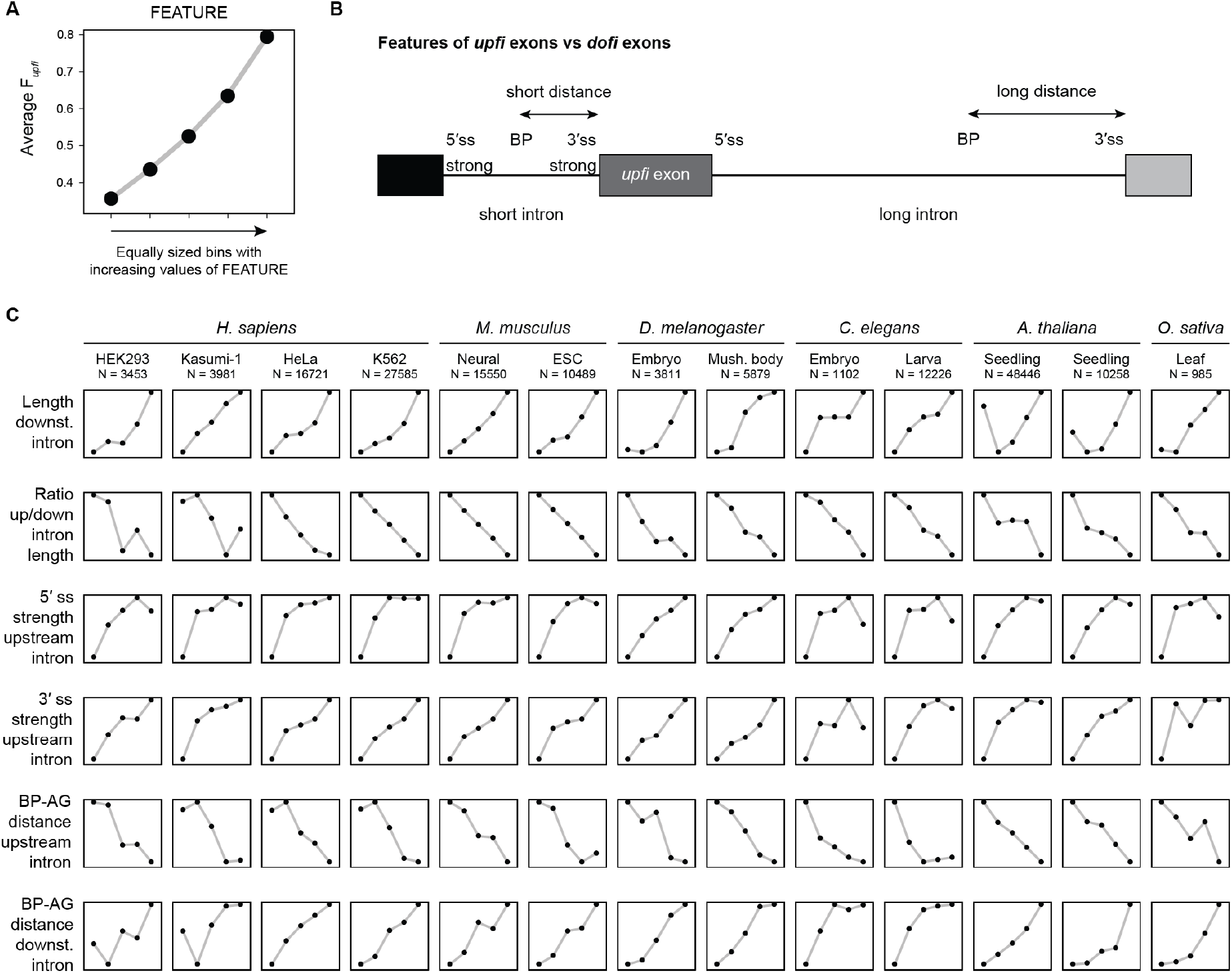
Genomic features universally associated with ISO. **A**) Schematic representation of the plots obtained in this analysis. For a given feature, the average of F*upfi* values (Y axis) is shown for five equal-sized groups of exons binned by increasing values of that feature (X axis). Consistently increasing or decreasing associations are considered. **B**) Summary representation of the features consistently associated with exons whose upstream intron is spliced first. BP, branch point; ss, splice site. **C**) Summary plots for the main features universally associated with ISO patterns across samples and species. Further details for all features are provided in Supplementary File 1.

### Splicing of *SRRM4*-dependent neural microexons is predominantly *dofi*

Given these consistent genomic features correlating with ISO and the global consistency in the patterns across cell types, we asked how tissue-specific regulation of alternative splicing relates to ISO. Previous studies have shown that alternatively spliced exons are more often *dofi*, compared to constitutive exons (15). Interestingly, by separating mouse alternatively spliced exons (0.1 < PSI < 0.9 in liver and/or brain) into those that are or that are not regulated in a tissue-dependent manner (|ΔPSI liver vs. neural| ≥ 0.25), we found that the excess of *dofi* is only substantial and statistically significant for tissue-regulated (TR) exons and not for all alternatively spliced exons (Fig. 6A). However, when looking at the key genomic features identified above for TR exons we found a mix of patterns, indicating that their genomic features cannot solely explain the excess of *dofi* splicing (Fig. 6B and Supplementary File 2).

**Figure 6.**
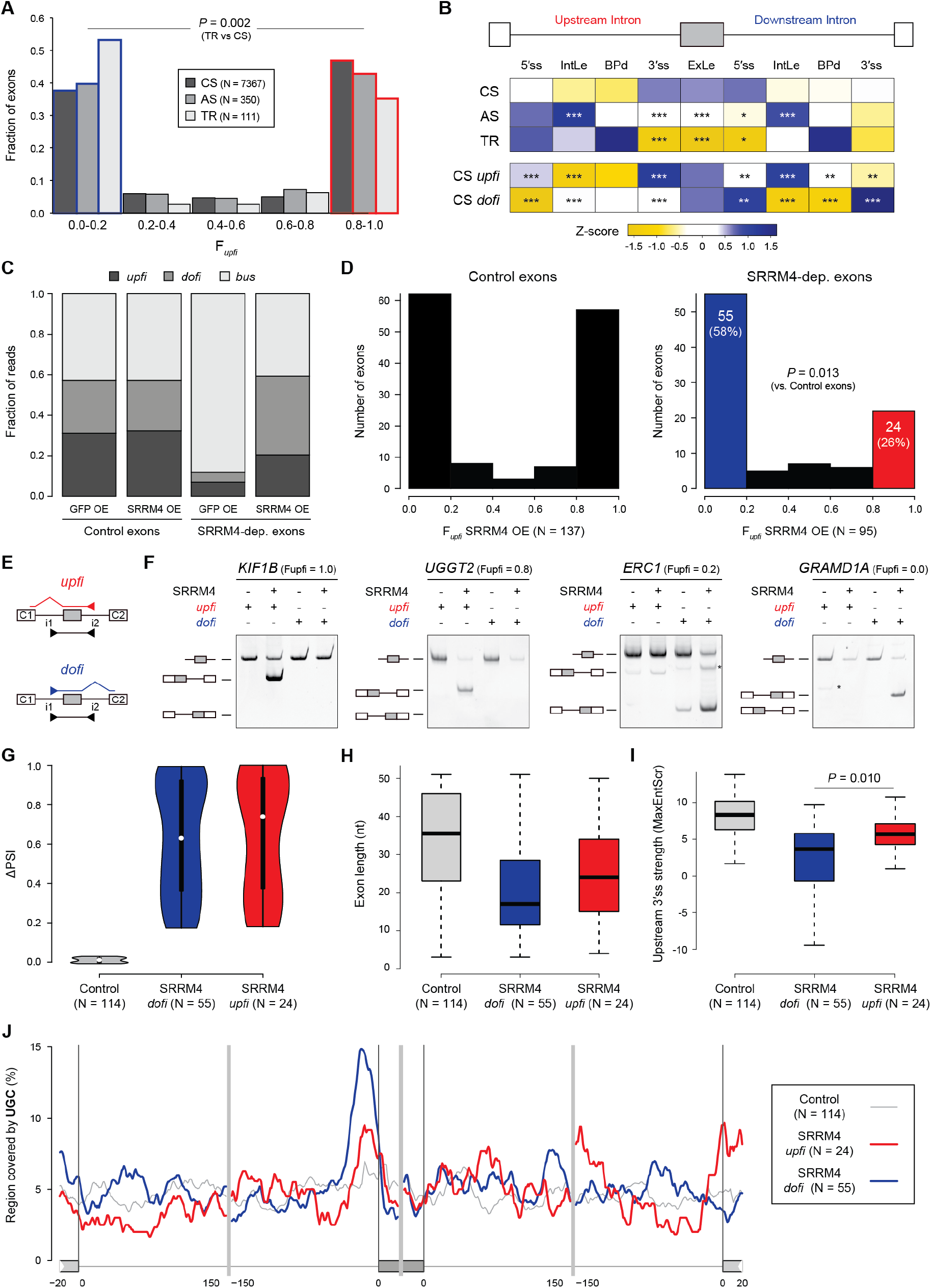
ISO patterns differentiate two subsets of SRRM4-dependent microexons. **A**) Distribution of constitutive (CS), alternative (AS) and tissue-regulated (TR) exons according to F*upfi* values. P-value corresponds to the comparison between *upfi* (red) and *dofi* (blue) TR vs. CS exons using a two-sided Fisher’s Exact test. **B**) Z-scored median values for each feature for each exon type as well as CS exons that are strongly *upfi* (“CS *upfi*”) and *dofi* (“CS *dofi*”). IntLe, intron length; BPd, distance from branch point to 3′ ss; ExLe, exon length. P-values correspond to Bonferroni-corrected Wilcoxon Rank-Sum tests against the CS distributions for each feature. * 0.05 < *P* ≤ 0.01, ** 0.01 < *P* ≤ 0.001, *** *P* < 0.001. All details for all features are provided in Supplementary File 2. **C**) Distribution of unprocessed reads (*upfi, dofi* and *bus*) for control and *SRRM4*-dependent (SRRM4-dep.) microexons in each condition. **D**) Distribution of exons according to F*upfi* values for control (left) and *SRRM4*-dependent (right) microexons in HEK293 cells ectopically expressing human *SRRM4*. P-value corresponds to the comparison between *upfi* (red) and *dofi* (blue) control and *SRRM4*-dependent exons using a two-sided Fisher’s Exact test. **E**,**F**) Validation of ISO for four *SRRM4*-dependent microexons with *upfi* or *dofi* patterns in *SRRM4*-expressing HEK293 cells. Primers in upstream and downstream introns (i1 and i2) in combination with exon junction primers between the exon C1 upstream and the microexon (*upfi*, red) and between the microexon and the exon C2 downstream (*dofi*, blue) were used for specific detection of partially processed transcripts, as depicted in E. Unidentified amplification products in F are indicated with an asterisk. **G**,**H**,**I**) Distributions of ΔPSI (SRRM4 OE vs control; G), exon length (H) and 3′ ss strength of the upstream intron (using the MaxEntScr metric; I) for control and *SRRM4*-dependent exons with strong *dofi* (blue) and *upfi* (red) patterns. P-value in I corresponds to a Wilcoxon Rank-Sum test. **J**) RNA map showing the percent sequence covered by UGC motifs using a sliding window of 25 nts for each microexon type. For C-J, only microexons with *N(upfi+dofi)* ≥ 5 in the SRRM4 OE sample were considered for all the analyses.

Therefore, we hypothesized that their tissue-specific splicing regulators may impose unique regulatory architectures with non-canonical ISO patterns. To begin assessing this possibility, we focused on a particular case of extreme tissue-specific regulation: neural-specific microexons that are dependent on the splicing factor *SRRM4* for their inclusion. The mode of action of the splicing factor *SRRM4* is not fully understood, but interactions between its major functional protein domain, the eMIC domain, and early spliceosomal components have been demonstrated (28,29). To shed more light into the mechanism of *SRRM4*-dependent microexon inclusion, we generated total ribo-depleted RNA-seq data from human HEK293 cells ectopically expressing either GFP or SRRM4 and applied *Insplico* for PSI and F*upfi* estimations. We identified 677 exons that were more included upon SRRM4 overexpression (OE) compared to the control (ΔPSI ≥ 0.15) and that were lowly included in control cells (PSI < 0.2). As expected, most of these (511/677, 76%) were microexons (defined here as length ≤ 51 nt). We then compared these *SRRM4*-dependent microexons with a control exon set of length ≤ 51 nt, high inclusion in the control (PSI > 0.8) and not affected by SRRM4 OE (|ΔPSI| < 0.1). As expected, by looking at all *Insplico* counts for unprocessed transcripts (i.e., *upfi, dofi, bus*; Fig. 1), we found that *SRRM4*-dependent microexons need SRRM4 even for partial processing (Fig. 6C). Remarkably, in SRRM4 OE cells, we found that 58% of *SRRM4*-dependent microexons had strongly *dofi* splicing patterns (F*upfi* ≤ 0.2), while only 26% of them seem to be spliced in an *upfi* manner (F*upfi* ≥ 0.8; Fig. 6D), in contrast to roughly equal percentages in the control set (*P* = 0.013, Fisher’s Exact test). Importantly, RNA-seq-based ISO patterns were validated experimentally using *upfi* and *dofi* specific primers for four *SRRM4*-dependent microexons (Fig. 6E,F).

We next asked whether *upfi* and *dofi SRRM4*-dependent microexons showed distinct characteristics. First, being *upfi* or *dofi* did not impact the magnitude of the response to SRRM4 OE (Fig. 6G). However, analysis of intron-exon features with *Matt* revealed several genomic differences between *upfi* and *dofi* SRRM4-dependent microexons (Supplementary File 3). Interestingly, despite all being microexons by definition, exon length was a relevant discriminating feature, since *dofi* microexons tended to be shorter than *upfi* microexons (medians 17 vs 24 nts, respectively) (Fig. 6H). In addition, another major feature previously associated with *SRRM4*-dependent microexons, weaker 3′ ss contexts in the upstream intron (28-30), was much more prominent for *dofi* microexons (Fig. 6I), consistent with the general pattern of all *dofi* exons (Figs. 5 and 6B). Finally, the presence of the UGC motif near the 3′ ss, characteristic of eMIC-dependent regulation by SRRM4 (28-31), was much more prevalent in *dofi SRRM4*-dependent microexons (Fig. 6J). In summary, these results reveal that the previously reported features associated with SRRM4 regulation are chiefly associated with *dofi* splicing and raise the possibility that two modes of regulation by SRRM4 may exist in connection with opposed ISO patterns.

## DISCUSSION

We developed *Insplico*, the first standalone software dedicated to the study of the order of intron splicing applicable to both short and long read sequencing technologies, and demonstrated its effectiveness by comparing it with two previous studies of ISO (15,16). Importantly, despite specifically studying intron splicing, *Insplico* is an exon-centric, not intron-centric tool. It obtains key measurements to investigate pre-mRNA processing around the exons of interest (*upfi, dofi, bus* and *bos* counts) and estimates the fraction of transcripts in which the upstream intron is spliced before the downstream one (F*upfi*). Moreover, *Insplico* quantifies the level of inclusion of exons (PSI) and the level of intron retention (PIR) of the two neighbouring introns, allowing more complex integrated analyses as well as further filtering and stratification strategies within a single software.

Contrary to short reads, which only give local information, long reads have the potential to provide a snapshot of all introns of a transcript at once, thus being particularly promising to study ISO. Intriguingly, a pioneer study (16) obtained ISO patterns in humans using long reads that did not match those previously reported for short reads. Here, we showed that such discrepancies could be due to 5′ and 3′ biases in long read sequencing and library preparation that, when corrected for, harmonise the patterns obtained by both types of sequencing technologies. The module to perform these corrections is a unique feature of *Insplico*. On the other hand, short read sequencing has its own limitations. For example, given that *upfi* and *dofi* fragments require mapping across the entire exon to cover both splice sites (Fig. 1A), the length of the reads (and the fragment size of paired-end reads) is expected to limit the length of the exons that are quantifiable. Moreover, although we have explicitly focused on the relative ISO of the pairs of introns directly flanking the tested exons, short reads cannot provide ISO information beyond them.

After demonstrating its effectiveness, we applied *Insplico* to multiple public and newly generated RNA-seq datasets to gain insights into ISO across cell types, conditions and species. Splicing is known to mainly occur co-transcriptionally and be influenced by transcriptional related features, RNA *cis*-acting elements and binding proteins. How these factors determine ISO and how it varies across different conditions is largely unknown. Remarkably, through our analyses, we not only observed a strong consistency of ISO across cell and tissue types from the same species, but also even under disruptive spliceosomal conditions such as those induced by SF3B1-targeting drugs. These findings thus suggest that, even though experimentally induced changes in ISO have been shown to affect splicing outcomes (20,22), ISO is largely independent of the splicing outcome under physiological and non-physiological conditions. Moreover, these results argue that, for most exons, the main contributors to this predefined ISO are likely hardcoded in the genome. In line with this idea, our multi-species analysis revealed several universal genomic features that are strongly associated with exons whose upstream intron is spliced before the downstream one across various animal and plant species. In particular, such exons are typically flanked by a short upstream intron with strong 5′ ss and 3′ ss (including a long polypyrimidine tract and short BP-AG distance) and a long downstream intron with large BP-AG distance, but otherwise regular splice sites. Those features are in line with and expand previous studies (15,16,32), and are consistent with the model known as “first come, first spliced”, whereby a cotranscriptional recruitment of the spliceosome occurs and triggers the splicing of well-defined introns as soon as they are fully transcribed (10-14). However, consistent with previous studies (13,15,16), we also found that there are nearly as many exons whose downstream introns are spliced prior to the upstream one as there are of the converse “canonical” case, overall resulting in a strong bimodal distribution of F*upfi* values genome-wide for all studied species. This large number of *dofi* exons, which are particularly associated with tissue-regulated alternative exons, is consistent with previous studies indicating large fractions of delayed or post-transcriptional splicing (33) and the exceptional dependence of these introns on specific *trans*-acting factors for their splicing (1,34).

Along these lines, we focused here on a unique case of tissue-regulated exons, neural microexons, which are characterised by their short length (defined here as smaller than 51 nt) and dependence on SRRM4 for inclusion (30). An appealing hypothesis, which we investigated here, is that these microexons are highly dependent on a particular ISO of their associated introns for inclusion. Along this line and consistent with the pattern of ISO for TR exons, we observed that the majority of *SRRM4*-dependent microexons are preferentially spliced in a *dofi* manner. Such *dofi* microexons are shorter, have weaker 3′ ss, and much more marked UGC peak at a shorter distance to the 5′ ss than *upfi* ones. This suggests that the simultaneous assembly of U1 and U2 snRNPs is not possible on the surrounding 5′ ss and 3′ ss splice sites of these microexons due to steric hindrance. Therefore, it could be possible that they are mainly recognized at the level of their 5′ ss, which are known to be particularly strong (28,30), by the U1 snRNP whose recruitment could then favour splicing of the downstream intron. After splicing of the downstream intron and, possibly, recruitment of RNPS1 (component of the Exon Junction Complex) (35), SRRM4 binding could favour 3′ ss recognition by promoting the recruitment of early spliceosomal components (28) and subsequent splicing of the upstream intron. Intriguingly, however, a non-negligible fraction of *SRRM4*-dependent microexons are spliced in an *upfi* manner. Those tend to be longer, with stronger 3′ ss and a less marked UGC peak, suggesting that they may be recognized and spliced in a way more similar to constitutive exons. Although those hypotheses remain to be formally tested, our ISO analysis thus points at two possible mechanisms of splicing of *SRRM4*-dependent neural microexons.

## MATERIAL AND METHODS

### *Insplico*: input data generation and exon definition

*Insplico* takes as input mapped RNA-seq reads in BAM format. These must have been aligned with a splice-aware mapper, eg. STAR or HISAT2 for short reads, or with Minimap2 for ONT/PacBio long reads. Furthermore, *Insplico* needs a tab-separated table defining the exons for which read statistics are desired. Exons are defined by their start and end coordinates, strand, and the end/start coordinate(s) of their upstream/downstream neighbour exons, respectively. More details on this exon-defining table can be found on gitlab.com/aghr/insplico. In addition, a script (*extract_exons_from_gtf*.*pl*) that allows users to create such a suitable table from a gene annotation file in GTF format is provided. This script clusters overlapping exons from several transcripts of the same gene into complex exon entities with potentially several start and end coordinates (following the logic of *vast-tools* (36)) and it implements a heuristic to identify intron-retention events that are not considered true exons. In addition, it identifies and assigns an exon type to each exon (Fig. 2C,D), e.g. *sfrst* (second first), if the exon is the second-first exon in all transcripts where it appears, *slst* (second last), if it is always the second-last exon, or *diverse* if it appears in different positions in transcripts. From the exon table, *Insplico* extracts for each exon the set of start coordinates, the set of end coordinates, the strand, and the upstream and downstream intron regions. These regions are defined by the intronic region between the exon and its direct upstream and downstream neighbour exons while considering the maximal extent of these exons. Importantly, to reduce potential bias stemming from upstream and downstream introns of different lengths, *Insplico* utilizes as effective length of both introns the length of the shorter intron.

### *Insplico*: algorithmic details

*Insplico* is implemented in Perl, uses exclusively Perl libraries shipped with the standard installation of Perl and depends on Samtools (37) and Bedtools. As such, it can be run readily after download on Unix-like systems where Perl, Samtools, and Bedtools are available. Considering all exon starts and ends and the upstream and downstream intronic regions of same length, *Insplico* inspects the mapped reads to identify those in *upfi, dofi, bos* and *bus* configurations (Fig. 1), as well as the counts for exon skipping, inclusion upstream and inclusion downstream. These read count statistics, together with estimates of F*upfi* (fraction of *upfi* reads over the total number of *upfi*+*dofi* reads), percent of exon inclusion (using the standard percent-spliced-in metric, PSI) and percent of intron retention (PIR) of the upstream and downstream introns, are output as a tab-separated table where rows (exons) are ordered correspondingly to the rows of the input table. Unavailable estimates are indicated by NA.

Three features of *Insplico*’s implementation facilitate its usability: (i) *Insplico* works out-of-the-box with mapped short and long reads. Algorithmically, long reads are treated identically to single-end short reads as there exists no conceptual difference between these two; (ii) users do not need to define the strandness of the RNA-seq reads, which is often a cumbersome matter for incompletely documented RNA-seq data; and (iii) users can use coordinates in 0- or 1-based genetic coordinate systems without the need to change the reference system because of fuzzy coordinate matching. When comparing the splice site coordinates of reads to exon starts and ends, *Insplico* can apply a user-definable fuzziness to detect matching coordinates by default up to +/-3 nt. Fuzzy matching of coordinates is useful when ambiguities in splice site mapping occur and *Insplico* is capable of rescuing such reads for its count statistics. Furthermore, *Insplico* will auto-detect the read strandness and consider it to reduce potential corruption of read statistics caused by reads mapping to the opposite strand.

Another capability of *Insplico* that sets it apart from previous approaches is the bias correction for processing long reads such as ONT or PacBio. When these reads do not cover the entire transcript, from transcription start site to termination, then they can cause a bias of the *upfi* and *dofi* count-statistics extracted for the first and last internal exons covered by the reads. *Insplico* offers a reduction of this bias, which can be activated by the user (see below). Further details on *Insplico*, available command arguments and applications can be found on gitlab.com/aghr/insplico.

### Comparative re-analysis of published data (Kim *et al*., 2017)

GEO IDs of all 57 RNA-seq data sets used by Kim *et al*. (15) from total or nuclear RNA without poly-selection are listed in Table S1. After download, we removed the Illumina universal adapter AGATCGGAAGAGC with cutadapt v2.4 from the 3′ ends of read1 and read2, keeping only reads with a minimum length of 15 nt. As per the original study, these reads were mapped to the human hg19 genome assembly with the splice-aware mapper STAR v2.7.1a, requiring a minimal overlap of 5 nt on both sides of splice junctions and keeping only uniquely mapping reads. The resulting BAM files with mapped reads were analyzed with *Insplico*. To create the exon definition table, we applied the script *extract_exons_from_gtf*.*pl* to the Ensembl v85 hg19 gene annotation GTF, which gave 266,950 exons together with their exon types across all transcripts where they appear. The most prominent exon types were diverse/mixed (39.2%), internal (30.1%), first (10.7%) and last (8.9%), as shown in Fig. 2D. We discarded exons of the “diverse” type as they may introduce noise, and focused on second-first, internal and second-last exons. Only exons with a minimum of ten *upfi*+*dofi* reads (N(*upfi+dofi)* ≥ 10) were included in the analysis. To directly compare these results with those published by Kim *et al*., we downloaded the tables with their results as provided on http://fairbrother.biomed.brown.edu/data/Order. The table first_introns_splicing_pair_counts.txt contained a list of 43,547 second-first exons as identified by Kim *et al*. with F*upfi* values, of which 17,273 had N(*upfi+*d*ofi*) ≥ 10. The tables middle_intron_scores.txt and last_intron_scores.txt contained 73,941 internal and 12,313 second-last exons, respectively, all of which were plotted since N(*upfi+dofi)* was not provided. Finally, to investigate the cause of the difference in ISO profiles obtained by our and Kim’s et al. analysis for second-first exons, we matched the 17,273 second-first exons identified by Kim *et al*. to the exons types we extracted from the Ensembl hg19 gene annotation. We found that most of these exons are of the “diverse” type. When plotting the profile for those of Kim’s exons that are strictly defined as second-first exons by our annotation, we obtained a pattern of slight upfi excess consistent with our re-analysis with *Insplico*.

### Comparative re-analysis of published data (Drexler *et al*., 2020) and long read bias correction

Drexler *et al*. (16) sequenced chromatin, 4sU enriched RNA from K562 cells using ONT. We downloaded this dataset and mapped it to the human genome hg38 assembly with the Minimap2 v2.17-r974-dirty (38) in splice mode with seeds of length 14 nts. To create the exon input table, we applied the script *extract_exons_from_gtf*.*pl* to the gene annotation GTF from Ensembl v88, together with all exons from VastDB (36), obtaining a total of 233,306 unique exons. The BAM file from Minimap2 and the exon table were used to run with *Insplico* in standard mode to extract raw read count statistics. A histogram of F*upfi* values was plotted for all exons with N(*upfi+*d*ofi*) ≥ 4. In addition, we downloaded Illumina paired-end short reads for chromatin, 4sU-enriched RNA also generated by Drexler *et al*. (Table S1). We removed the Illumina universal adapter AGATCGGAAGAGC with cutadapt v2.4 from the 3′ ends of read1 and read2, and kept reads with a minimum length of 15 nts. These reads were mapped to the same human hg38 assembly with STAR v2.7.1a, requiring a minimal overlap of 5 nts on both sides of splice junctions and keeping only uniquely mapping reads. We then ran *Insplico* on the resulting BAM files with the same exon input table used for the analysis of ONT long reads and generated F*upfi* histograms for all exons with N(*upfi+*d*ofi*) ≥ 10. Given the different profiles obtained for short and long reads, we hypothesised these could be caused by specific biases introduced by ONT with respect to Illumina data. Thus, we ran *Insplico* again for the ONT data applying the bias correction for the 5’ end, 3’ end, or both ends together. The *dofi*/*upfi* corrections consisted in effectively removing the first and last exons that are covered by a given ONT read, respectively. Since these exons are not considered, this implies that the following and preceding exons cannot receive neither *upfi* nor *dofi* counts. Therefore, under the bias correction, *Insplico* focuses only on the subsequent internal exons mapped by the specific ONT read (further algorithmic information can be found on gitlab.com/aghr/insplico). Each correction shifted the *Fupfi* distribution in the expected direction and the double correction provided a profile consistent with that of the short reads.

### Consistency of ISO across different replicates, tissues, experimental conditions and species

For studying ISO across different replicates, cell types, tissues and experimental conditions, we first obtained exon input tables for human hg38 and mouse mm10 Ensembl v88 annotations enriched with VastDB exons, as described above. We then downloaded RNA-seq datasets from various sources (Table S1), removed the Illumina universal adapter AGATCGGAAGAGC with cutadapt v2.4 from the 3′ ends of read1 and read2, kept the reads with length ≥ 15 nts, and mapped them to the respective genomes using STAR v2.7.1a requiring a minimal overlap of 5 nts on both sides of splice junctions and keeping only uniquely mapping reads. These BAM files were used to run *Insplico*. In addition, for the datasets from Tilgner *et al*. (8), Ke *et al*. (23), Burke *et al*. (39), as well as the mouse tissue-specific data we generated, we also mapped and analysed with *Insplico* the associated polyA-selected and/or cytoplasmic RNA samples, and utilized the estimated PIR values to filter out exons associated with retained introns (PIR ≤ 0.15 for the upstream or downstream intron), and only truly internal exons (as defined above) were used for these comparisons. Then, to compare ISO patterns between two conditions (biological replicates, different cell types, different species), we first selected exons with N(*upfi+dofi)* ≥ 10 in both conditions. Next, we defined *upfi* or *dofi* exons in the query condition (left) as those with F*upfi* ≥ 0.8 and F*upfi* ≤ 0.2, respectively, and investigated what fraction of each of them was *upfi, dofi* or had intermediate F*upfi* values in the target condition (right). As explained in Fig. 4A, if ISO is similar in both conditions (“Consistent”), most *upfi* exons of the query condition will also be *upfi* in the target condition (and the same for *dofi* exons). On the other hand, if ISO patterns are not maintained between conditions (“Random”), the *upfi* and *dofi* exon sets in the query condition should have an ISO pattern similar to the genome-wide pattern in the target condition.

### Universal features associated with ISO patterns

To investigate which intron-exon related features affected ISO genome-wide across multiple species, we first selected 13 RNA-seq datasets from different species (Table S1) and processed them with *Insplico*. As a standard procedure, we removed the Illumina universal adapter AGATCGGAAGAGC with cutadapt v2.4 from the 3′ ends of read1 and read2, keeping only reads with length ≥ 15 nts. These reads were mapped to the corresponding genomes of each species with the splice-aware mapper STAR v2.7.1a requiring a minimal overlap of 5 nts on both sides of splice junctions and keeping only uniquely mapping reads. The STAR index was built considering the gene annotations for each species and we obtained the exon-definition table applying *extract_exons_from_gtf*.*pl* to each GTF file together with exons annotated in *vast-tools* (36), except for rice (not available). Specifically, we used the following species and genome versions: *Homo sapiens* (hg38, Ensembl v88), *M. musculus* (mm10, Ensembl v88), *D. melanogaster* (dm6, Ensembl Metazoa v26), *C. elegans* (ce11, Ensembl v87), *A. thaliana* (araTha10, Ensembl Plants v31) and *O. sativa* (IRGSP1, Ensembl Plants v48).

Next, we plotted the distribution of F*upfi* values for subsets of exons according to multiple intron-exon related features extracted using *Matt* (27). Specifically, we used the *Matt* command *cmpr_features*, and, for each studied feature, we split the exons into five subsets of equal size with increasing feature value. These plots allow us to study how the F*upfi* pattern changes for subsets of exons with different feature values. A summary of these plots for those features consistently and significantly associated with ISO across species is shown in Fig. 5c for all datasets. All the results, together with further details (axis values, sample sizes, etc.) can be found in Supplementary File 1. In general, for all datasets, we only considered exons with N*(upfi+dofi)* ≥ 10 and for exons with multiple start and/or end coordinates, we chose the version with the longest length. In addition, for those datasets with matched polyA/cytoplasmic RNA-seq (Table S1 and Supplementary File 1), we used the *Insplico* information to discard those exons whose upstream and/or downstream introns had PIR > 0.1.

### ISO and feature comparisons for exons based on their tissue-specific regulation

We generated a barplot of F*upfi* values for mouse exons depending on their splicing pattern (Fig. 6a) as determined from the cytoplasmic polyA-selected brain and liver data that we generated for this study (Table S1). In particular, we defined the following groups: (i) constitutively spliced (CS): exons with PSI > 0.99 in both tissues; (ii) tissue-regulated (TR): exons with an absolute difference in PSI between brain and liver higher than 0.25; and (iii) alternatively spliced (AS): exons with 0.1 < PSI < 0.9 in liver and/or brain and that is not TR. Only exons with at least 20 reads contributing to the PSI estimates in both brain and in liver polyA-selected samples and at least N*(upfi+dofi)* ≥ 10 in the brain chromatin-associated RNA-seq sample were used for the analysis. Exons with multiple start and/or end coordinates were discarded. Exon-intron related features were retrieved using *Matt cmpr_exons* and plotted as Z-score values (Fig. 6a). P-values corresponded to Bonferroni-corrected p-values from Wilcoxon Rank-Sum tests with respect to the distribution of the CS exons. Full details of the comparisons and all statistical tests are reported in Supplementary File 2.

### Analysis of SRRM4-dependent microexons

To define *SRRM4*-dependent microexons, we processed with *Insplico* as described above two replicates of total (generated for this study, see below) and matched polyA-selected (from (28) and (40)) RNA-seq data from human HEK293 cells ectopically expressing *GFP* (control) or 3xFlag-tagged human *SRRM4*. The two replicates were pooled together to increase read depth. To obtain the exon-definition table, we clustered Ensembl hg38 v88 annotations and VastDB exons with *extract_exons_from_gtf*.*pl*. We then defined two sets of microexons, defined here as exons of length ≤ 51 nts: (i) *SRRM4*-dependent microexons: exons with a ΔPSI (SRRM4 - control) > 0.15 and a PSI in control cells ≤ 0.2 in the polyA-selected samples; and (ii) control microexons: with |ΔPSI| < 0.02 and 0.05 ≤ PSI ≤ 0.95 in at least one of the samples. For exons with multiple start and/or end coordinates, we chose the version with the shortest length. Only exons with N(*upfi+dofi*) ≥ 5 in the SRRM4 OE total RNA sample were used for ISO analyses. Exons with *upfi* and *dofi* patterns were defined as those with F*upfi* ≥ 0.8 and ≤ 0.2, respectively. Intron-exon related features were extracted using *Matt cmpr_exons* and the full report is shown in Supplementary File 3. The RNA map showing the distribution of UGC motifs was generated using *Matt rna_maps* (27), using a sliding window of 25 nts and limits of 20 and 150 nts for the exonic and intronic regions, respectively.

### Validation of ISO patterns through RT-PCR assays

Flp-in-T-REx 293 cells expressing either GFP or 3xFlag-tagged human SRRM4 were induced with 1 ug/ml for 24 h (28). Total RNA was extracted using the illustra RNAspin Mini Isolation kit (GE Healthcare). Reverse transcription was performed using oligo-dT and random hexamer primers with an *in-house* enzyme produced by the Protein Technologies Unit at CRG. PCRs were performed using GoTag DNA polymerase (Promega) and primers annealing either of the flanking exons to look at the pattern of splicing or exon-junction (EJ) overlapping and intronic (I) primers to investigate the order of intron splicing. Primers (5′ to 3′, Sense and AntiSense) used: *ERC1_EJ1_S*: AGCTGAGTTGGAAAGTCTCACCTC;*ERC1_I2_AS*:TCCCCTCCTCTTTCCTCGTA;*ERC 1_I1_S*:TGTGACTCCTTCCCTTCTCT;*ERC1_EJ2_AS*:TATTCTGGTCTTTCACTTGCCTTGAGGTG;*GRAMD1A_EJ1_S*:TCATCAGCATTGTGATCTGT;*GRAMD1A_I2_AS*:CCCA TTGCAGAGGAGGAGAA;*GRAMD1A_I1_S*:CGTCCTGAGAGAGTGGAGAC;*GRAMD1A _EJ2_AS*:GAGGATGATAAGGCTCACAC;*KIF1B_EJ1_S*:CTTGGCCGAGGTGGATAAC T*;KIF1B_I2_AS*:ACCCACAGACACACAATCCA;*KIF1B_I1_S*:ATGCTGTTGATTTGAG GGCC;*KIF1B_EJ2_AS*:TCTTCTTTTTACTCTTGCTA;*UGGT2_EJ1_S*:TTTCTCTTTGGG AAACTAAAACAAGGAA;*UGGT2_I2_AS*:GAGAACCACCCTGAGAGTCC;*UGGT2_I1_ S*:GCCCCAAAGAAAAGAAAACGT;*UGGT2_EJ2_AS*:TCTAAGATCTGAATATATTTCTCATGCTATTCCTTG. Events were selected among those having ΔPSI (SRRM4-GFP) > 40 and N*(upfi+dofi)* ≥ 3 in both *SRRM4* total and *SRRM4* polyA-selected RNA-seq.

### Tissue Dissociation and Cellular Fractionation

Female mice (6-7 weeks old, B6CBAF1) were injected intraperitoneally with 5 IU of pregnant mare serum gonadotropin (PMSG), followed by intraperitoneal injection of 5 IU of human chorionic gonadotropin (hCG) 47 h after. Females were mated after hCG injection and tissues collected 20 h post hCG injection. Mouse euthanasia was performed by cervical dislocation. All animal related protocols were carried out in accordance to the European Community Council Directive 2010/63/EU and approved by the local Ethics Committee for Animal Experiments (Comitè Ètic d’Experimentació Animal-Parc de Recerca Biomèdica de Barcelona, CEEA-PRBB, CEEA number 9086).

The tissues (liver, cerebellum and cortex) were collected post-mortem in cold PBS and rinsed to remove the excess of blood. The tissues were sliced into small pieces using a blade and resuspended in 40 ml of dissociation buffer (trypsin 0.05% (ThermoFisher); 0.02 units/ml dispase (Life Technologies); 0.025 mg/ml collagenase (Life Technologies); 18 units/ml DNAse I (Sigma)). Samples were incubated at 4 ºC head-over-tail overnight before being filtered through a 100 micron strainer (BD Biosciences) to remove the undissociated tissues. Cells pellets were obtained by centrifugation at 1000 rpm for 5 min at 4 ºC and washed once in 1 ml PBS. Pellets were resuspended in pre-chilled HMKE buffer (20 mM Hepes pH 7.2; 5 mM MgCl2; 10mM KCl; 1 mM EDTA; 250 mM sucrose; 1X protease inhibitors cocktail (Roche); 200 ug/ml digitonin (Sigma)) described in (41); supplemented with 0.1% NP40. Samples were incubated for 10 min on ice and centrifuged at 500 g for 10 min at 4 ºC. The supernatant for each tissue was kept and saved as cytoplasmic fraction. The pellet, containing the nuclei, was washed in PBS supplemented with 1 mM DTT, centrifuged again and treated following the protocol described by (42). Briefly, pellets were resuspended in pre-chilled buffer 1 (20 mM Tris-HCl pH 7.9; 75 mM NaCl; 0.5 mM EDTA; 0.85 mM DTT; 1X protease inhibitors cocktail (Roche); 50% glycerol). An equal volume of pre-chilled buffer 2 (10 mM Hepes pH 7.6; 1 mM DTT; 7.5 mM MgCl2; 0.2 mM EDTA; 0.3 M NaCl; 1 M Urea; 1% NP40) was added and the samples were vortexed twice for 2 sec before being incubated for 10 min on ice.

The samples were centrifuged at 15 000 g for 2 min at 4 ºC. The pellet (chromatin fraction) was resuspended in PBS. All fractions were treated with proteinase K for one hour at 65 ºC (an equal volume of Proteinase 2x buffer (200 mM Tris 7.5, 25 mM EDTA, 300 mM NaCl, 2% SDS) was added to each fraction supplemented with 2 mg/ml proteinase K, Roche Diagnostics). A phenol/chloroform extraction was performed followed by a chloroform extraction and ethanol precipitation. The nucleic acid pellets were resuspended in water and treated with DNAse (RQ1, Promega) for one hour at 37 ºC according to the manufacturer’s instructions. Phenol/chloroform and chloroform extractions and ethanol precipitation were performed and the pellets were resuspended in water.

### Preparation of RNA-seq libraries and short read sequencing

Total RNA libraries were prepared using the TruSeq Stranded Total RNA Library Prep Kit with Ribo-Zero Human/Mouse/Rat Kit (REF. RS-122-2201/2202, Illumina) according to the manufacturer’s protocol. Briefly, from 11.7 to 100 ng of total RNA were used for ribosomal RNA depletion. Then, ribosomal depleted RNA was fragmented for 4.5 min at 94 ºC. The remaining steps of the library preparation were followed according to the manufacturer’s instructions. Final libraries were analysed on an Agilent Technologies 2100 Bioanalyzer system using the Agilent DNA 1000 chip to estimate the quantity and validate the size distribution, and were then quantified by qPCR using the KAPA Library Quantification Kit KK4835 (REF. 07960204001, Roche) prior to amplification with Illumina’s cBot. PolyA-selected libraries were prepared using the TruSeq stranded mRNA Library Prep according to the manufacturer’s protocol using from 25 ng to 200 ng of total RNA as starting material.

Both total (cerebellum, cortex, liver and HEK293) and polyA-selected (cerebellum, cortex and liver) RNA libraries were sequenced on an Illumina HiSeq 2500 machine at the CRG Genomics Unit to generate 125 nt paired-end reads. Read numbers and mapping statistics are provided in Table S1, and all samples were submitted to Gene Expression Omnibus (GEO), under the ID GSE207459. For all analyses, cerebellum and cortex RNA-seq reads were pooled together to generate a single “brain” sample.

## ACKNOWLEDGEMENTS

We thank Juan Valcárcel for his support and invaluable feedback throughout the development of the project, Barbara Pernaute and Jon Permanyer for their help to extract the mouse tissues, the CRG Genomics Unit for the RNA sequencing services and the Protein Technologies Unit for enzyme production. The research has been funded by the European Research Council (ERC) under the European Union’s Horizon 2020 research and innovation program (ERC-StG-LS2-637591 and ERCCoG-LS2-101002275 to MI and ERCAdvG 670146 to JV), the Spanish Ministry of Economy and Competitiveness (BFU-2017-89308-P to JV and BFU-2017-89201-P to MI) and the ‘Centro de Excelencia Severo Ochoa 2013-2017’(SEV-2012-0208).

## SUPPLEMENTARY MATERIALS

**Supplementary File 1 -** Plots showing the associations between ISO patterns and all features as obtained by *Matt cmpr_features*. Further methodological details as well as exon sample sizes are included.

**Supplementary File 2 -** Report from *Matt cmpr_exons* for constitutive (CS), alternative (AS) and tissue-regulated (TR) exons, CS *upfi* and CS *dofi* exons.

**Supplementary File 3 -** Report from *Matt cmpr_exons* for control and SRRM4-dependent *upfi* and *dofi* microexons.

**Table S1 - RNA-seq samples used in this study**.

