## Supplementary File 1 for "*Insplico*: Effective computational tool for studying intron splicing order genome-wide with short and long RNA-seq reads"

### 1 Which exon related features affect intron removal order?

The plots show avg(Fupfi) for up to 5 groups of exons defined by increasing feature values. E.g.:

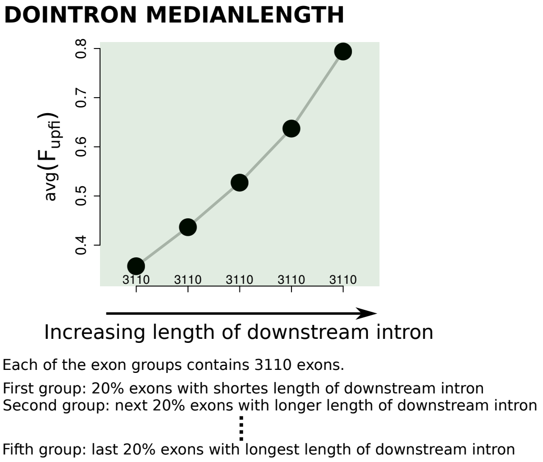


All RNA-Seq data sets come from the nuclear or chromatin RNA fraction. Exons considered here are those for which we have sufficient read support ($N_{\text{upfidofi}} > 9$, exception Chen: $N_{\text{upfidofi}} > 4$) in Insplico analyses. For data sets Jia, Ke, Tilgner, Bonnal, McCorcindale we had in addition poly-A selected RNA-seq data from the cytoplasm. For these data sets, the set of considered exons was further reduced to those with both flanking introns with PIR<10%. Green color indicates a p value $\leq$ 5% applying the Kruskal test across all exon groups, red color indicates a p value > 5%. Features highlighted in red seem most consistent across all data sets of this study.

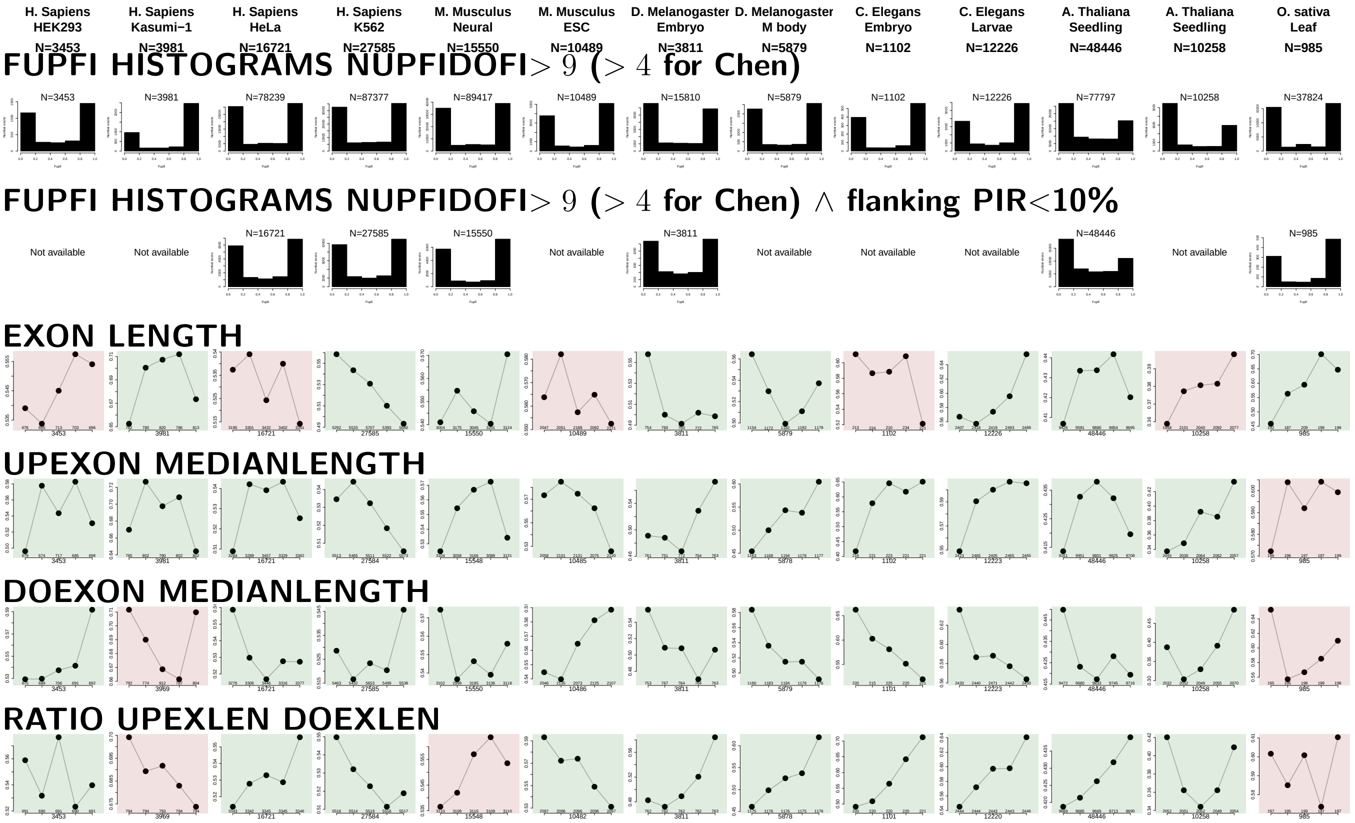


RATIO UPEXON EXON LENGTH

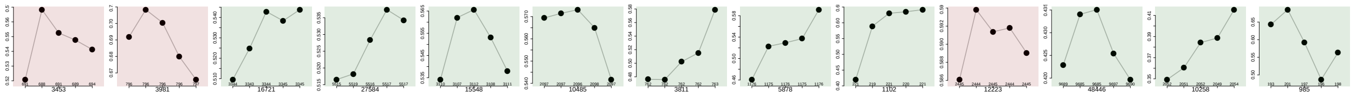

RATIO DOEXON EXON LENGTH

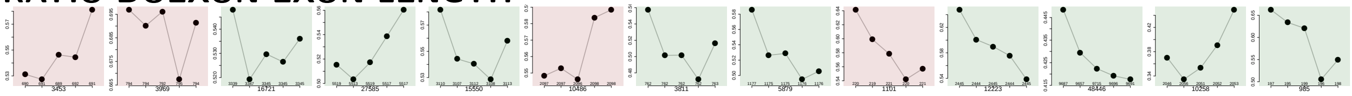

UPINTRON MEDIANLENGTH

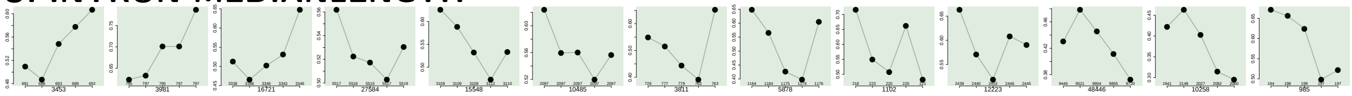

DOINTRON MEDIANLENGTH

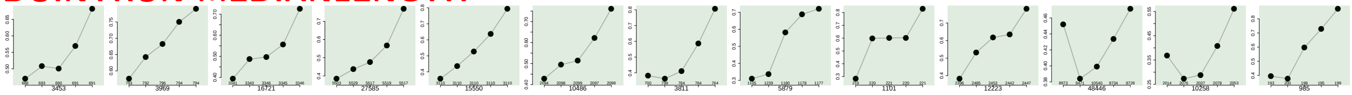

RATIO UPINTRON EXON LENGTH

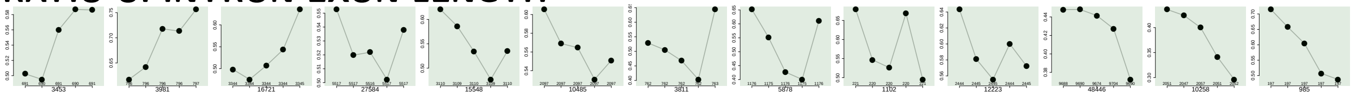

RATIO DOINTRON EXON LENGTH

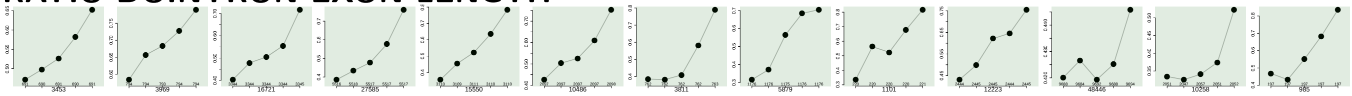

RATIO UPINTLEN DOINTLEN

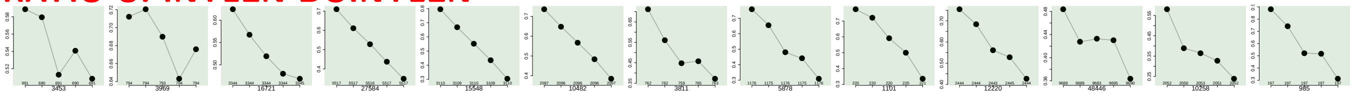

EXON GCC

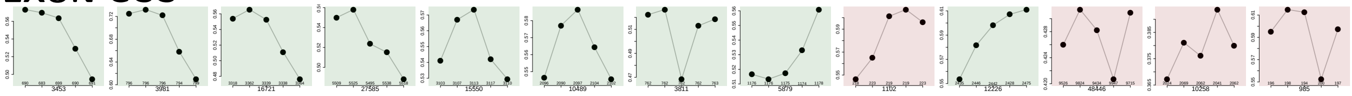

UPINTRON GCC

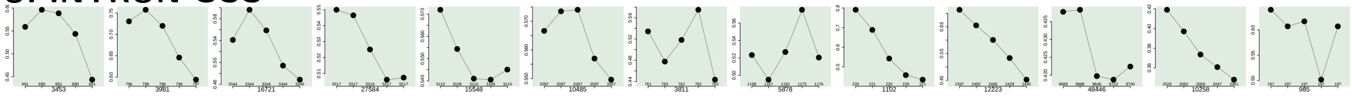

UPEXON GCC

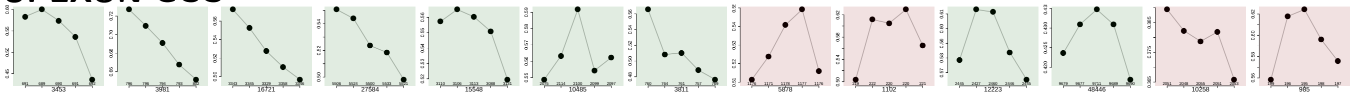

DOINTRON GCC

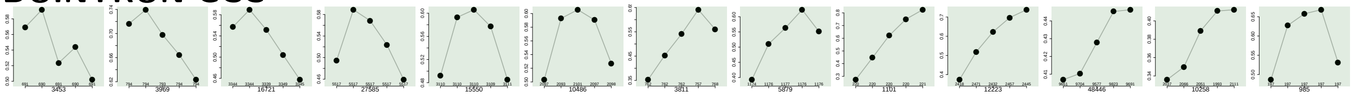

DOEXON GCC

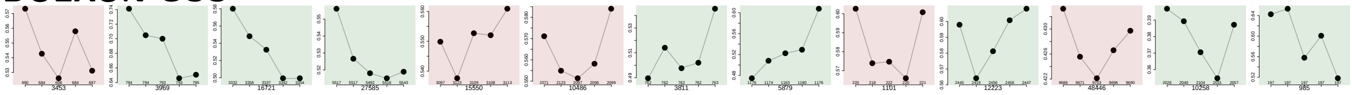

RATIO UPEXON EXON GCC

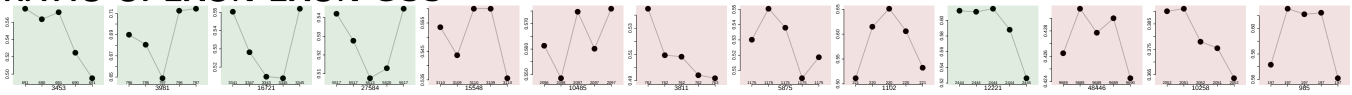

RATIO UPINTRON EXON GCC

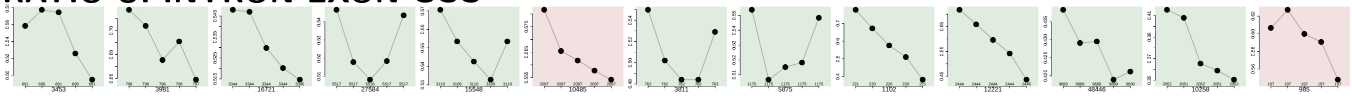

RATIO DOINTRON EXON GCC

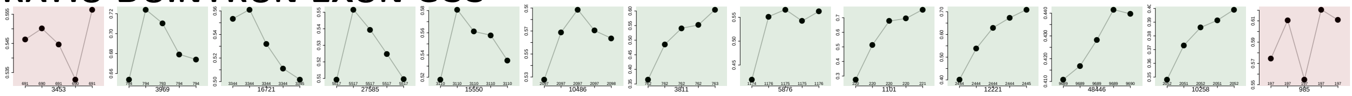

RATIO DOEXON EXON GCC

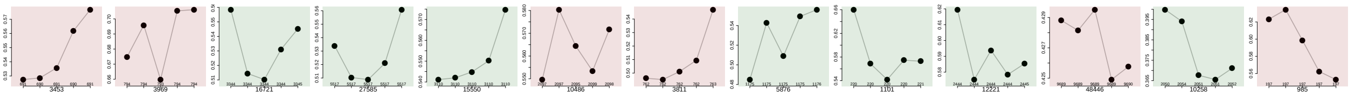

**SF1 HIGHESTSCORE 3SS UPINTRON**

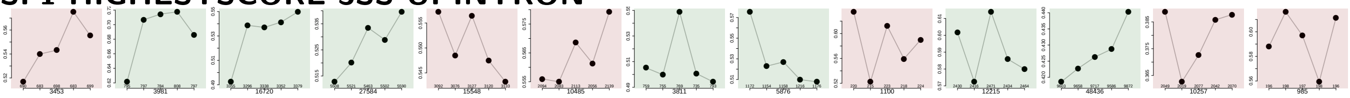

**SF1 HIGHESTSCORE 3SS DOINTRON**

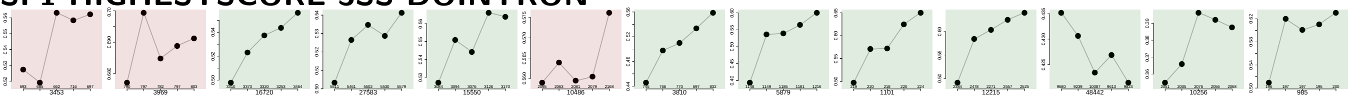

**UP 5SS 20INT10EX GCC**

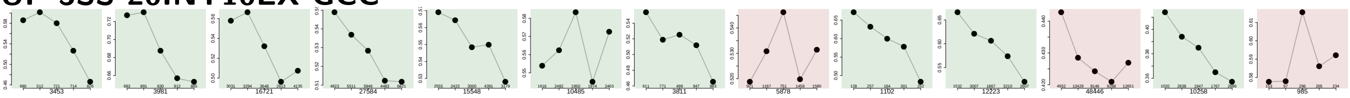

**GCC 3SS 20INT10EX**

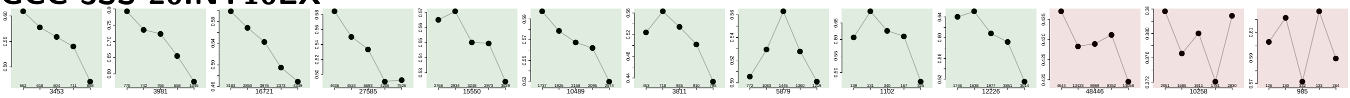

**GCC 5SS 20INT10EX**

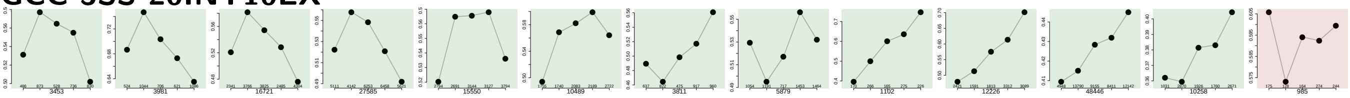

**DO 3SS 20INT10EX GCC**

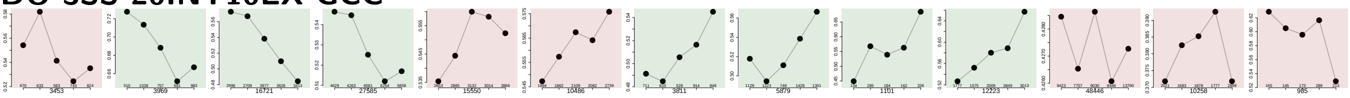

**MAXENTSCR HSAMODEL UPSTRM 5SS**

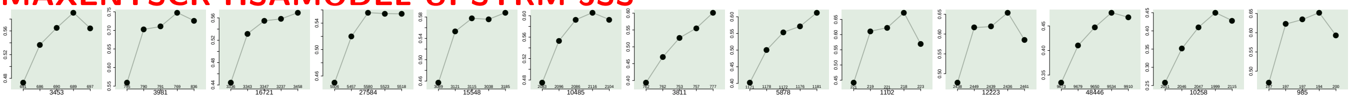

**MAXENTSCR HSAMODEL 3SS**

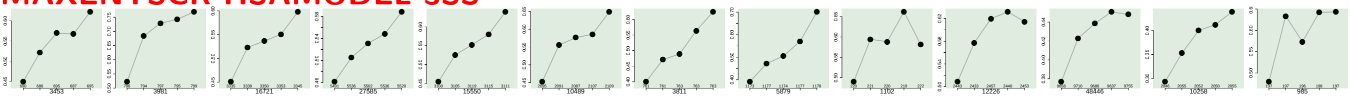

**MAXENTSCR HSAMODEL 5SS**

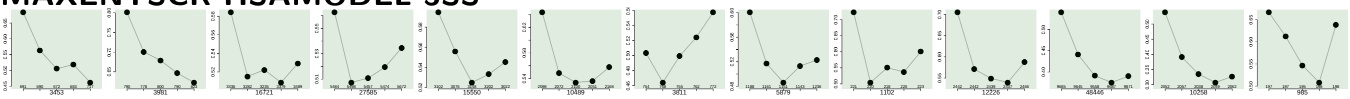

**MAXENTSCR HSAMODEL DOWNSTRM 3SS**

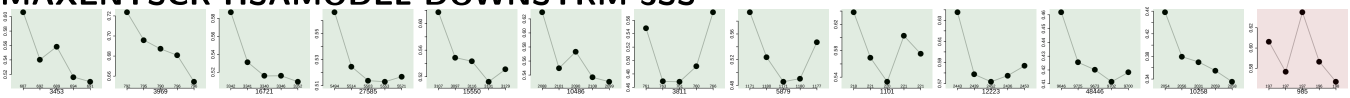

**DIST FROM MAXBP TO 3SS UPINTRON**

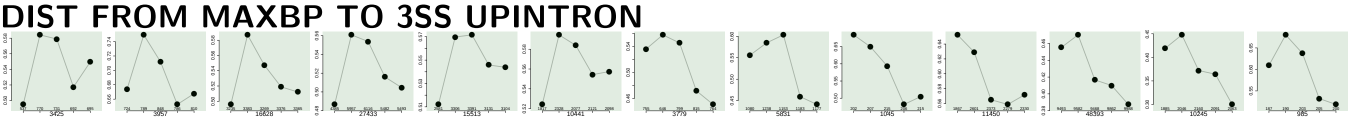

**SCORE FOR MAXBP SEQ UPINTRON**

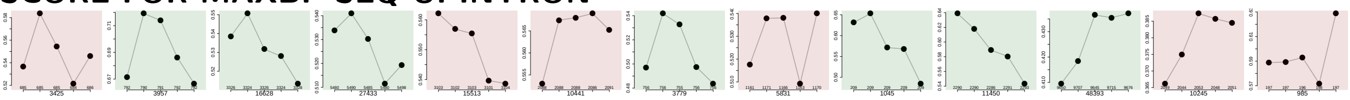

**PYRIMIDINECONT MAXBP UPINTRON**

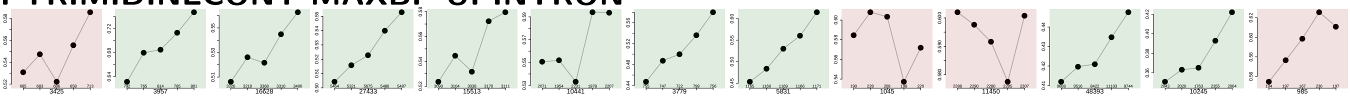

**POLYPYRITRAC OFFSET MAXBP UPINTRON**

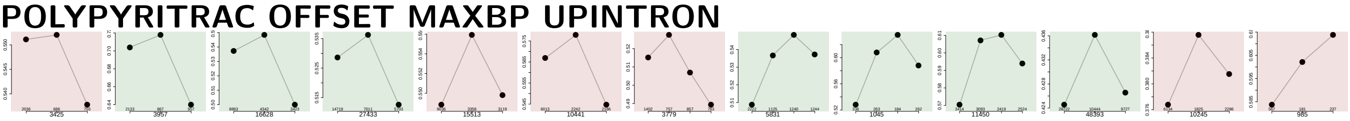

**POLYPYRITRAC LEN MAXBP UPINTRON**

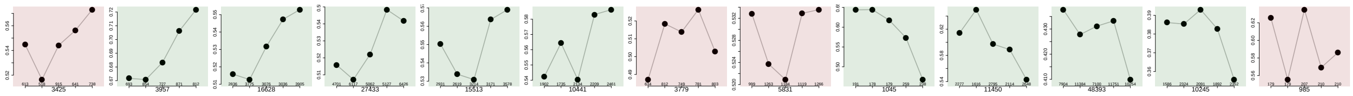

POLYPYRITRAC SCORE MAXBP UPINTRON

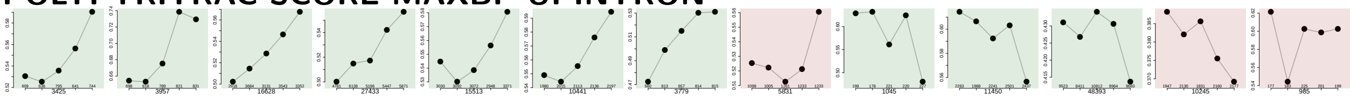

BPSCORE MAXBP UPINTRON

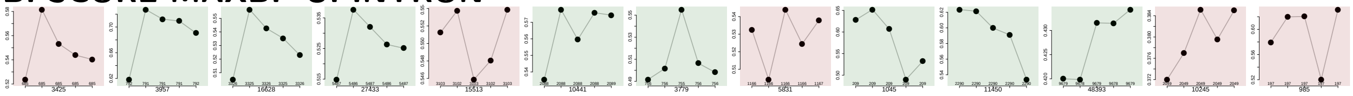

NUM PREDICTED BPS UPINTRON

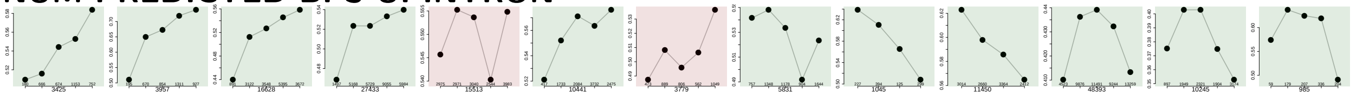

MEDIAN DIST FROM BP TO 3SS UPINTRON

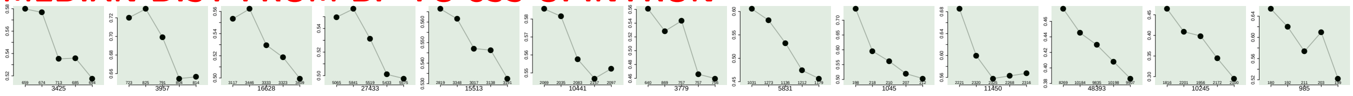

MEDIAN SCORE FOR BPSEQ UPINTRON

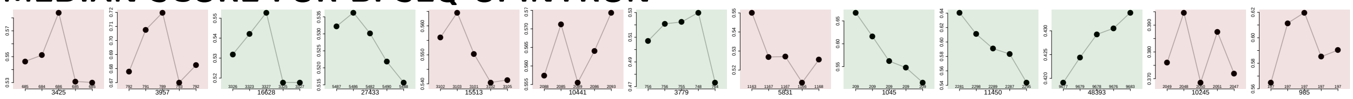

MEDIAN PYRIMIDINECONT UPINTRON

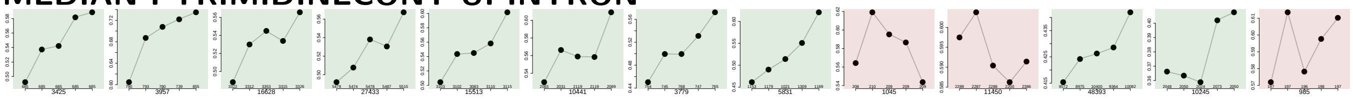

MEDIAN POLYPYRITRAC OFFSET UPINTRON

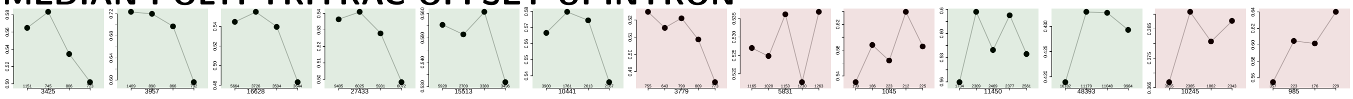

MEDIAN POLYPYRITRAC LEN UPINTRON

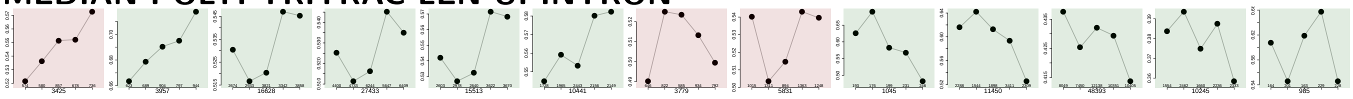

MEDIAN POLYPYRITRAC SCORE UPINTRON

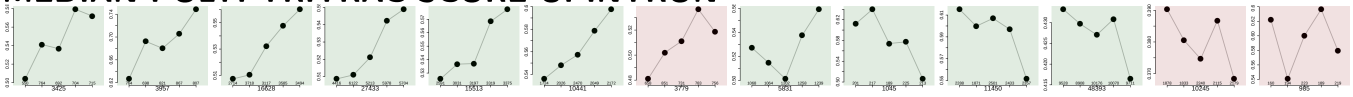

MEDIAN BPSCORE UPINTRON

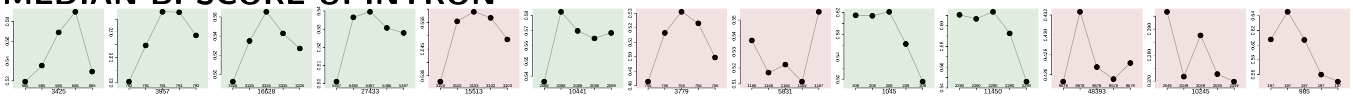

DIST FROM MAXBP TO 3SS DOINTRON

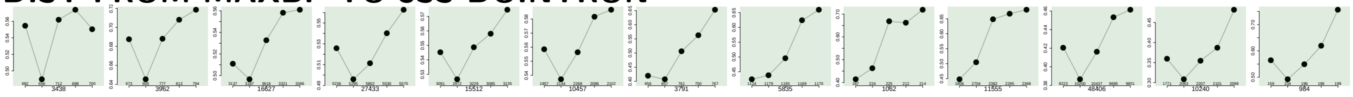

SCORE FOR MAXBP SEQ DOINTRON

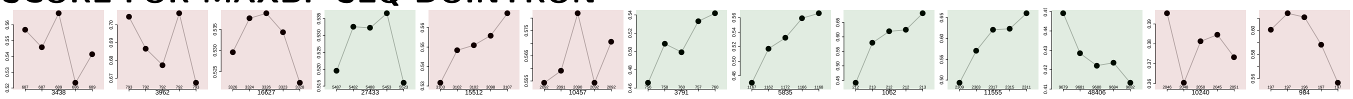

PYRIMIDINECONT MAXBP DOINTRON

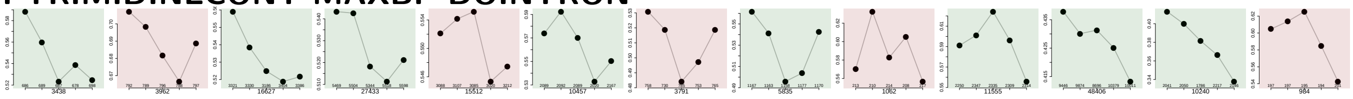

POLYPYRITRAC OFFSET MAXBP DOINTRON

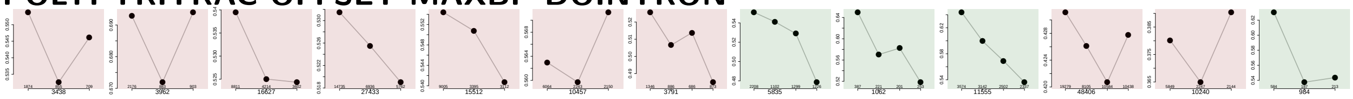

POLYPYRITRAC LEN MAXBP DOINTRON

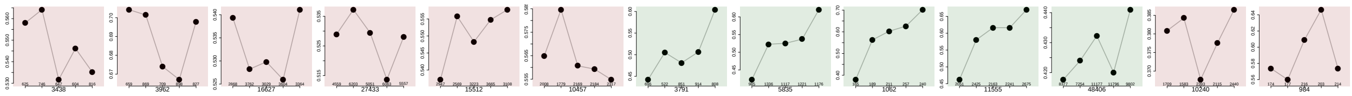

POLYPYRITRAC SCORE MAXBP DOINTRON

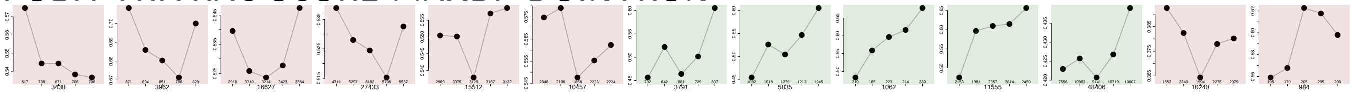

BPSCORE MAXBP DOINTRON

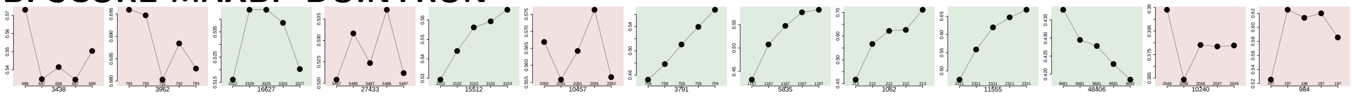

NUM PREDICTED BPS DOINTRON

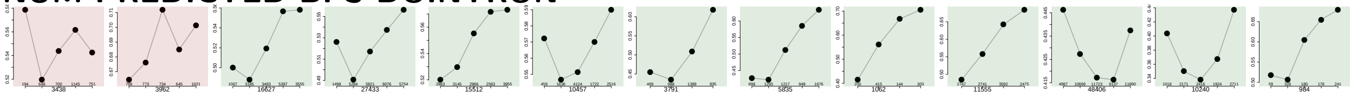

MEDIAN DIST FROM BP TO 3SS DOINTRON

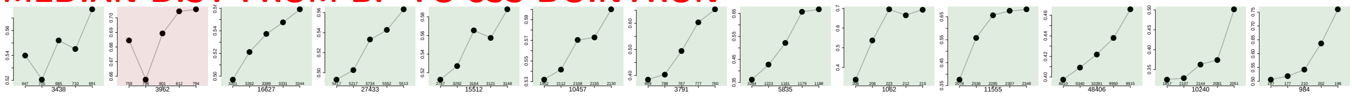

MEDIAN SCORE FOR BPSEQ DOINTRON

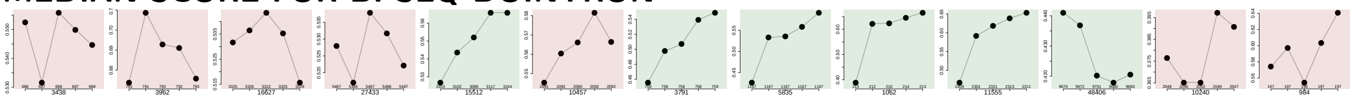

MEDIAN PYRIMIDINECONT DOINTRON

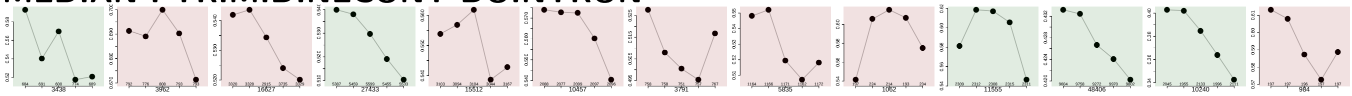

MEDIAN POLYPYRITRAC OFFSET DOINTRON

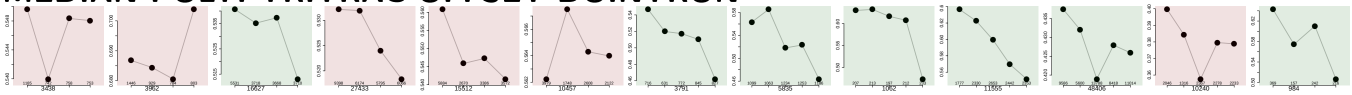

MEDIAN POLYPYRITRAC LEN DOINTRON

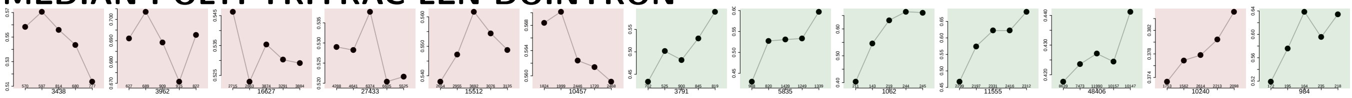

MEDIAN POLYPYRITRAC SCORE DOINTRON

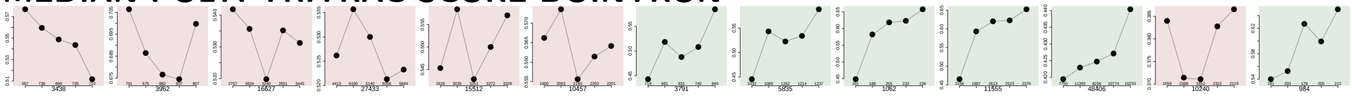

MEDIAN BPSCORE DOINTRON

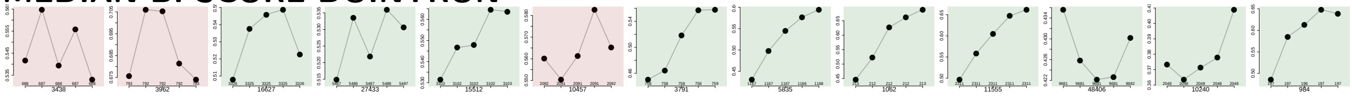

MEDIAN TR LENGTH

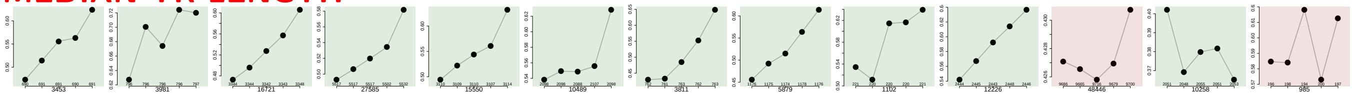

MEDIAN EXON NUMBER

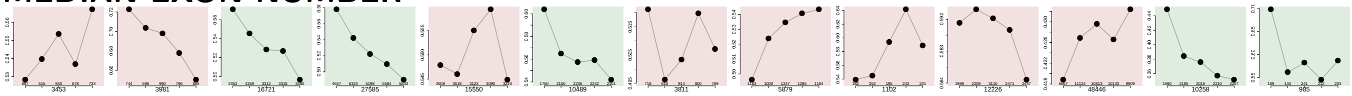

EXON MEDIANRANK

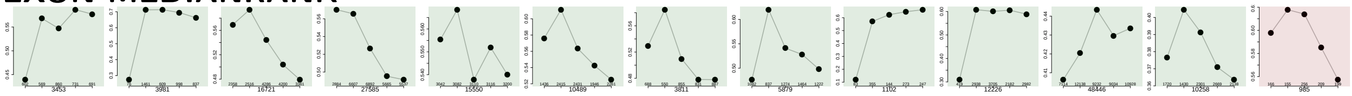

EXON MEDIANRELATIVERANK

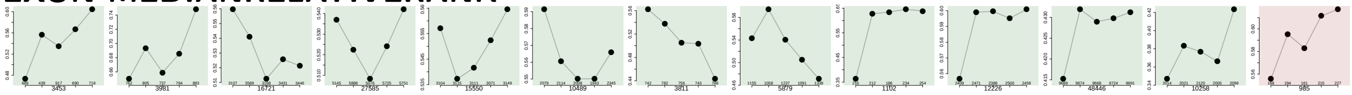

EXON MEDIANRELATIVERANK 3BINS

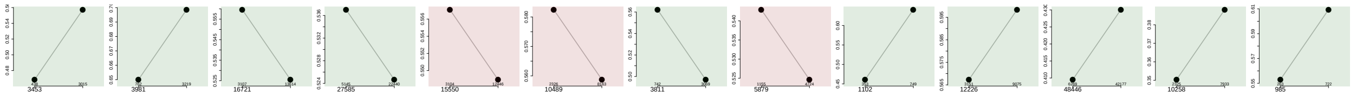

### EXON MEDIANRELATIVERANK 5BINS

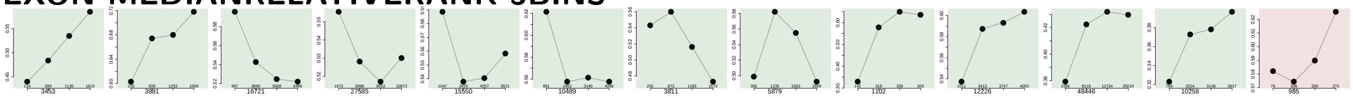

### EXON MEDIANRELATIVERANK 10BINS

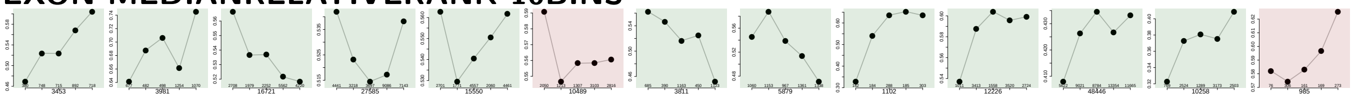

### PROP FIRST EXON

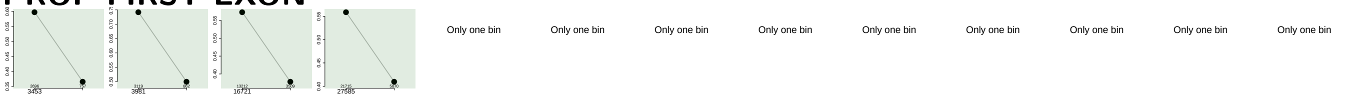


### PROP LAST EXON

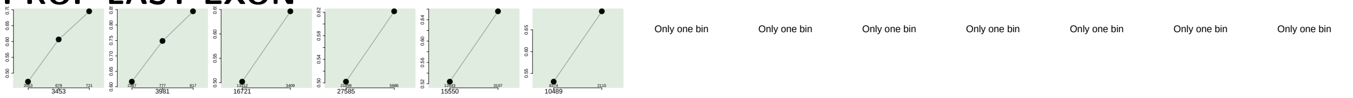


### PROP INTERNAL EXON

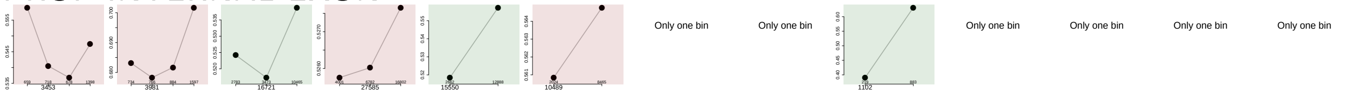


### PROP EXON IN UTR

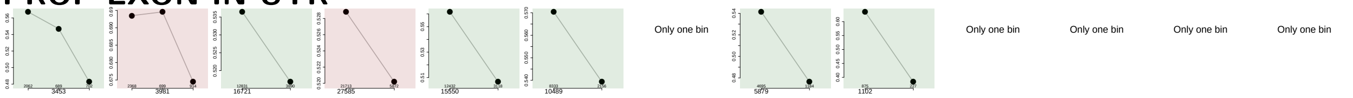


### EXON COOCCURS WITH OTHER EXONS

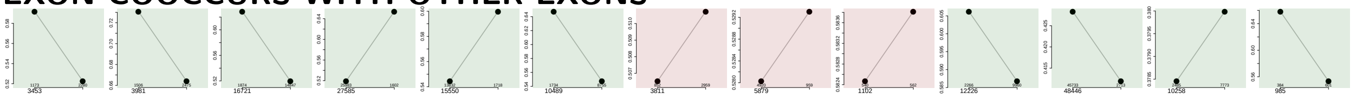

### MEDIAN EXON EXON COOCCURRENCE NUMBER OVER ALL TRS

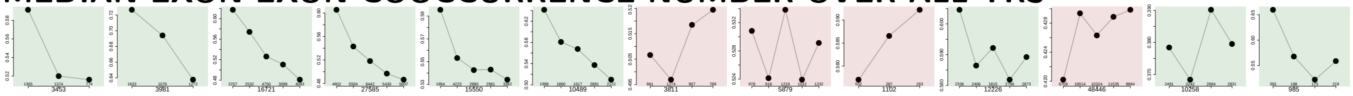
