## Supplementary File 2 for "*Insplico*: Effective computational tool for studying intron splicing order genome-wide with short and long RNA-seq reads"

### Comparison of exons grouped into: CS-Upfi, CS-Dofi, CS, AS, TS

June 6, 2022
Matt version 1.3.0

#### Contents

**1 Infos** 4

**2 Warning: Please read this note carefully** 4

**3 Notes for publishing results** 4

**4 Data sets** 5

**5 Overview: Features with statistically significant differences (p-val $\leq 0.05$)** 6

**6 Details: Box plots and statistical assessments for all features** 19

#### 1 Infos

Visualizations of exon features for different groups of exons. Each exon occurs in exactly one gene, but might occur in several transcripts of that gene. Hence, for some features like the exon length, there is exactly one value for each exon. For other features, e.g., length of the up-stream exon(s), which could be different in different transcripts, there might be several values for each exon. Consequently, in the latter cases, the median of these value gets reported.

#### 2 Warning: Please read this note carefully

Please keep in mind that some features might affect other features. Especially: all branch-point features get extracted from sub-sequences of introns, by standard the last 150 nt at the 3' end of each intron (if you haven't changed this) always neglecting the first 20 nt at their 5' end. If introns of one set are especially short, i.e., many are shorter then these 150 nt, then the shorter intron length might affect branch-point features. For example, there might be less branch points found in shorter introns or their distance to the 3' intron ends might be generally shorter simply because of their shorter intron length.

#### 3 Notes for publishing results

The Matt paper: *Matt: Unix tools for alternative splicing analysis, A. Gohr, M. Irimia, Bioinformatics, 2018, bty606, DOI: 10.1093/bioinformatics/bty606*
When publishing results wrt. splice site strengths which you determined for your data using matt, please cite: *Maximum entropy modeling of short sequence motifs with applications to RNA splicing signals, Yeo et al., 2003, DOI: 10.1089/1066527041410418*
When publishing results wrt. branch point features which you determined for your data with matt, please cite: *Genome-wide association between branch point properties and alternative splicing, Corvelo et al., 2010, DOI: 10.1371/journal.pcbi.1001016*
When publishing results with respect to the binding strength of the human Sf1 splicing factor, you might refert to where the Sf1 binding motif comes from: *Analysis of in situ pre-mRNA targets of human splicing factor SF1 reveals a function in alternative splicing, Margherita Corioni, Nicolas Antih, Goranka Tanackovic, Mihaela Zavolan, and Angela Kramer, 2011, DOI: 10.1093/nar/gkq1042*
The Sf1 binding motif is described in supplement, page 13, table S2: Weight matrix of the binding specificity of SF1.

#### 4 Data sets

Input file:
Exon_types_for_Matt-Sophie_RNAseq-Mar2022-20-10-0.25-bothTR.txt
Selection criteria for defining exons groups:
CS_Upfi : having value CS_Upfi in column GROUP
CS_Dofi : having value CS_Dofi in column GROUP
CS : having value CS in column GROUP
AS : having value AS in column GROUP
TS : having value TS in column GROUP
Exon duplicates removal: yes
Numbers of exons per group before / after neglecting exons which were not found in GTF file (gene annotation). For the comparisons only exons which were found in the gene annotation are used. These numbers might change slightly for each feature if NAs occur.
CS_Upfi: 3372 / 3362
CS_Dofi: 2710 / 2706
CS: 7367 / 7352
AS: 350 / 348
TS: 111 / 110

### 5 Overview: Features with statistically significant differences (p-val $\leq 0.05$)

#### DOINTRON MEDIANLENGTH

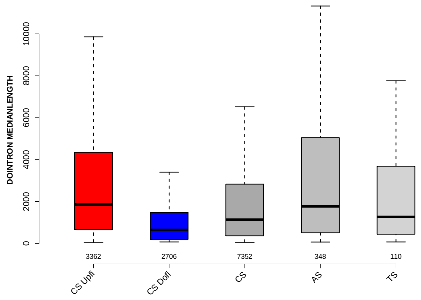

#### RATIO DOINTRON EXON LENGTH

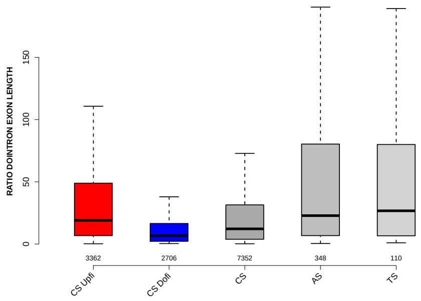

#### PROP FIRST EXON

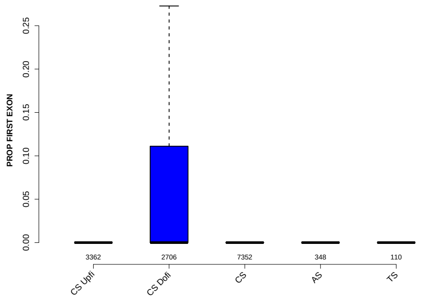

#### PROP LAST EXON

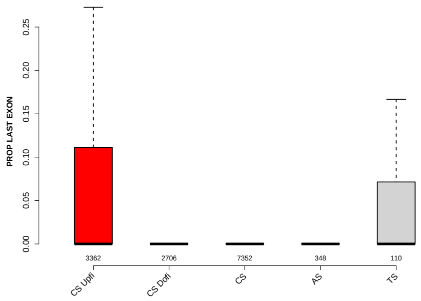

### MAXENTSCR HSAMODEL 3SS

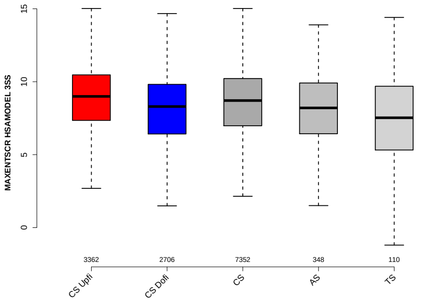

### MAXENTSCR HSAMODEL UPSTRM 5SS

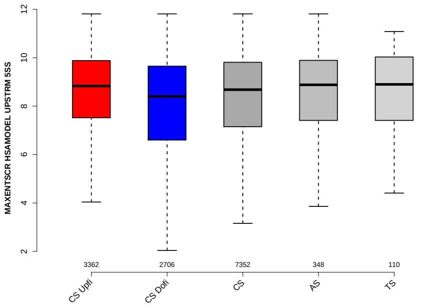

### MEDIAN TR LENGTH

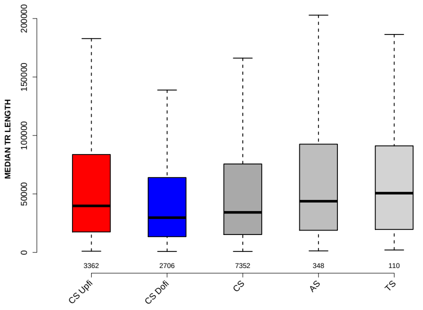

### UPINTRON MEDIANLENGTH

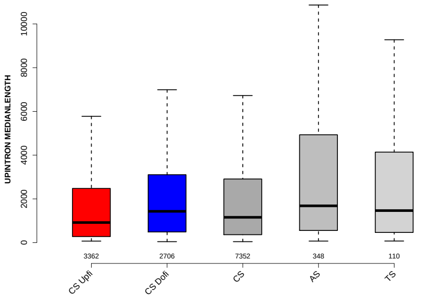

### RATIO UPINTRON EXON LENGTH

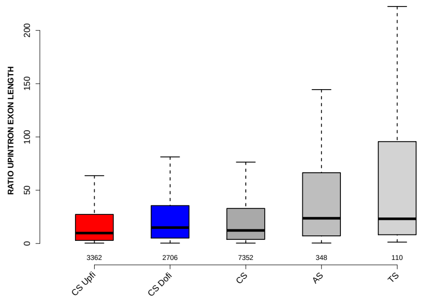

### EXON LENGTH

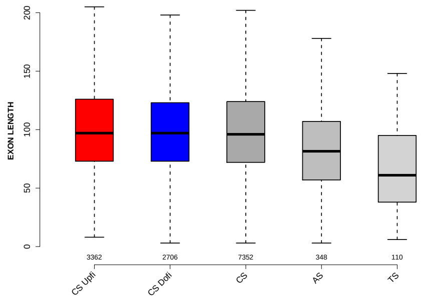

### MAXENTSCR HSAMODEL DOWNSTRM 3SS

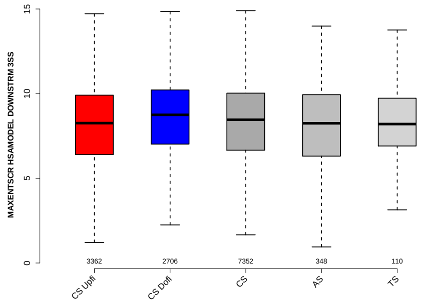

### MEDIAN PYRIMIDINECONT UPINTRON

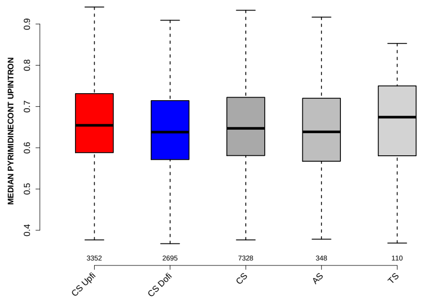

### MEDIAN DIST FROM BP TO 3SS DOINTRON

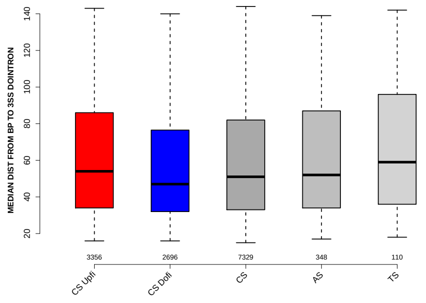

### MAXENTSCR HSAMODEL 5SS

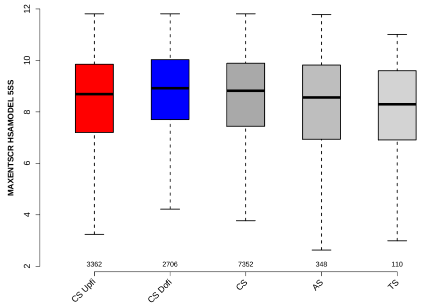

### RATIO UPEXON EXON LENGTH

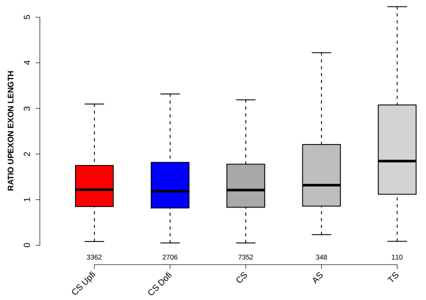

### PYRIMIDINECONT MAXBP UPINTRON

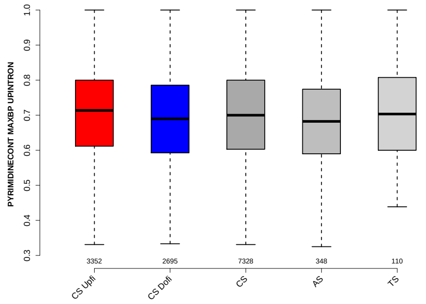

### RATIO DOEXON EXON LENGTH

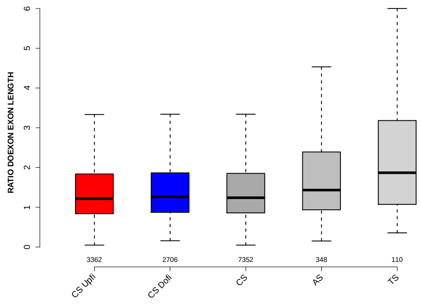

### PROP EXON IN UTR

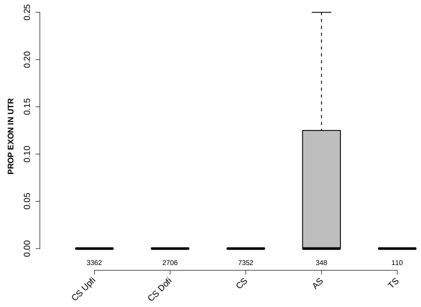

### GCC 3SS 20INT10EX

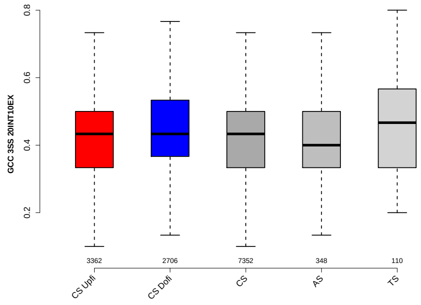

### NUM PREDICTED BPS DOINTRON

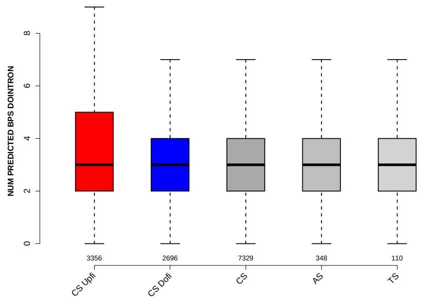

#### POLYPYRITRAC SCORE MAXBP UPINTRON

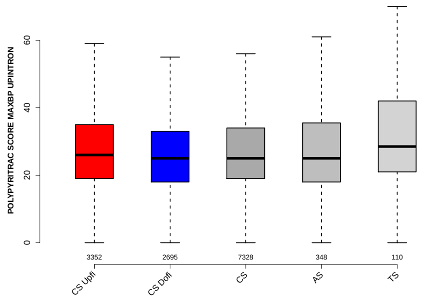

#### MEDIAN POLYPYRITRAC SCORE UPINTRON

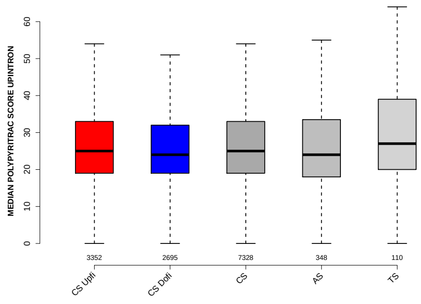

#### SF1 HIGHESTSCORE 3SS DOINTRON

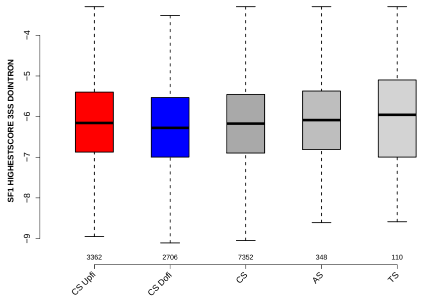

#### RATIO DOEXON EXON GCC

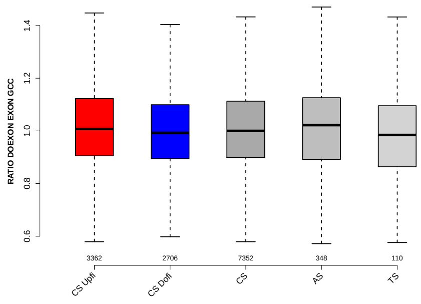

### MEDIAN POLYPYRITRAC LEN UPINTRON

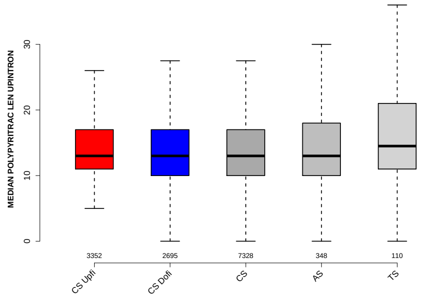

### EXON GCC

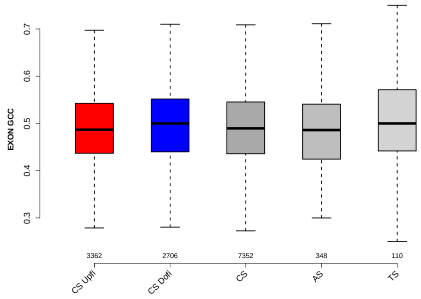

### DOINTRON GCC

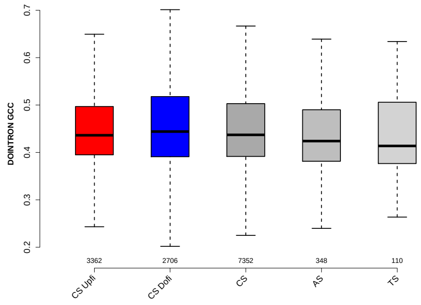

### MEDIAN DIST FROM BP TO 3SS UPINTRON

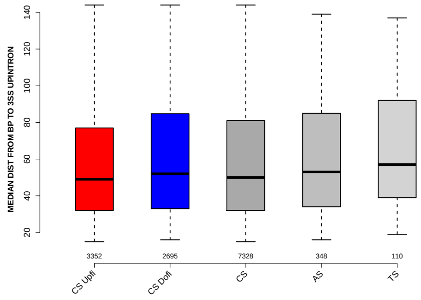

### POLYPYRITRAC LEN MAXBP UPINTRON

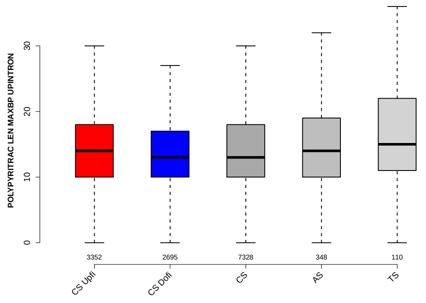

### NTRS ALL FOR GENE

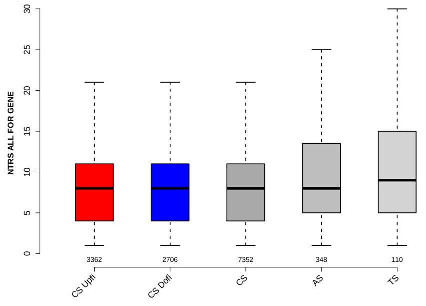

### UPINTRON GCC

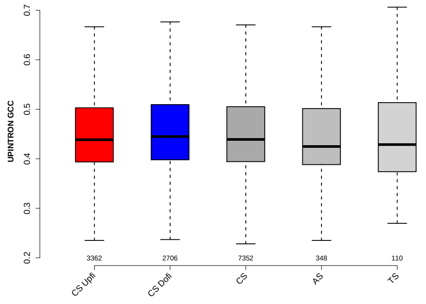

### DIST FROM MAXBP TO 3SS UPINTRON

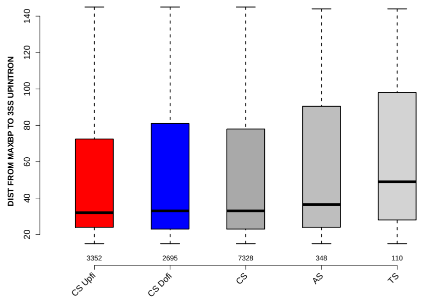

#### UP 5SS 20INT10EX GCC

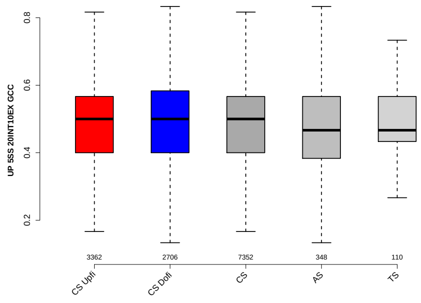

#### MEDIAN EXON NUMBER

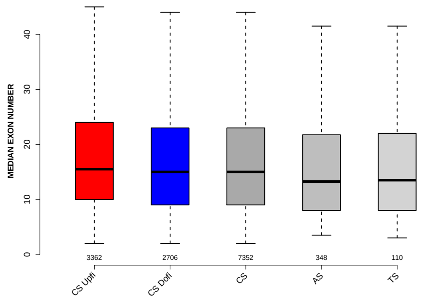

#### DIST FROM MAXBP TO 3SS DOINTRON

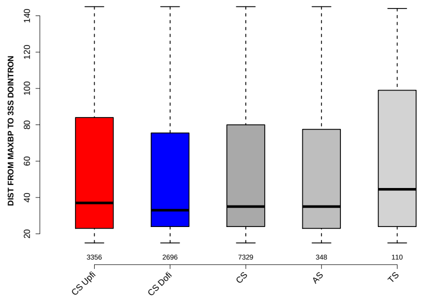

#### RATIO DOINTRON EXON GCC

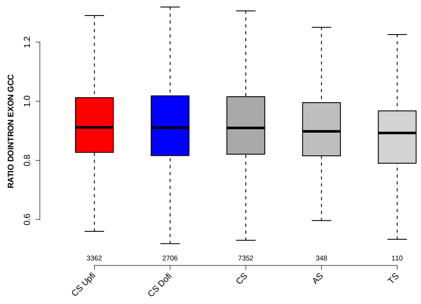

### MEDIAN PYRIMIDINECONT DOINTRON

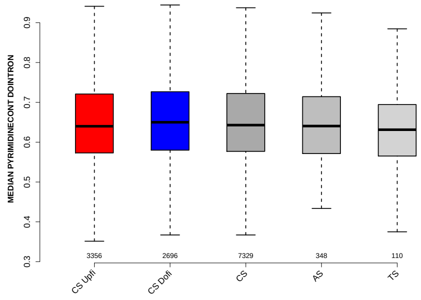

### UPEXON GCC

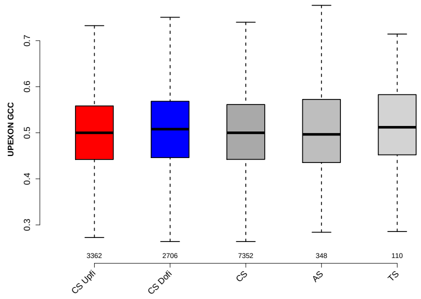

### PROP INTERNAL EXON

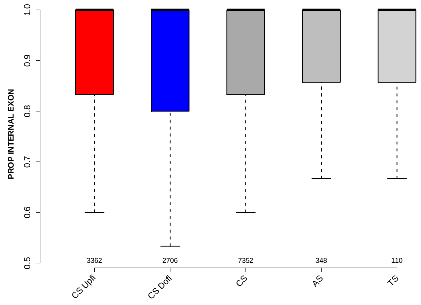

### EXON MEDIANRELATIVERANK 5BINS

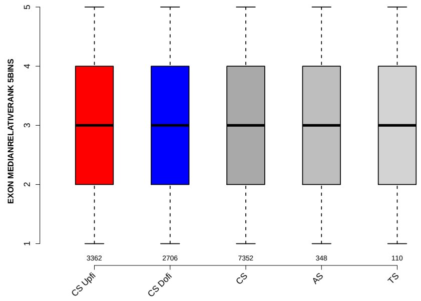

### EXON MEDIANRELATIVERANK

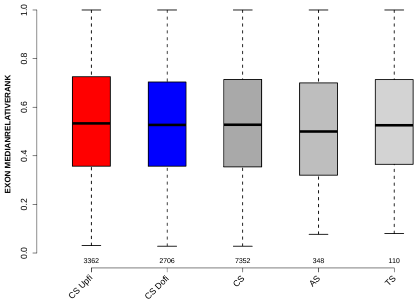

### SF1 HIGHESTSCORE 3SS UPINTRON

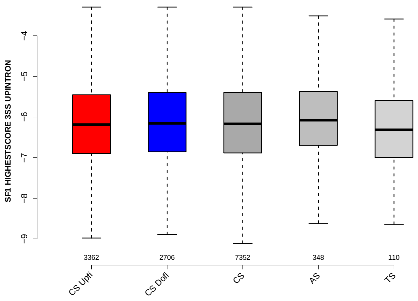

### MEDIAN POLYPYRITRAC OFFSET UPINTRON

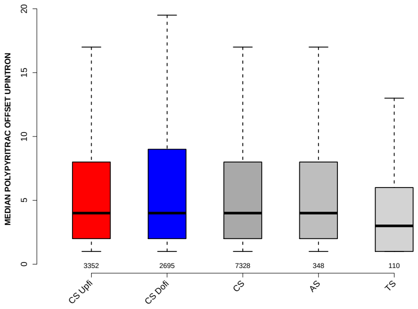

### EXON MEDIANRELATIVERANK 3BINS

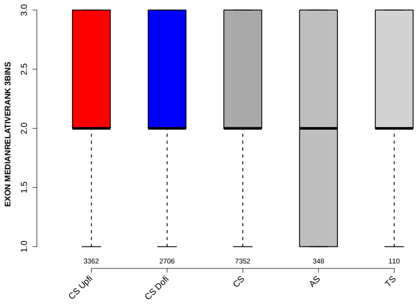

#### UPEXON MEDIANLENGTH

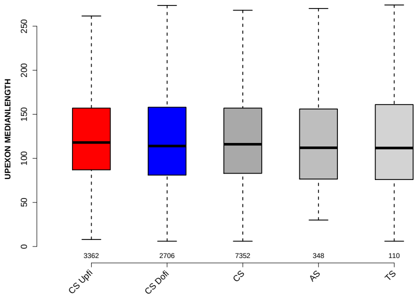

#### PYRIMIDINECONT MAXBP DOINTRON

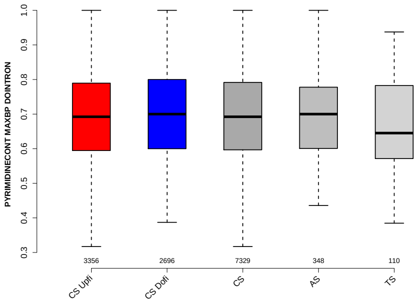

#### DO 3SS 20INT10EX GCC

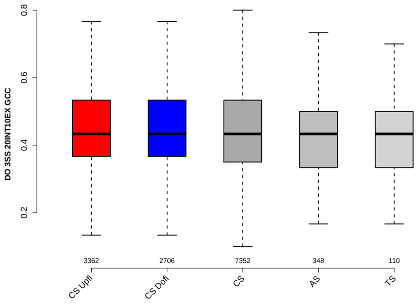

#### EXON MEDIANRELATIVERANK 10BINS

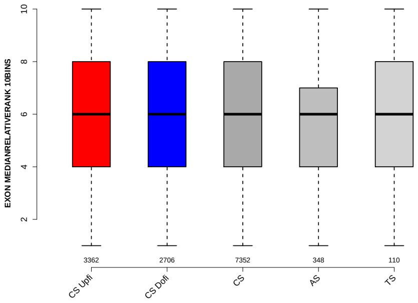

### SCORE FOR MAXBP SEQ UPINTRON

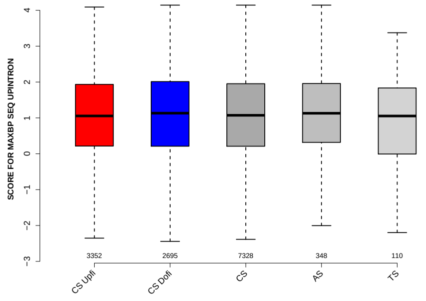

### RATIO UPINTRON EXON GCC

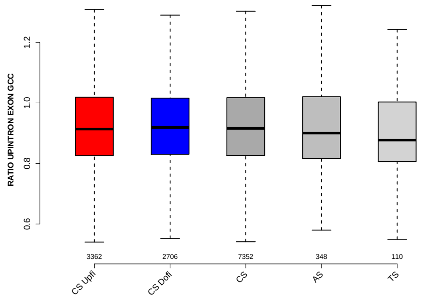

### NUM PREDICTED BPS UPINTRON

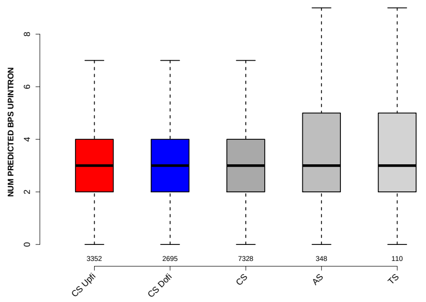

### GCC 5SS 20INT10EX

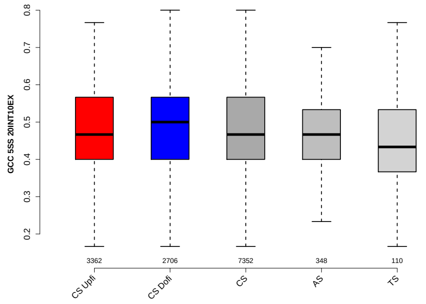

### 6 Details: Box plots and statistical assessments for all features

#### 6.1 EXON LENGTH

Back to: Overview | ToC

Meaning:

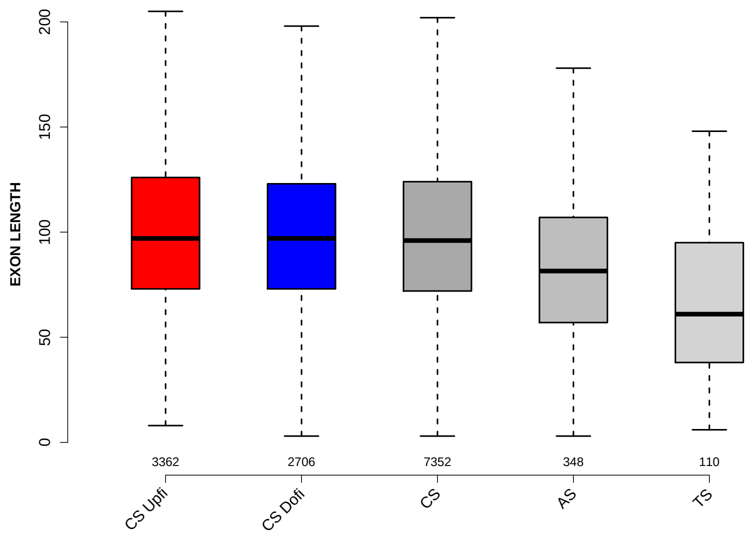

Significant results from Mann-Whitney U test:

- CS_Upfi vs CS : 0.0336456
  mean: 101.8412 > 99.773 , median: 97 > 96
- CS_Upfi vs AS : 1.98702e-13
  mean: 101.8412 > 85.9856 , median: 97 > 81.5
- CS_Upfi vs TS : 1.10529e-16
  mean: 101.8412 > 68.6636 , median: 97 > 61
- CS_Dofi vs AS : 3.15866e-11
  mean: 99.4926 > 85.9856 , median: 97 > 81.5
- CS_Dofi vs TS : 1.18259e-15
  mean: 99.4926 > 68.6636 , median: 97 > 61
- CS vs AS : 6.9502e-12
  mean: 99.773 > 85.9856 , median: 96 > 81.5
- CS vs TS : 7.03164e-16
  mean: 99.773 > 68.6636 , median: 96 > 61
- AS vs TS : 5.35093e-05
  mean: 85.9856 > 68.6636 , median: 81.5 > 61

#### 6.2 UPEXON MEDIANLENGTH

Back to: Overview | ToC

Meaning: median length of up-stream exon

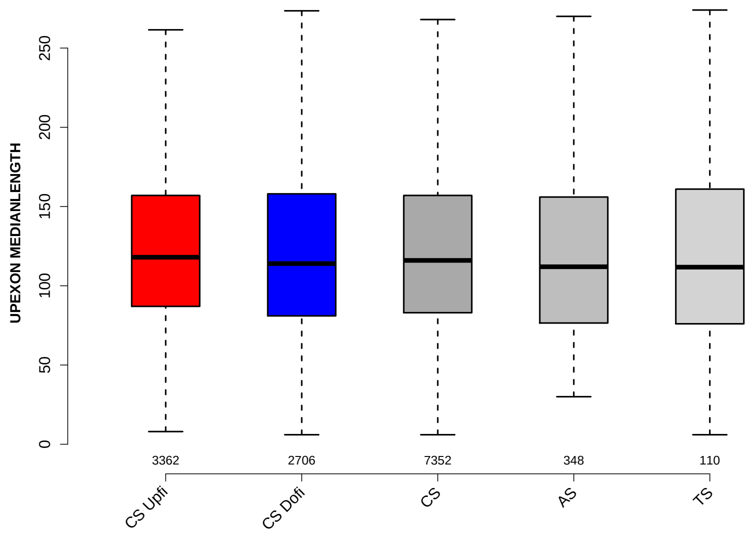

Significant results from Mann-Whitney U test:

- `CS_Upfi` vs `CS_Dofi` : 0.0254725
  mean: 137.5675 < 138.2862 , median: 118 > 114

#### 6.3 DOEXON MEDIANLENGTH

Back to: Overview | ToC

Meaning: median length of down-stream exon

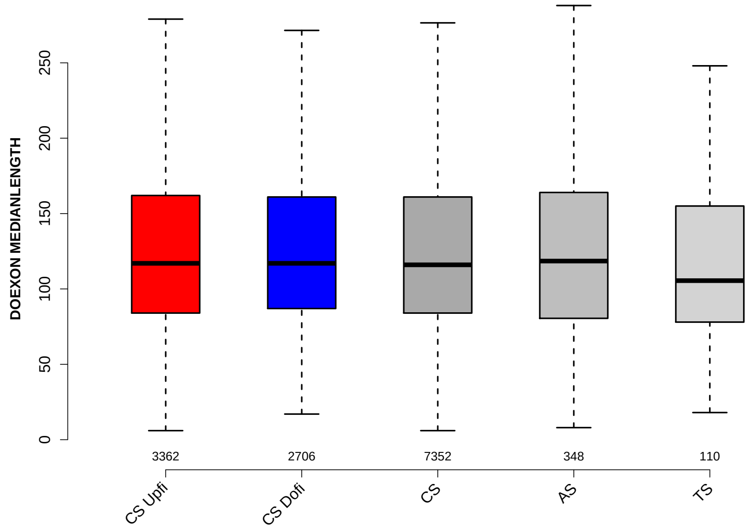

Significant results from Mann-Whitney U test:

- none

#### 6.4 RATIO UPEXON EXON LENGTH

Back to: Overview | ToC

Meaning: median up-stream exon length / exon length

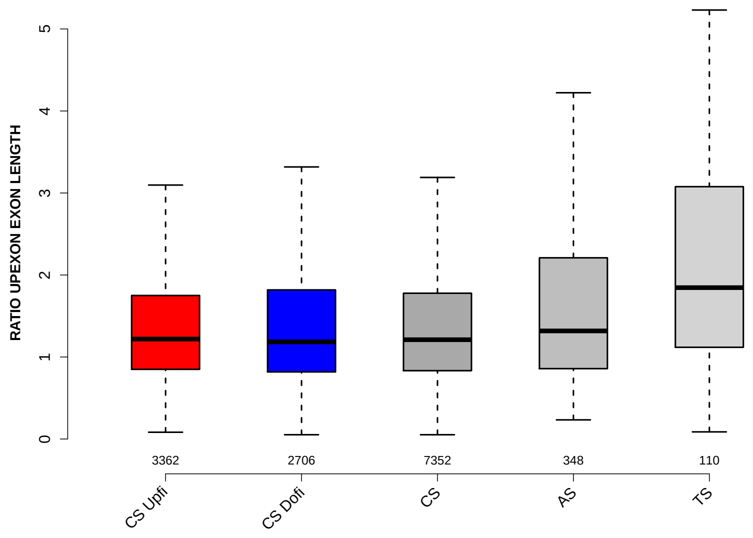

Significant results from Mann-Whitney U test:

- `CS_Upfi` vs `AS` : 0.00150421
  mean: 1.5586 < 2.6204 , median: 1.2187 < 1.3179
- `CS_Upfi` vs `TS` : 1.28201e-08
  mean: 1.5586 < 2.9605 , median: 1.2187 < 1.8458
- `CS_Dofi` vs `AS` : 0.000660032
  mean: 1.6103 < 2.6204 , median: 1.1848 < 1.3179
- `CS_Dofi` vs `TS` : 1.23495e-08
  mean: 1.6103 < 2.9605 , median: 1.1848 < 1.8458
- `CS` vs `AS` : 0.000877341
  mean: 1.6056 < 2.6204 , median: 1.2109 < 1.3179
- `CS` vs `TS` : 1.03331e-08
  mean: 1.6056 < 2.9605 , median: 1.2109 < 1.8458
- `AS` vs `TS` : 0.00154331
  mean: 2.6204 < 2.9605 , median: 1.3179 < 1.8458

#### 6.5 RATIO DOEXON EXON LENGTH

Back to: Overview | ToC

Meaning: median down-stream exon length / exon length

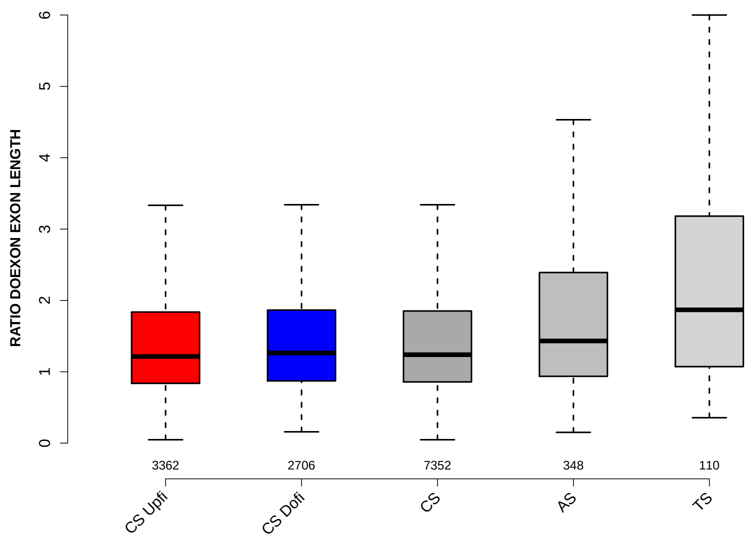

Significant results from Mann-Whitney U test:

- `CS_Upfi` vs `CS_Dofi` : 0.0198101
  mean: 2.1179 < 2.1889 , median: 1.2143 < 1.2632
- `CS_Upfi` vs `AS` : 8.21828e-06
  mean: 2.1179 < 3.0015 , median: 1.2143 < 1.4322
- `CS_Upfi` vs `TS` : 4.97451e-07
  mean: 2.1179 < 3.9427 , median: 1.2143 < 1.8685
- `CS_Dofi` vs `AS` : 0.000417353
  mean: 2.1889 < 3.0015 , median: 1.2632 < 1.4322
- `CS_Dofi` vs `TS` : 4.37454e-06
  mean: 2.1889 < 3.9427 , median: 1.2632 < 1.8685
- `CS` vs `AS` : 5.77337e-05
  mean: 2.1687 < 3.0015 , median: 1.2396 < 1.4322
- `CS` vs `TS` : 1.45613e-06
  mean: 2.1687 < 3.9427 , median: 1.2396 < 1.8685
- `AS` vs `TS` : 0.024612
  mean: 3.0015 < 3.9427 , median: 1.4322 < 1.8685

#### 6.6 UPINTRON MEDIANLENGTH

Back to: Overview | ToC

Meaning: median length of up-stream introns

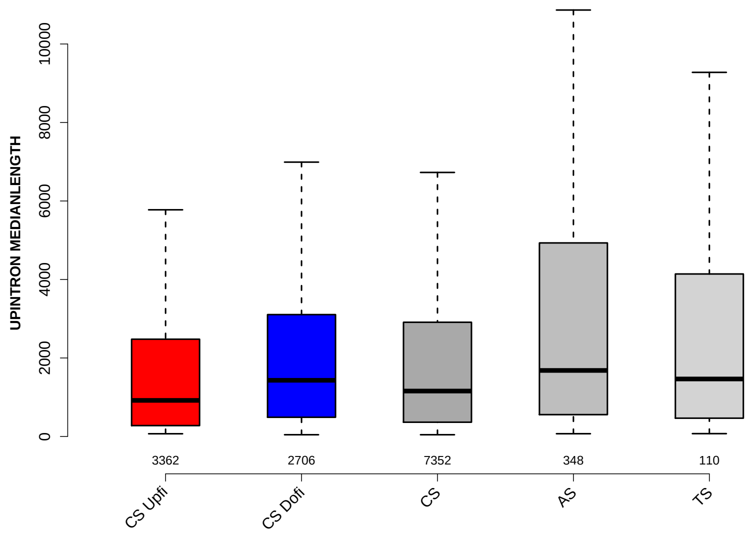

Significant results from Mann-Whitney U test:

- CS_Upfi vs CS_Dofi : 5.51652e-19
  mean: 4481.841 > 2853.2838 , median: 919 < 1431
- CS_Upfi vs CS : 1.34448e-08
  mean: 4481.841 > 3671.7159 , median: 919 < 1157
- CS_Upfi vs AS : 4.77984e-11
  mean: 4481.841 < 5434.1178 , median: 919 < 1682
- CS_Upfi vs TS : 0.00161856
  mean: 4481.841 < 5171.6727 , median: 919 < 1465
- CS_Dofi vs CS : 1.18711e-06
  mean: 2853.2838 < 3671.7159 , median: 1431 > 1157
- CS_Dofi vs AS : 0.00166602
  mean: 2853.2838 < 5434.1178 , median: 1431 < 1682
- CS vs AS : 8.88039e-07
  mean: 3671.7159 < 5434.1178 , median: 1157 < 1682
- CS vs TS : 0.0434662
  mean: 3671.7159 < 5171.6727 , median: 1157 < 1465

#### 6.7 DOINTRON MEDIANLENGTH

Back to: Overview | ToC

Meaning: median length of down-stream introns

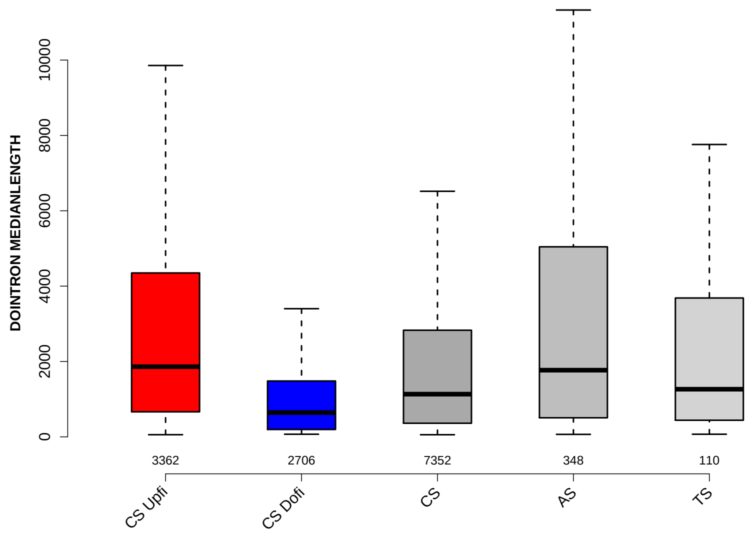

Significant results from Mann-Whitney U test:

- CS_Upfi vs CS_Dofi : 6.32187e-177
  mean: 5957.5623 > 1310.1515 , median: 1862 > 644
- CS_Upfi vs CS : 7.14187e-56
  mean: 5957.5623 > 3621.7709 , median: 1862 > 1133
- CS_Dofi vs CS : 5.12192e-74
  mean: 1310.1515 < 3621.7709 , median: 644 < 1133
- CS_Dofi vs AS : 2.00058e-32
  mean: 1310.1515 < 6487.954 , median: 644 < 1769.5
- CS_Dofi vs TS : 1.57922e-08
  mean: 1310.1515 < 5126.9864 , median: 644 < 1264
- CS vs AS : 9.27271e-08
  mean: 3621.7709 < 6487.954 , median: 1133 < 1769.5

#### 6.8 RATIO UPINTRON EXON LENGTH

Back to: Overview | ToC

Meaning: median up-stream intron length / exon length

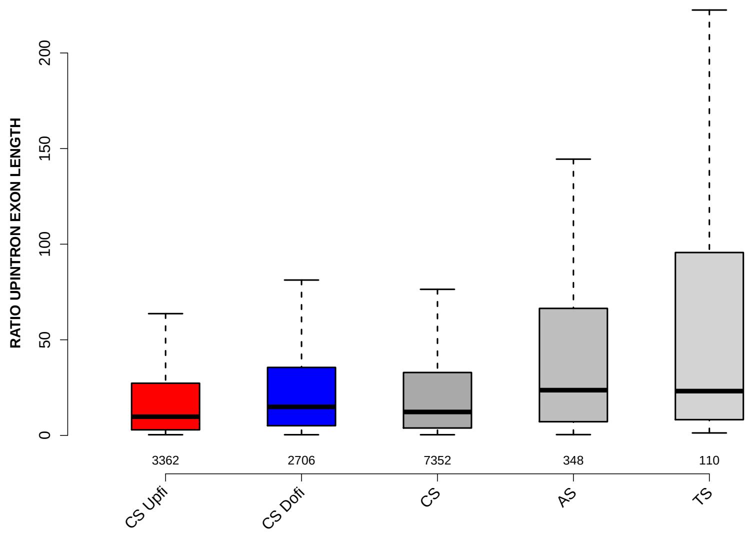

Significant results from Mann-Whitney U test:

- `CS_Upfi` vs `CS_Dofi` : 6.80601e-19
  mean: 49.7789 > 34.4376 , median: 9.7753 < 14.939
- `CS_Upfi` vs `CS` : 3.25567e-09
  mean: 49.7789 > 42.6213 , median: 9.7753 < 12.2652
- `CS_Upfi` vs `AS` : 4.78699e-18
  mean: 49.7789 < 82.8979 , median: 9.7753 < 23.655
- `CS_Upfi` vs `TS` : 6.51836e-09
  mean: 49.7789 < 98.9254 , median: 9.7753 < 23.169
- `CS_Dofi` vs `CS` : 3.94784e-06
  mean: 34.4376 < 42.6213 , median: 14.939 > 12.2652
- `CS_Dofi` vs `AS` : 5.47309e-08
  mean: 34.4376 < 82.8979 , median: 14.939 < 23.655
- `CS_Dofi` vs `TS` : 0.000115735
  mean: 34.4376 < 98.9254 , median: 14.939 < 23.169
- `CS` vs `AS` : 1.49712e-12
  mean: 42.6213 < 82.8979 , median: 12.2652 < 23.655
- `CS` vs `TS` : 1.81264e-06
  mean: 42.6213 < 98.9254 , median: 12.2652 < 23.169

#### 6.9 RATIO DOINTRON EXON LENGTH

Back to: Overview | ToC

Meaning: median down-stream intron length / exon length

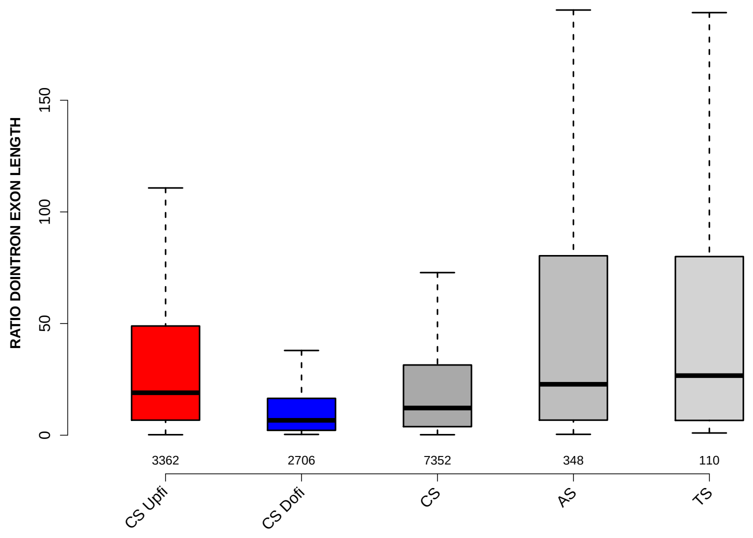

Significant results from Mann-Whitney U test:

- CS_Upfi vs CS_Dofi : 7.11211e-161
  mean: 71.4218 > 18.1512 , median: 19.0114 > 6.6596
- CS_Upfi vs CS : 7.8955e-49
  mean: 71.4218 > 44.4044 , median: 19.0114 > 12.1822
- CS_Upfi vs AS : 0.0443122
  mean: 71.4218 < 101.4713 , median: 19.0114 < 22.8082
- CS_Dofi vs CS : 9.6231e-70
  mean: 18.1512 < 44.4044 , median: 6.6596 < 12.1822
- CS_Dofi vs AS : 8.93068e-42
  mean: 18.1512 < 101.4713 , median: 6.6596 < 22.8082
- CS_Dofi vs TS : 2.01455e-16
  mean: 18.1512 < 105.6229 , median: 6.6596 < 26.6922
- CS vs AS : 2.06688e-13
  mean: 44.4044 < 101.4713 , median: 12.1822 < 22.8082
- CS vs TS : 3.06184e-06
  mean: 44.4044 < 105.6229 , median: 12.1822 < 26.6922

#### 6.10 EXON GCC

Back to: Overview | ToC

Meaning: GC content of entire exon sequence

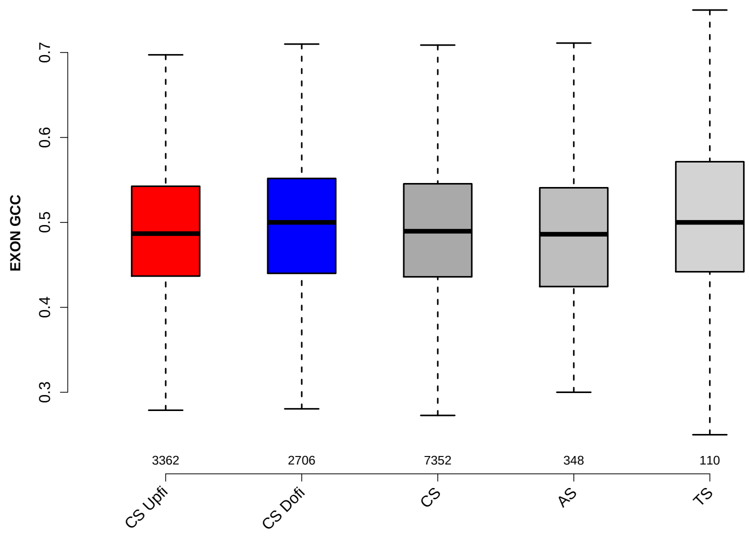

Significant results from Mann-Whitney U test:

- `CS_Upfi` vs `CS_Dofi` : 0.000619771
  mean: 0.489273 < 0.495525 , median: 0.486784 < 0.5
- `CS_Dofi` vs `CS` : 0.0019473
  mean: 0.495525 > 0.490429 , median: 0.5 > 0.489583
- `CS_Dofi` vs `AS` : 0.0160817
  mean: 0.495525 > 0.487212 , median: 0.5 > 0.486046

#### 6.11 UPINTRON GCC

Back to: Overview | ToC

Meaning: GC content of entire up-stream intron sequence

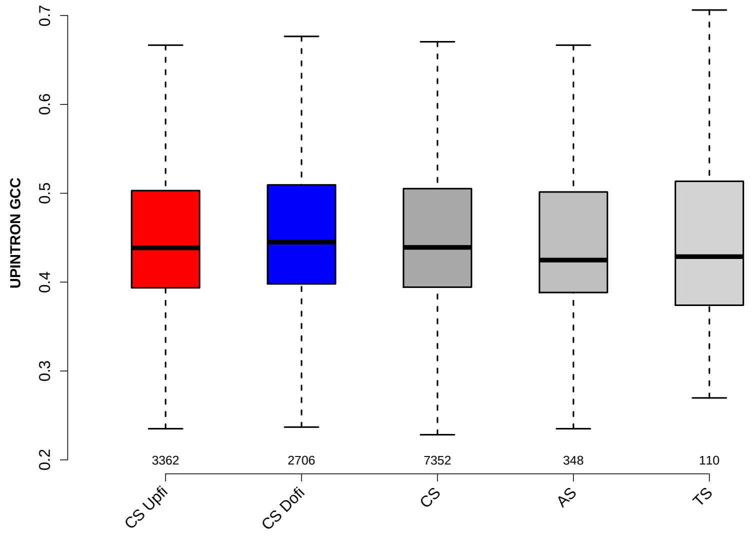

Significant results from Mann-Whitney U test:

- `CS_Upfi` vs `CS_Dofi` : 0.00855573
  mean: 0.450379 < 0.456304 , median: 0.438529 < 0.444956
- `CS_Upfi` vs `AS` : 0.0378502
  mean: 0.450379 > 0.444874 , median: 0.438529 > 0.424791
- `CS_Dofi` vs `CS` : 0.0113577
  mean: 0.456304 > 0.451768 , median: 0.444956 > 0.439085
- `CS_Dofi` vs `AS` : 0.00117554
  mean: 0.456304 > 0.444874 , median: 0.444956 > 0.424791
- `CS` vs `AS` : 0.0196202
  mean: 0.451768 > 0.444874 , median: 0.439085 > 0.424791

#### 6.12 UPEXON GCC

Back to: Overview | ToC

Meaning: GC content of entire up-stream exon sequence

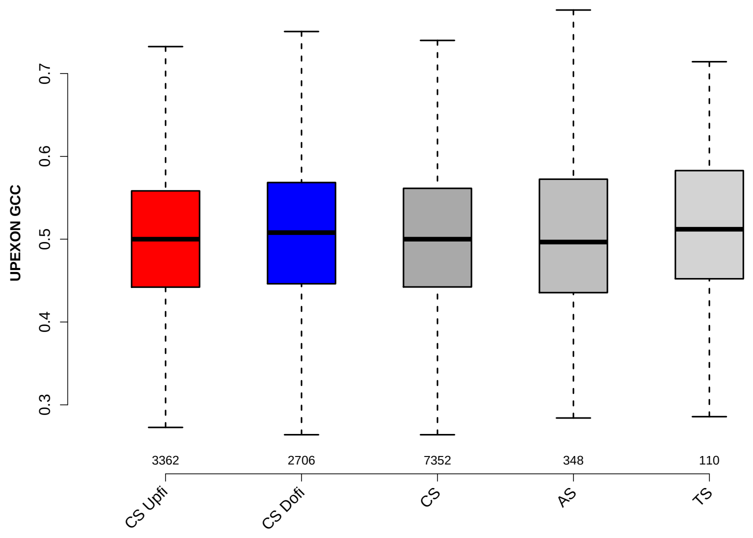

Significant results from Mann-Whitney U test:

- CS_Upfi vs CS_Dofi : 0.0157089
  mean: 0.503419 < 0.509078 , median: 0.5 < 0.507937
- CS_Dofi vs CS : 0.0191853
  mean: 0.509078 > 0.504414 , median: 0.507937 > 0.5

#### 6.13 DOINTRON GCC

Back to: Overview | ToC

Meaning: GC content of entire down-stream intron sequence

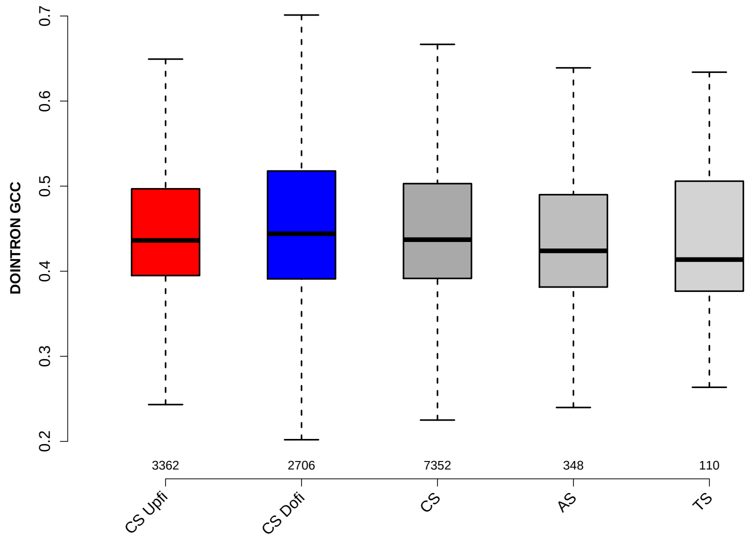

Significant results from Mann-Whitney U test:

- CS_Upfi vs CS_Dofi : 0.0188287
  mean: 0.447729 < 0.452639 , median: 0.436221 < 0.444031
- CS_Upfi vs AS : 0.00514828
  mean: 0.447729 > 0.439282 , median: 0.436221 > 0.423957
- CS_Dofi vs CS : 0.0120043
  mean: 0.452639 > 0.448211 , median: 0.444031 > 0.43709
- CS_Dofi vs AS : 0.000814099
  mean: 0.452639 > 0.439282 , median: 0.444031 > 0.423957
- CS_Dofi vs TS : 0.0431793
  mean: 0.452639 > 0.43681 , median: 0.444031 > 0.413701
- CS vs AS : 0.00728796
  mean: 0.448211 > 0.439282 , median: 0.43709 > 0.423957

#### 6.14 DOEXON GCC

Back to: Overview | ToC

Meaning: GC content of entire down-stream exon sequence

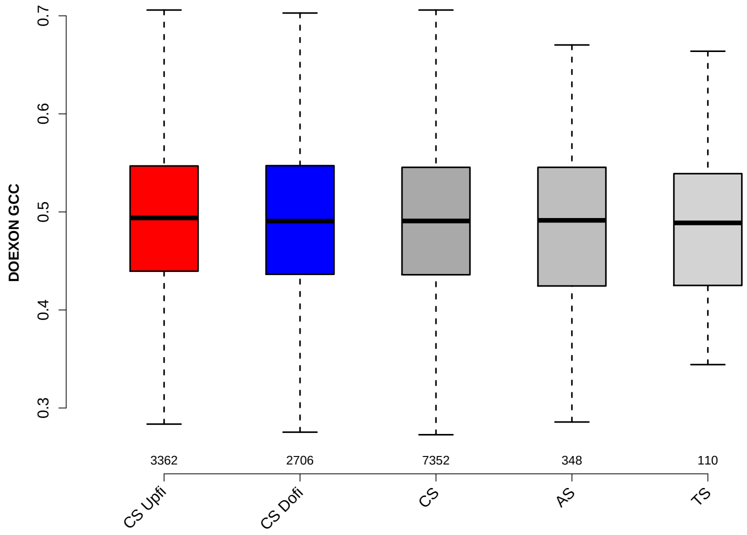

Significant results from Mann-Whitney U test:

- none

#### 6.15 RATIO UPEXON EXON GCC

Back to: Overview | ToC

Meaning: UPEXON GCC / EXON GCC

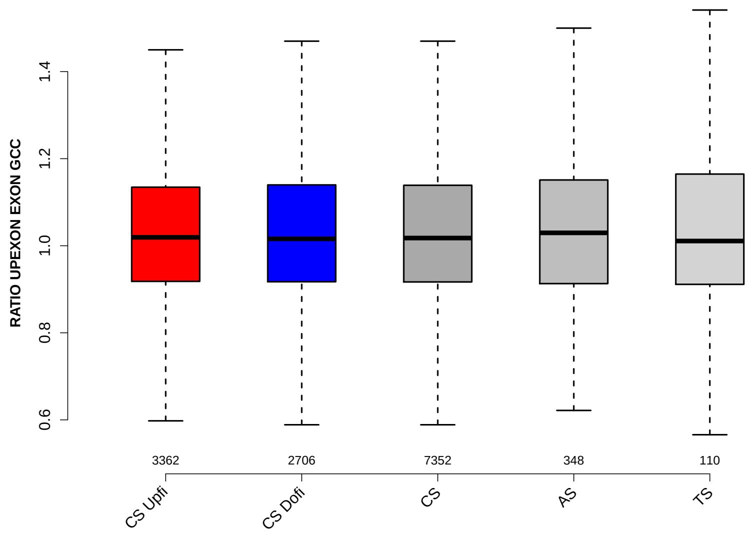

Significant results from Mann-Whitney U test:

- none

#### 6.16 RATIO UPINTRON EXON GCC

Back to: Overview | ToC

Meaning: UPINTRON GCC / EXON GCC

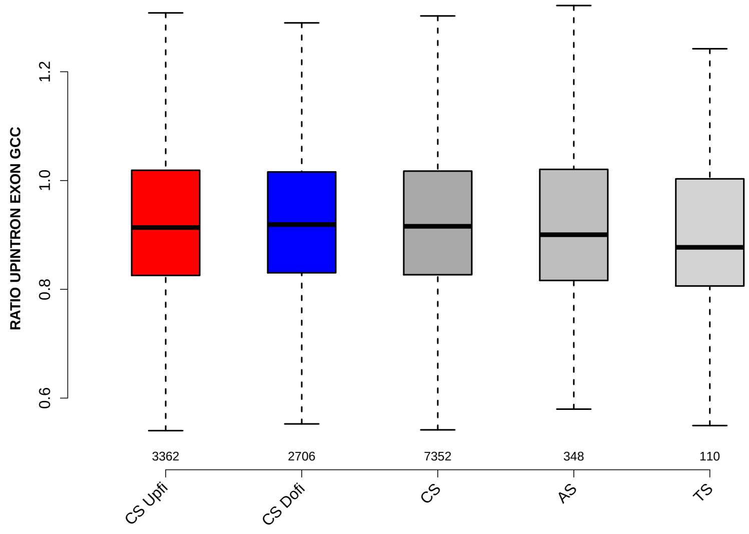

Significant results from Mann-Whitney U test:

- `CS_Dofi` vs `TS` : 0.0329208
  mean: 0.929458 > 0.913539 , median: 0.91904 > 0.877206
- `CS` vs `TS` : 0.0394753
  mean: 0.929886 > 0.913539 , median: 0.915879 > 0.877206

#### 6.17 RATIO DOINTRON EXON GCC

Back to: Overview | ToC

Meaning: DOINTRON GCC / EXON GCC

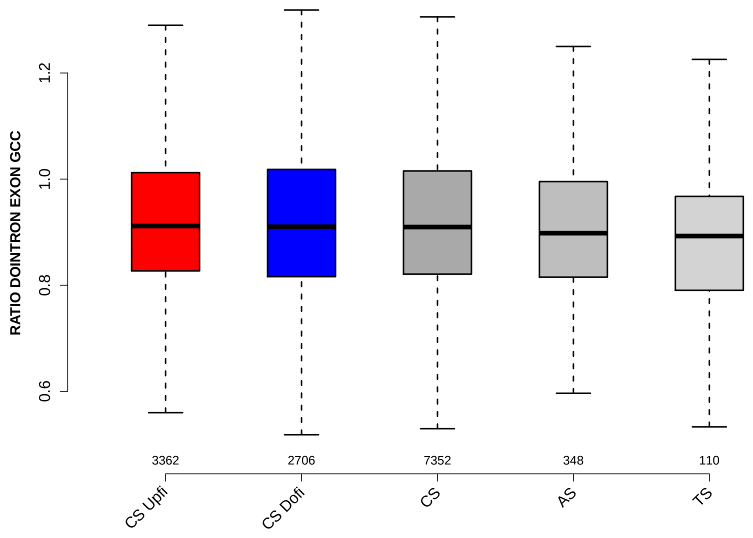

Significant results from Mann-Whitney U test:

- `CS_Upfi` vs `TS` : 0.00940976
  mean: 0.925535 > 0.890057 , median: 0.911598 > 0.892731
- `CS_Dofi` vs `TS` : 0.0231156
  mean: 0.920836 > 0.890057 , median: 0.910379 > 0.892731
- `CS` vs `TS` : 0.0157486
  mean: 0.923309 > 0.890057 , median: 0.909696 > 0.892731

#### 6.18 RATIO DOEXON EXON GCC

Back to: Overview | ToC

Meaning: DOEXON GCC / EXON GCC

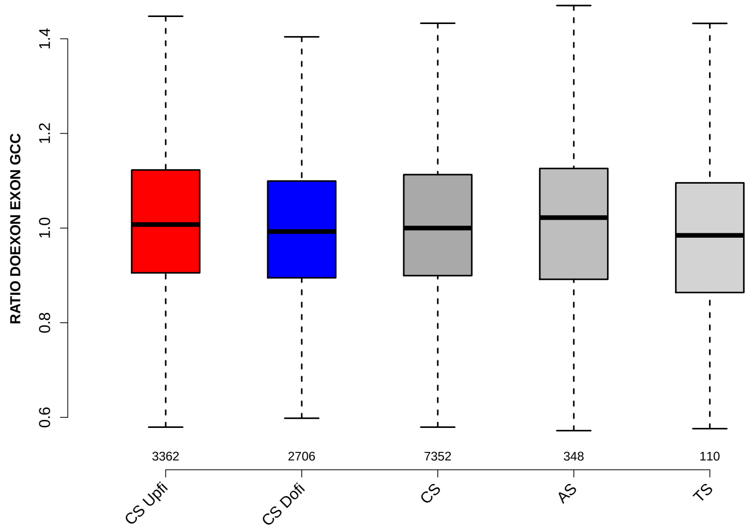

Significant results from Mann-Whitney U test:

- CS_Upfi vs CS_Dofi : 0.000343999
  mean: 1.0195 > 1.0051 , median: 1.0074 > 0.992711
- CS_Upfi vs CS : 0.0290437
  mean: 1.0195 > 1.0127 , median: 1.0074 > 1
- CS_Dofi vs CS : 0.0386185
  mean: 1.0051 < 1.0127 , median: 0.992711 < 1

#### 6.19 SF1 HIGHESTSCORE 3SS UPINTRON

Back to: Overview | ToC

Meaning: highest score of a SF1 positon weight matrix trained with human data in the last 150 nt 3 prime intron positons of up-stream intron

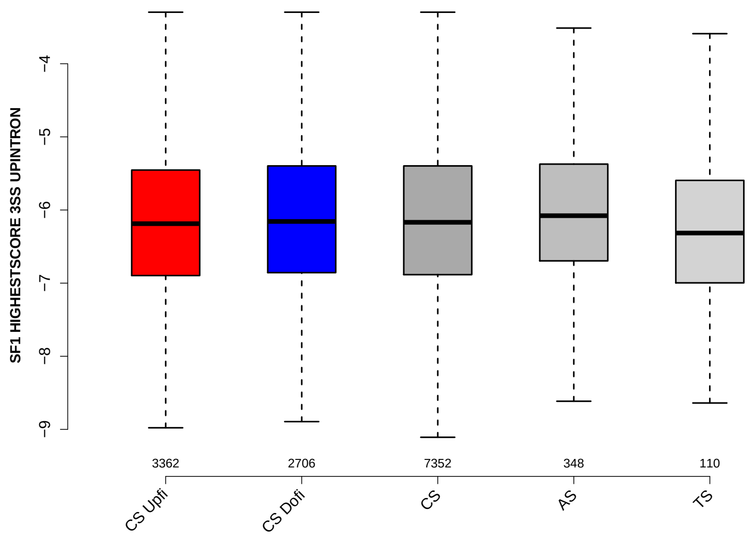

Significant results from Mann-Whitney U test:

- `CS_Upfi` vs `AS` : 0.0206018
  mean: -6.1775 < -6.0383 , median: -6.18725 < -6.07814
- `CS` vs `AS` : 0.049154
  mean: -6.15077 < -6.0383 , median: -6.1683 < -6.07814

#### 6.20 SF1 HIGHESTSCORE 3SS DOINTRON

Back to: Overview | ToC

Meaning: highest score of a SF1 positon weight matrix trained with human data in the last 150 nt 3 prime intron positons of down-stream intron

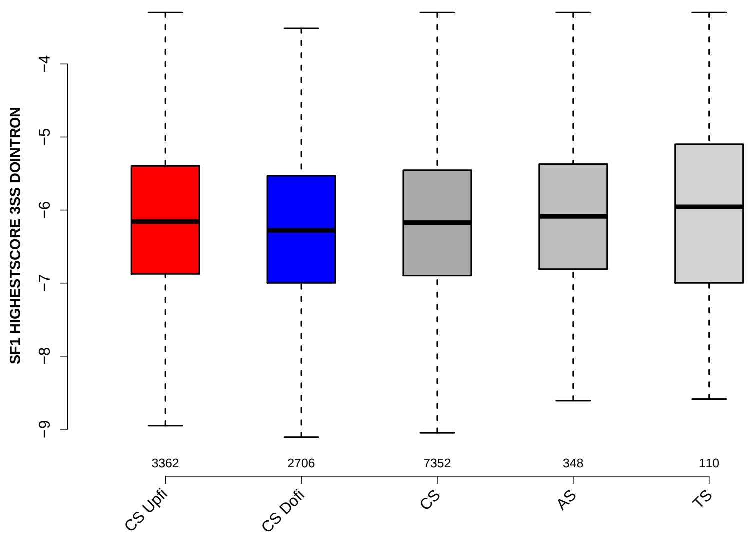

Significant results from Mann-Whitney U test:

- `CS_Upfi` vs `CS_Dofi` : 0.00015327
  mean: -6.13092 > -6.24778 , median: -6.15674 > -6.27734
- `CS_Dofi` vs `CS` : 0.00585974
  mean: -6.24778 < -6.17718 , median: -6.27734 < -6.17225
- `CS_Dofi` vs `AS` : 0.000971733
  mean: -6.24778 < -6.03364 , median: -6.27734 < -6.08558
- `CS` vs `AS` : 0.0191529
  mean: -6.17718 < -6.03364 , median: -6.17225 < -6.08558

#### 6.21 UP 5SS 20INT10EX GCC

Back to: Overview | ToC

Meaning: GC content of up-stream 5ss sequence (20int+10ex positions)

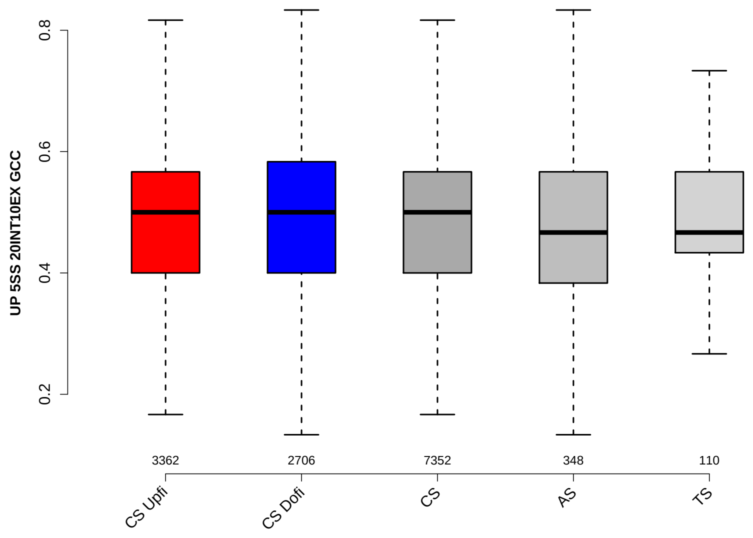

Significant results from Mann-Whitney U test:

- CS_Upfi vs CS_Dofi : 0.00185647
  mean: 0.484776 $<$ 0.495799 , median: 0.5 $=$ 0.5
- CS_Dofi vs CS : 0.00493331
  mean: 0.495799 $>$ 0.487384 , median: 0.5 $=$ 0.5
- CS_Dofi vs AS : 0.0285229
  mean: 0.495799 $>$ 0.48137 , median: 0.5 $>$ 0.466667

##### 6.22 GCC 3SS 20INT10EX

Back to: Overview | ToC

Meaning: GC content of 3ss sequence (20int+10ex positions)

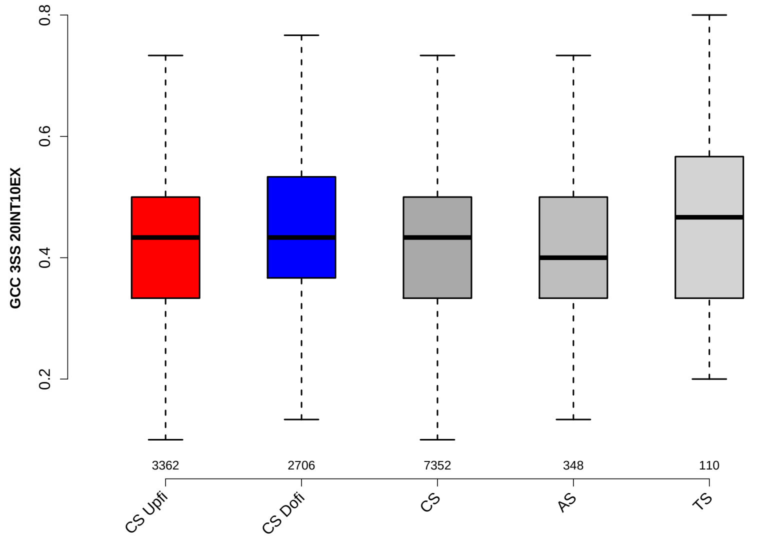

Significant results from Mann-Whitney U test:

- `CS_Upfi` vs `CS_Dofi` : 2.07363e-06
  mean: 0.427761 < 0.442104 , median: 0.433333 = 0.433333
- `CS_Upfi` vs `TS` : 0.0185486
  mean: 0.427761 < 0.459394 , median: 0.433333 < 0.466667
- `CS_Dofi` vs `CS` : 0.0001844
  mean: 0.442104 > 0.432395 , median: 0.433333 = 0.433333
- `CS_Dofi` vs `AS` : 0.000692387
  mean: 0.442104 > 0.422701 , median: 0.433333 > 0.4
- `CS` vs `AS` : 0.0326909
  mean: 0.432395 > 0.422701 , median: 0.433333 > 0.4
- `CS` vs `TS` : 0.0451558
  mean: 0.432395 < 0.459394 , median: 0.433333 < 0.466667
- `AS` vs `TS` : 0.0102299
  mean: 0.422701 < 0.459394 , median: 0.4 < 0.466667

#### 6.23 GCC 5SS 20INT10EX

Back to: Overview | ToC

Meaning: GC content of 5ss sequence (20int+10ex positions)

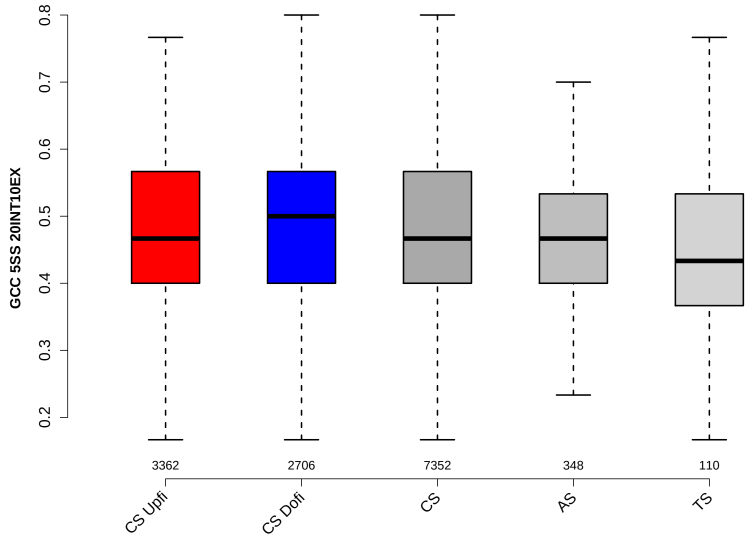

Significant results from Mann-Whitney U test:

- `CS_Dofi` vs `AS` : 0.0366631
  mean: 0.477987 > 0.46887 , median: 0.5 > 0.466667

#### 6.24 DO 3SS 20INT10EX GCC

Back to: Overview | ToC

Meaning: GC content of down-stream 3ss sequence (20int+10ex positions)

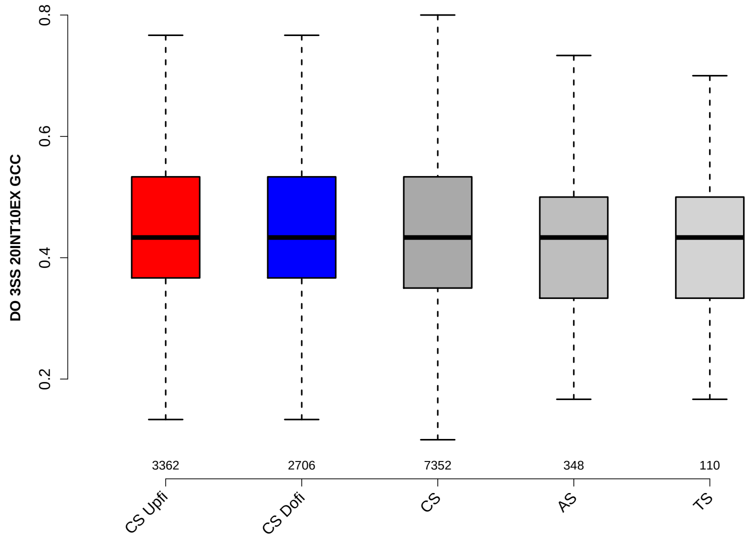

Significant results from Mann-Whitney U test:

- `CS_Upfi` vs `AS` : 0.0264216
  mean: 0.438786 > 0.425718 , median: 0.433333 = 0.433333
- `CS_Dofi` vs `AS` : 0.0418792
  mean: 0.438248 > 0.425718 , median: 0.433333 = 0.433333

#### 6.25 MAXENTSCR HSAMODEL UPSTRM 5SS

Back to: Overview | ToC

Meaning: maximum entropy score of 5ss of up-stream exon using a model trained with human splice sites

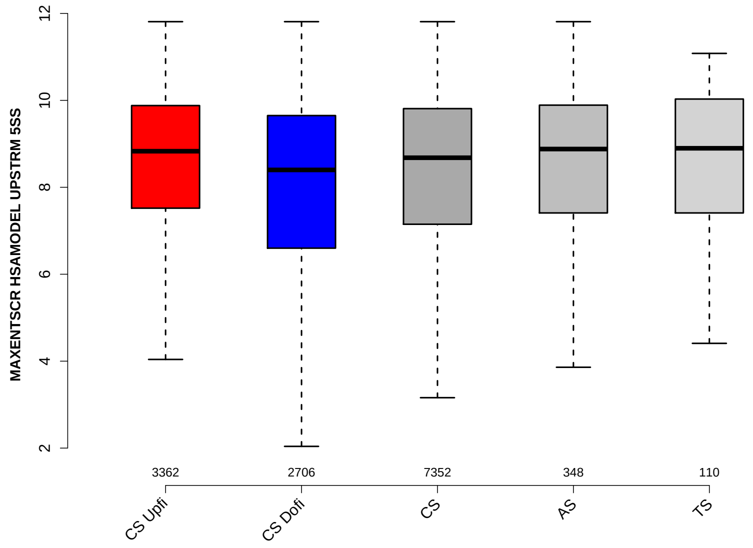

Significant results from Mann-Whitney U test:

- `CS_Upfi` vs `CS_Dofi` : 2.79369e-23
  mean: 8.4917 > 7.7588 , median: 8.83 > 8.4
- `CS_Upfi` vs `CS` : 4.5223e-09
  mean: 8.4917 > 8.1554 , median: 8.83 > 8.68
- `CS_Dofi` vs `CS` : 1.7306e-09
  mean: 7.7588 < 8.1554 , median: 8.4 < 8.68
- `CS_Dofi` vs `AS` : 0.000552638
  mean: 7.7588 < 8.2038 , median: 8.4 < 8.88
- `CS_Dofi` vs `TS` : 0.0142251
  mean: 7.7588 < 8.3675 , median: 8.4 < 8.8975

#### 6.26 MAXENTSCR HSAMODEL 3SS

Back to: Overview | ToC

Meaning: maximum entropy score of 3ss using a model trained with human splice sites

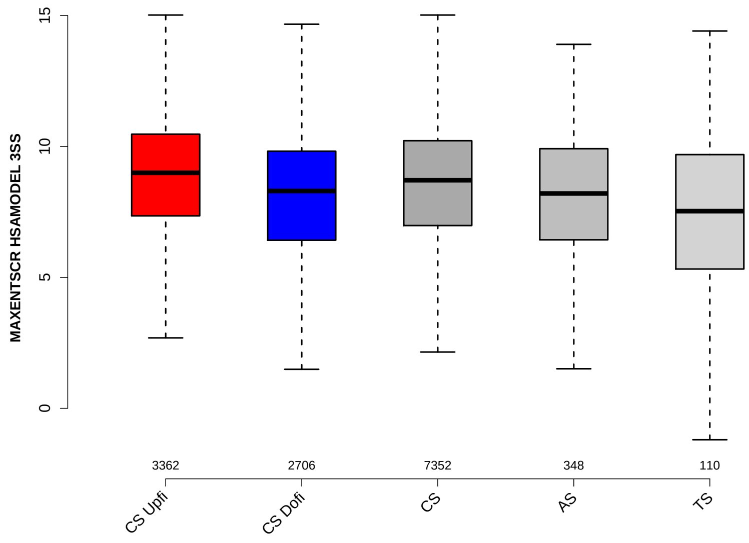

Significant results from Mann-Whitney U test:

- `CS_Upfi` vs `CS_Dofi` : 3.82506e-30
  mean: 8.814 > 8.019 , median: 8.995 > 8.3
- `CS_Upfi` vs `CS` : 3.47164e-09
  mean: 8.814 > 8.4795 , median: 8.995 > 8.71
- `CS_Upfi` vs `AS` : 6.46419e-08
  mean: 8.814 > 7.8019 , median: 8.995 > 8.205
- `CS_Upfi` vs `TS` : 4.5275e-07
  mean: 8.814 > 7.1541 , median: 8.995 > 7.53
- `CS_Dofi` vs `CS` : 2.26536e-14
  mean: 8.019 < 8.4795 , median: 8.3 < 8.71
- `CS_Dofi` vs `TS` : 0.0150106
  mean: 8.019 > 7.1541 , median: 8.3 > 7.53
- `CS` vs `AS` : 0.000672383
  mean: 8.4795 > 7.8019 , median: 8.71 > 8.205
- `CS` vs `TS` : 6.59356e-05
  mean: 8.4795 > 7.1541 , median: 8.71 > 7.53

#### 6.27 MAXENTSCR HSAMODEL 5SS

Back to: Overview | ToC

Meaning: maximum entropy score of 5ss using a model trained with human splice sites

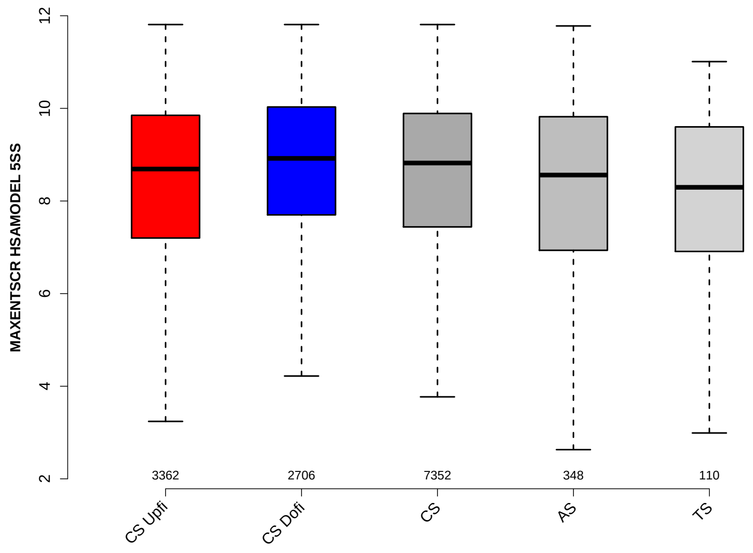

Significant results from Mann-Whitney U test:

- CS_Upfi vs CS_Dofi : 6.62158e-09
  mean: 8.149 < 8.5907 , median: 8.69 < 8.92
- CS_Upfi vs CS : 0.000635509
  mean: 8.149 < 8.3563 , median: 8.69 < 8.82
- CS_Upfi vs TS : 0.041549
  mean: 8.149 > 7.4175 , median: 8.69 > 8.295
- CS_Dofi vs CS : 0.000470766
  mean: 8.5907 > 8.3563 , median: 8.92 > 8.82
- CS_Dofi vs AS : 0.000126052
  mean: 8.5907 > 7.8916 , median: 8.92 > 8.56
- CS_Dofi vs TS : 0.000184394
  mean: 8.5907 > 7.4175 , median: 8.92 > 8.295
- CS vs AS : 0.0116221
  mean: 8.3563 > 7.8916 , median: 8.82 > 8.56
- CS vs TS : 0.00401386
  mean: 8.3563 > 7.4175 , median: 8.82 > 8.295

#### 6.28 MAXENTSCR HSAMODEL DOWNSTRM 3SS

Back to: Overview | ToC

Meaning: maximum entropy score of 3ss of down-stream exon using a model trained with human splice sites

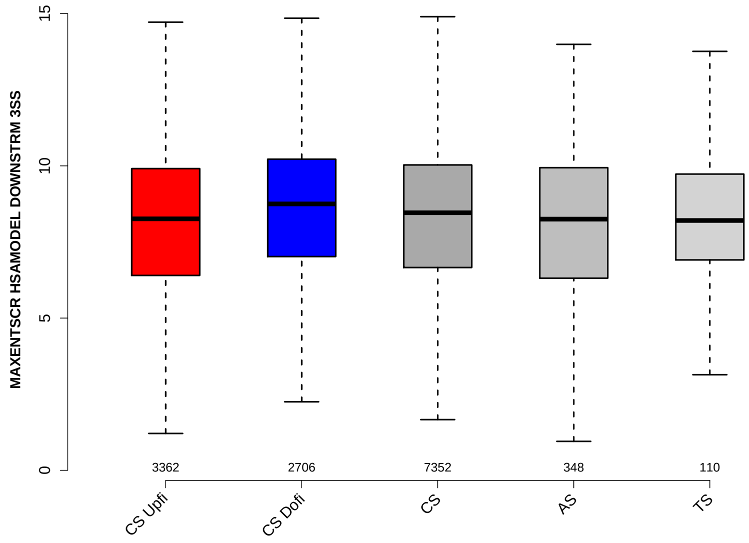

Significant results from Mann-Whitney U test:

- CS_Upfi vs CS_Dofi : 6.12101e-13
  mean: 7.9952 < 8.5475 , median: 8.26 < 8.75
- CS_Upfi vs CS : 0.000500696
  mean: 7.9952 < 8.2116 , median: 8.26 < 8.46
- CS_Dofi vs CS : 4.26202e-07
  mean: 8.5475 > 8.2116 , median: 8.75 > 8.46
- CS_Dofi vs AS : 0.00335947
  mean: 8.5475 > 8.1038 , median: 8.75 > 8.25

#### 6.29 DIST FROM MAXBP TO 3SS UPINTRON

Back to: Overview | ToC

Meaning: distance to 3ss of best precited BP

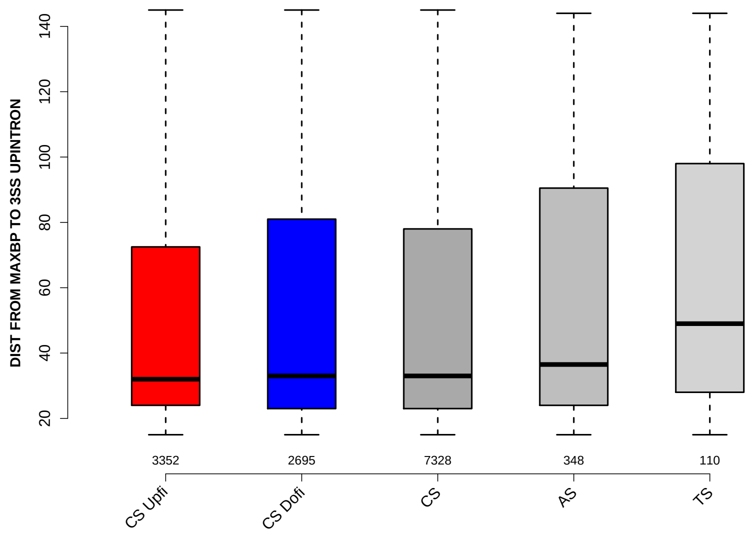

Significant results from Mann-Whitney U test:

- `CS_Upfi` vs `AS` : 0.0408239
  mean: 51.1116 < 56.4626 , median: 32 < 36.5
- `CS_Upfi` vs `TS` : 0.00142745
  mean: 51.1116 < 62.7455 , median: 32 < 49
- `CS_Dofi` vs `TS` : 0.0058158
  mean: 53.4471 < 62.7455 , median: 33 < 49
- `CS` vs `TS` : 0.00281747
  mean: 52.3594 < 62.7455 , median: 33 < 49

#### 6.30 SCORE FOR MAXBP SEQ UPINTRON

Back to: Overview | ToC

Meaning: BP sequence score of best predicted BP

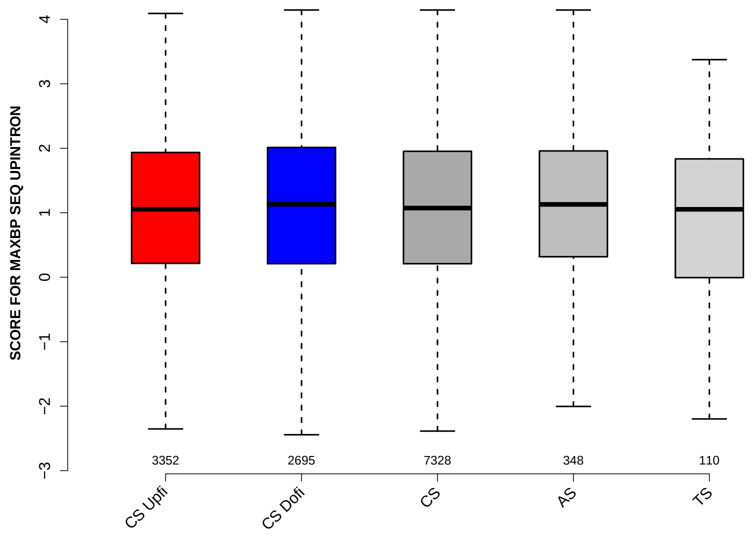

Significant results from Mann-Whitney U test:

- `CS_Upfi` vs `CS_Dofi` : 0.0311061
  mean: $1.0274 < 1.1022$ , median: $1.0545 < 1.1319$

#### 6.31 PYRIMIDINECONT MAXBP UPINTRON

Back to: Overview | ToC

Meaning: Pyrimidine content between the BP adenine and the 3 prime splice site for best BP

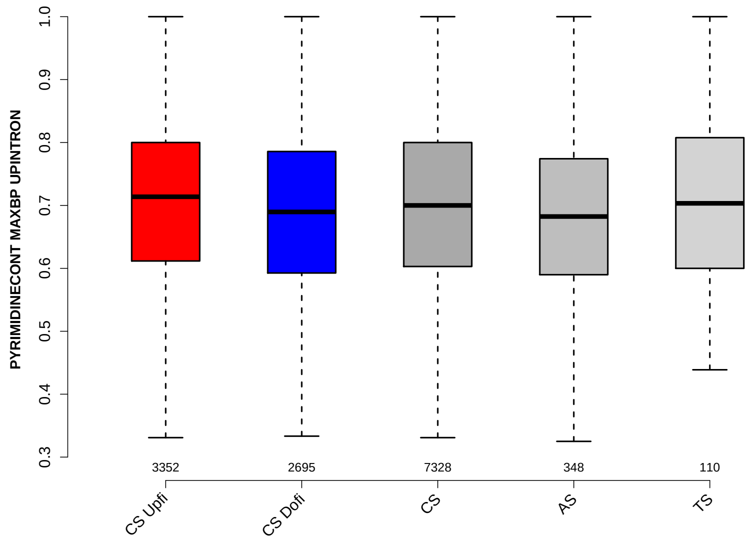

Significant results from Mann-Whitney U test:

- CS_Upfi vs CS_Dofi : 4.02367e-08
  mean: 0.710564 > 0.692165 , median: 0.713761 > 0.689655
- CS_Upfi vs CS : 0.00248731
  mean: 0.710564 > 0.702439 , median: 0.713761 > 0.7
- CS_Upfi vs AS : 0.00104654
  mean: 0.710564 > 0.686989 , median: 0.713761 > 0.682373
- CS_Dofi vs CS : 0.00045694
  mean: 0.692165 < 0.702439 , median: 0.689655 < 0.7
- CS vs AS : 0.0283894
  mean: 0.702439 > 0.686989 , median: 0.7 > 0.682373

#### 6.32 POLYPYRITRAC OFFSET MAXBP UPINTRON

Back to: Overview | ToC

Meaning: Polypyrimidine track offset relative to the BP adenine for best BP

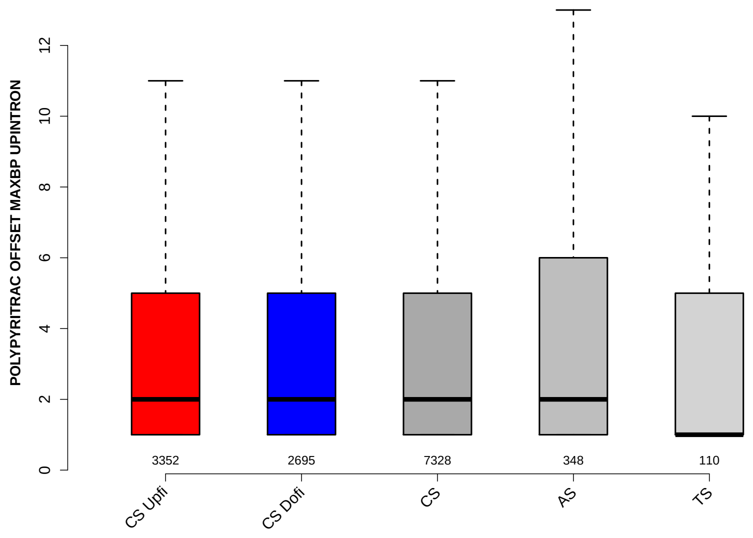

Significant results from Mann-Whitney U test:

- none

#### 6.33 POLYPYRITRAC LEN MAXBP UPINTRON

Back to: Overview | ToC

Meaning: Polypyrimidine track length for best BP

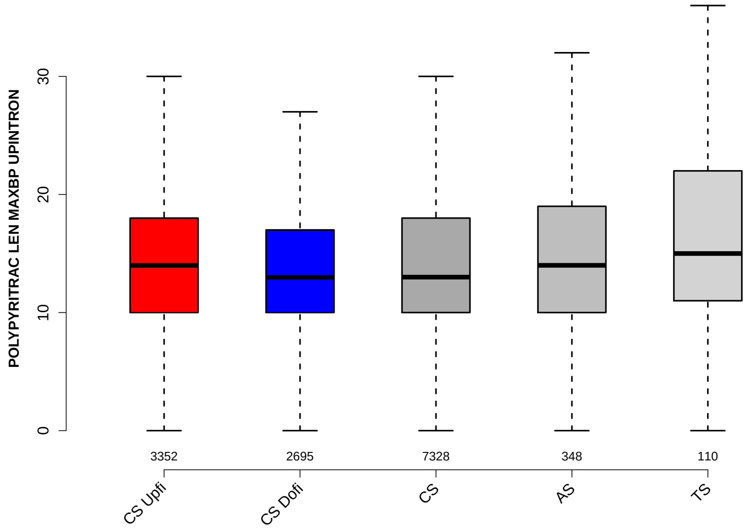

Significant results from Mann-Whitney U test:

- `CS_Upfi` vs `CS_Dofi` : 0.00113712
  mean: 15.1184 > 14.5102 , median: 14 > 13
- `CS_Upfi` vs `TS` : 0.0223696
  mean: 15.1184 < 19.5909 , median: 14 < 15
- `CS_Dofi` vs `CS` : 0.0339002
  mean: 14.5102 < 14.8691 , median: 13 = 13
- `CS_Dofi` vs `TS` : 0.00267445
  mean: 14.5102 < 19.5909 , median: 13 < 15
- `CS` vs `TS` : 0.00888757
  mean: 14.8691 < 19.5909 , median: 13 < 15
- `AS` vs `TS` : 0.0213521
  mean: 15.0029 < 19.5909 , median: 14 < 15

#### 6.34 POLYPYRITRAC SCORE MAXBP UPINTRON

Back to: Overview | ToC

Meaning: Polypyrimidine track score for best BP

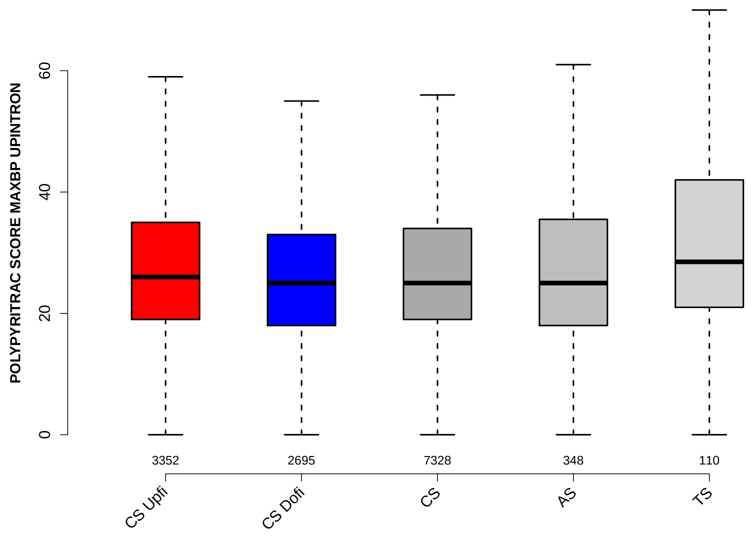

Significant results from Mann-Whitney U test:

- CS_Upfi vs CS_Dofi : 1.90603e-05
  mean: 29.0779 > 27.2508 , median: 26 > 25
- CS_Upfi vs CS : 0.0450509
  mean: 29.0779 > 28.3682 , median: 26 > 25
- CS_Upfi vs TS : 0.0185998
  mean: 29.0779 < 36.6909 , median: 26 < 28.5
- CS_Dofi vs CS : 0.00223216
  mean: 27.2508 < 28.3682 , median: 25 = 25
- CS_Dofi vs TS : 0.000632609
  mean: 27.2508 < 36.6909 , median: 25 < 28.5
- CS vs TS : 0.00534121
  mean: 28.3682 < 36.6909 , median: 25 < 28.5
- AS vs TS : 0.0121506
  mean: 28.4828 < 36.6909 , median: 25 < 28.5

#### 6.35 BPSCORE MAXBP UPINTRON

Back to: Overview | ToC

Meaning: SVM classification score of best BP

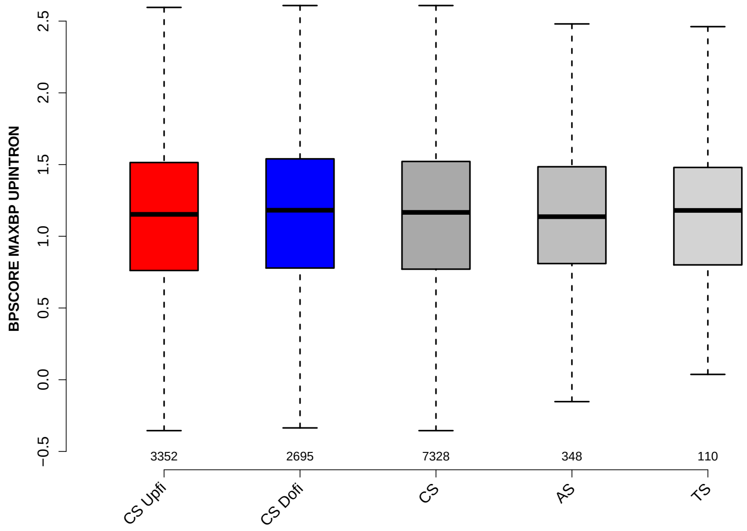

Significant results from Mann-Whitney U test:

- none

#### 6.36 NUM PREDICTED BPS UPINTRON

Back to: Overview | ToC

Meaning: number of all predicted BPs which have a positive BP score

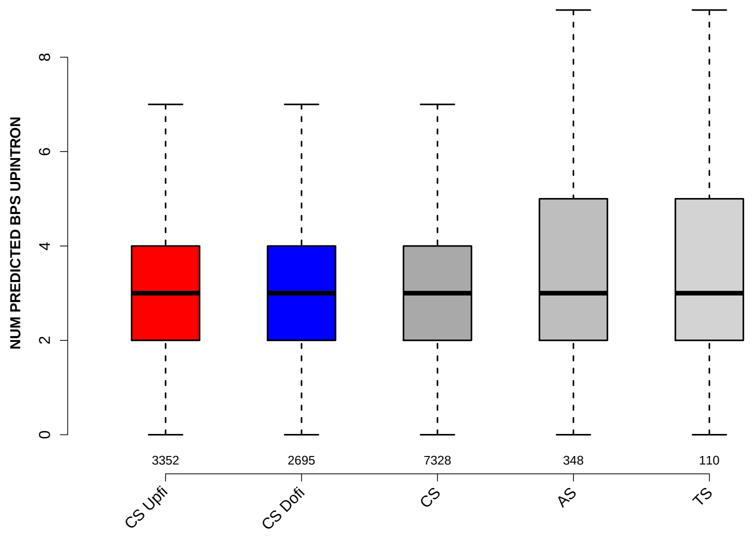

Significant results from Mann-Whitney U test:

- `CS_Upfi` vs `AS` : 0.0347717
  mean: 3.2405 $<$ 3.4483 , median: 3 $=$ 3

#### 6.37 MEDIAN DIST FROM BP TO 3SS UPINTRON

Back to: Overview | ToC

Meaning: like DIST FROM MAXBP TO 3SS but median over top-3 predicted BPs

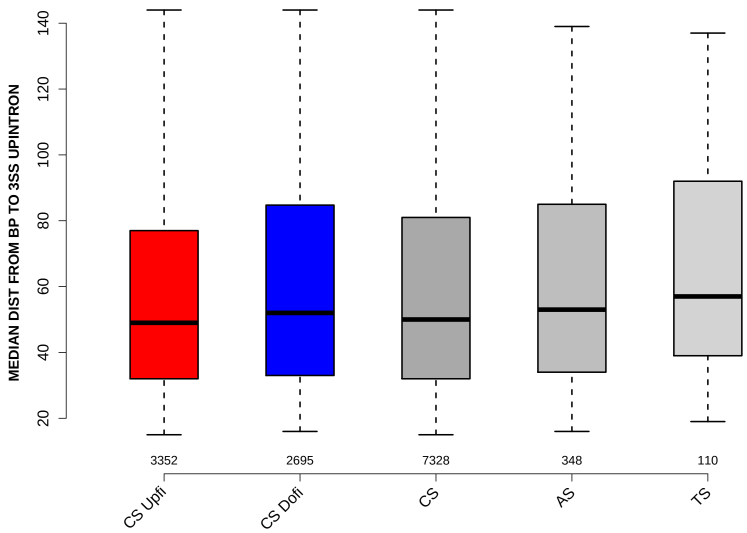

Significant results from Mann-Whitney U test:

- `CS_Upfi` vs `CS_Dofi` : 0.000858255
  mean: 57.4241 < 60.4377 , median: 49 < 52
- `CS_Upfi` vs `AS` : 0.0172388
  mean: 57.4241 < 61.819 , median: 49 < 53
- `CS_Upfi` vs `TS` : 0.00435542
  mean: 57.4241 < 65.9727 , median: 49 < 57
- `CS_Dofi` vs `CS` : 0.0261701
  mean: 60.4377 > 58.7454 , median: 52 > 50
- `CS` vs `TS` : 0.0138374
  mean: 58.7454 < 65.9727 , median: 50 < 57

#### 6.38 MEDIAN SCORE FOR BPSEQ UPINTRON

Back to: Overview | ToC

Meaning: like SCORE FOR MAXBP SEQ but median over top-3 predicted BPs

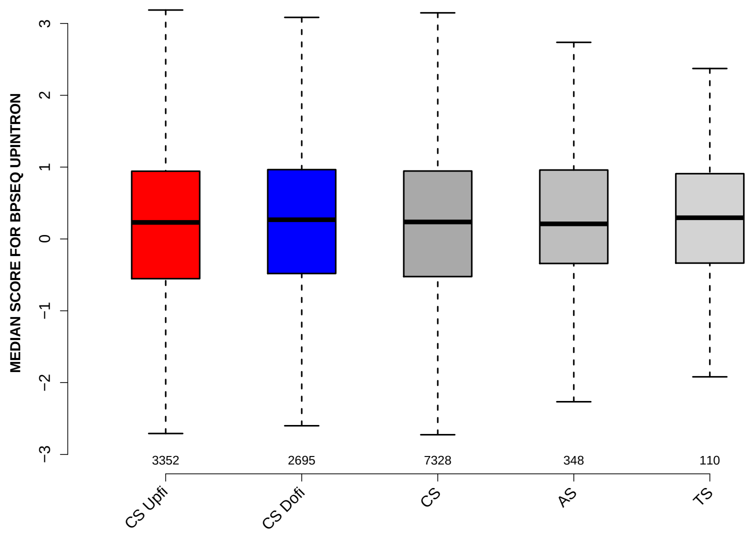

Significant results from Mann-Whitney U test:

- none

#### 6.39 MEDIAN PYRIMIDINECONT UPINTRON

Back to: Overview | ToC

Meaning: like PYRIMIDINECONT MAXBP but median over top-3 predicted BPs

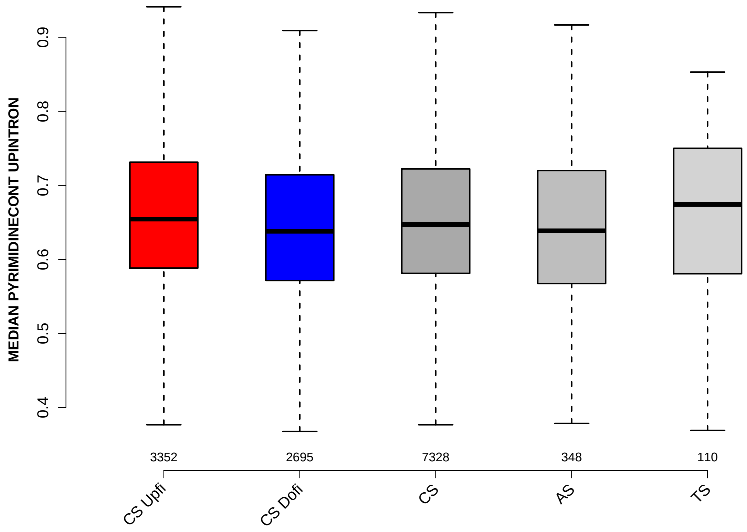

Significant results from Mann-Whitney U test:

- `CS_Upfi` vs `CS_Dofi` : 5.55741e-11
  mean: 0.661475 > 0.643318 , median: 0.654433 > 0.637931
- `CS_Upfi` vs `CS` : 0.000486214
  mean: 0.661475 > 0.653727 , median: 0.654433 > 0.647059
- `CS_Upfi` vs `AS` : 0.00464829
  mean: 0.661475 > 0.643806 , median: 0.654433 > 0.638593
- `CS_Dofi` vs `CS` : 1.718e-05
  mean: 0.643318 < 0.653727 , median: 0.637931 < 0.647059
- `CS_Dofi` vs `TS` : 0.0336011
  mean: 0.643318 < 0.660946 , median: 0.637931 < 0.674235

#### 6.40 MEDIAN POLYPYRITRAC OFFSET UPINTRON

Back to: Overview | ToC

Meaning: like POLYPYRITRAC OFFSET MAXBP but median over top-3 predicted BPs

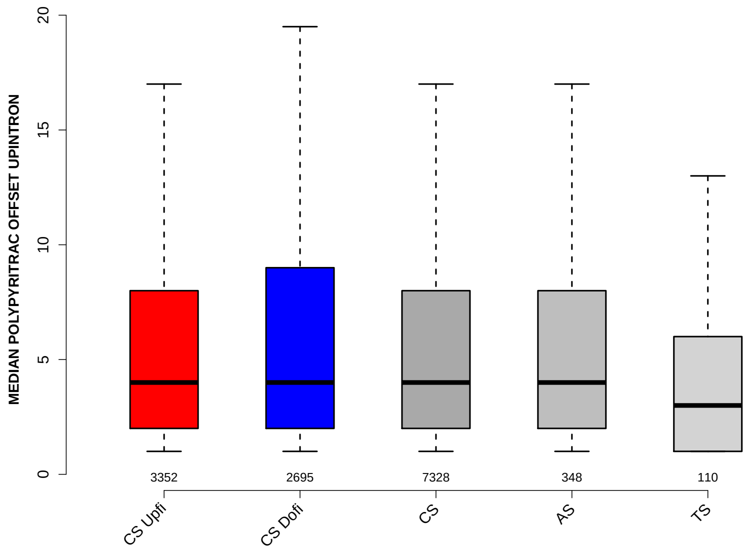

Significant results from Mann-Whitney U test:

- `CS_Dofi` vs `TS` : 0.0208119
  mean: 7.0301 < 7.3 , median: 4 > 3
- `CS` vs `TS` : 0.0403892
  mean: 6.5806 < 7.3 , median: 4 > 3

#### 6.41 MEDIAN POLYPYRITRAC LEN UPINTRON

Back to: Overview | ToC

Meaning: like POLYPYRITRAC LEN MAXBP but median over top-3 predicted BPs

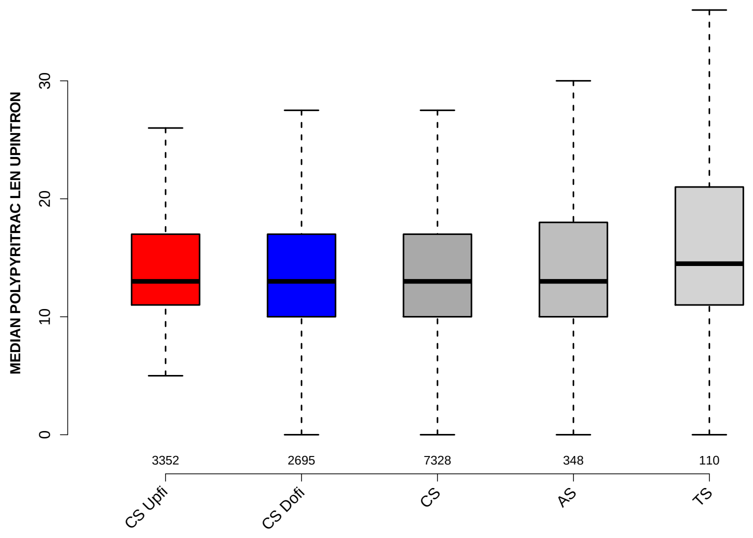

Significant results from Mann-Whitney U test:

- `CS_Upfi` vs `CS_Dofi` : 0.000507058
  mean: 14.751 > 14.2826 , median: 13 = 13
- `CS_Dofi` vs `CS` : 0.0272852
  mean: 14.2826 < 14.5568 , median: 13 = 13
- `CS_Dofi` vs `TS` : 0.0146261
  mean: 14.2826 < 19.0091 , median: 13 < 14.5
- `CS` vs `TS` : 0.0405443
  mean: 14.5568 < 19.0091 , median: 13 < 14.5

#### 6.42 MEDIAN POLYPYRITRAC SCORE UPINTRON

Back to: Overview | ToC

Meaning: like POLYPYRITRAC SCORE MAXBP but median over top-3 predicted BPs

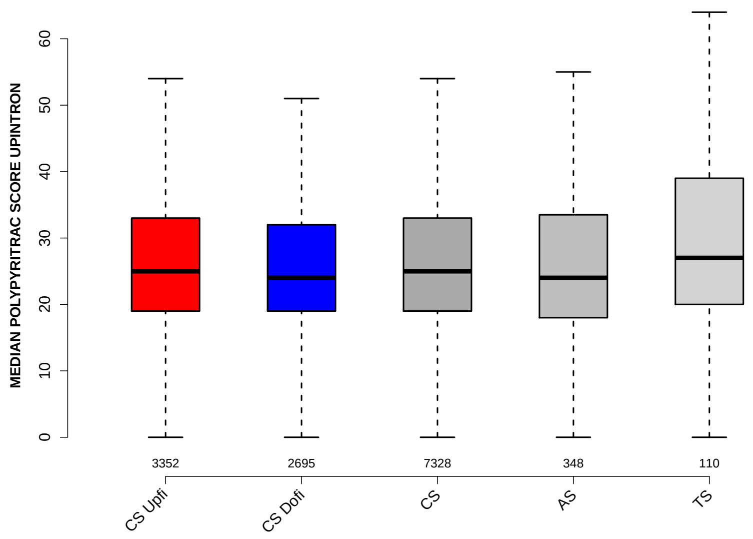

Significant results from Mann-Whitney U test:

- CS_Upfi vs CS_Dofi : 3.42929e-05
  mean: 28.105 > 26.7072 , median: 25 > 24
- CS_Upfi vs CS : 0.0407757
  mean: 28.105 > 27.5461 , median: 25 = 25
- CS_Dofi vs CS : 0.00428984
  mean: 26.7072 < 27.5461 , median: 24 < 25
- CS_Dofi vs TS : 0.00786612
  mean: 26.7072 < 35.8227 , median: 24 < 27
- CS vs TS : 0.0348223
  mean: 27.5461 < 35.8227 , median: 25 < 27
- AS vs TS : 0.0418988
  mean: 27.5244 < 35.8227 , median: 24 < 27

#### 6.43 MEDIAN BPSCORE UPINTRON

Back to: Overview | ToC

Meaning: like BPSCORE MAXBP but median over top-3 predicted BPs

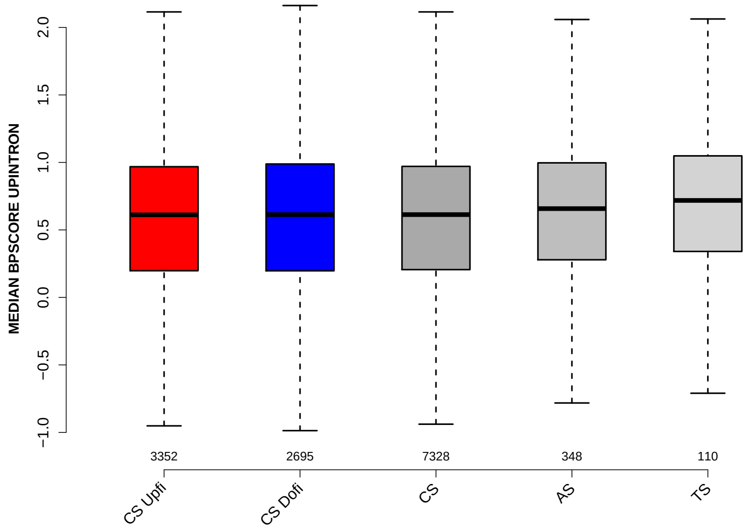

Significant results from Mann-Whitney U test:

- none

#### 6.44 DIST FROM MAXBP TO 3SS DOINTRON

Back to: Overview | ToC

Meaning: distance to 3ss of best precited BP

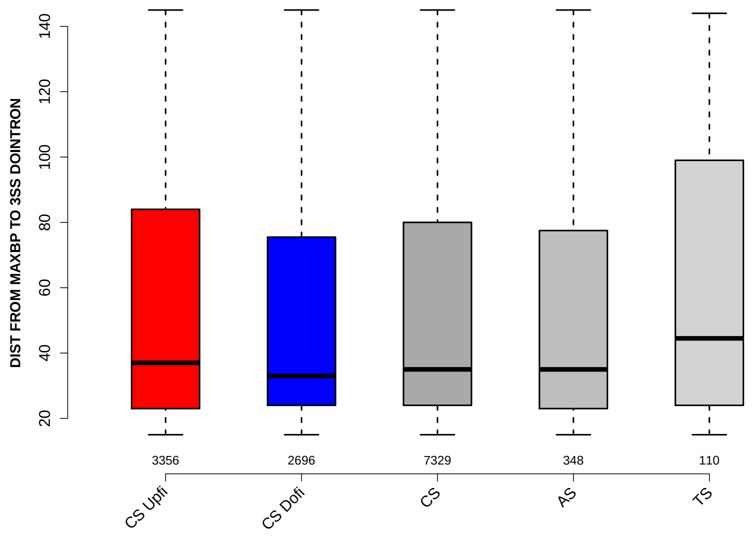

Significant results from Mann-Whitney U test:

- `CS_Upfi` vs `CS_Dofi` : 0.00585407
  mean: 55.1311 > 51.4989 , median: 37 > 33
- `CS_Dofi` vs `CS` : 0.047329
  mean: 51.4989 < 53.6578 , median: 33 < 35
- `CS_Dofi` vs `TS` : 0.0335349
  mean: 51.4989 < 61.3 , median: 33 < 44.5

#### 6.45 SCORE FOR MAXBP SEQ DOINTRON

Back to: Overview | ToC

Meaning: BP sequence score of best predicted BP

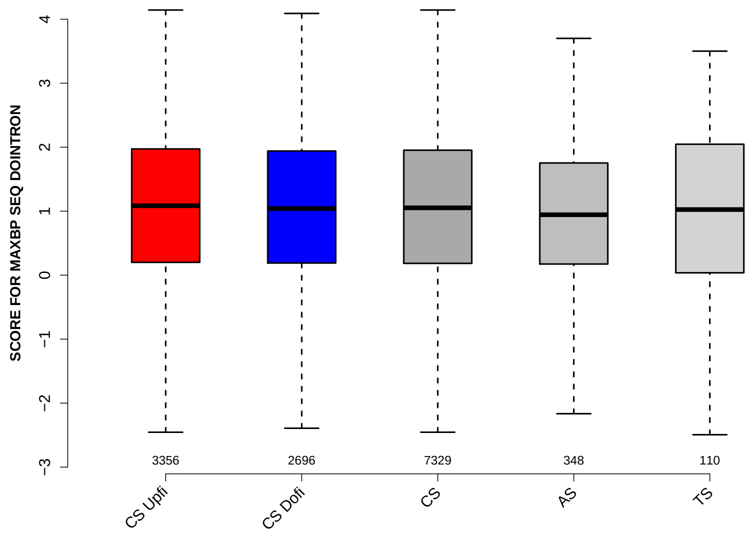

Significant results from Mann-Whitney U test:

- none

#### 6.46 PYRIMIDINECONT MAXBP DOINTRON

Back to: Overview | ToC

Meaning: Pyrimidine content between the BP adenine and the 3 prime splice site for best BP

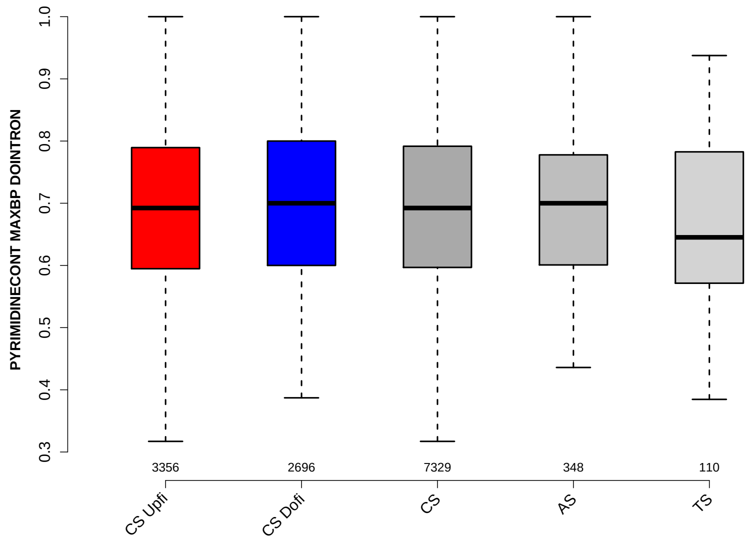

Significant results from Mann-Whitney U test:

- `CS_Dofi` vs `TS` : 0.0254834
  mean: 0.70023 > 0.672514 , median: 0.7 > 0.645161

#### 6.47 POLYPYRITRAC OFFSET MAXBP DOINTRON

Back to: Overview | ToC

Meaning: Polypyrimidine track offset relative to the BP adenine for best BP

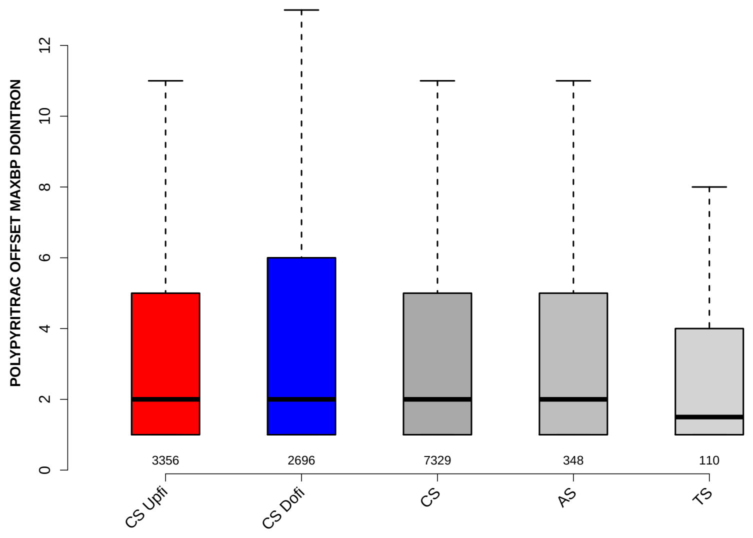

Significant results from Mann-Whitney U test:

- none

#### 6.48 POLYPYRITRAC LEN MAXBP DOINTRON

Back to: Overview | ToC

Meaning: Polypyrimidine track length for best BP

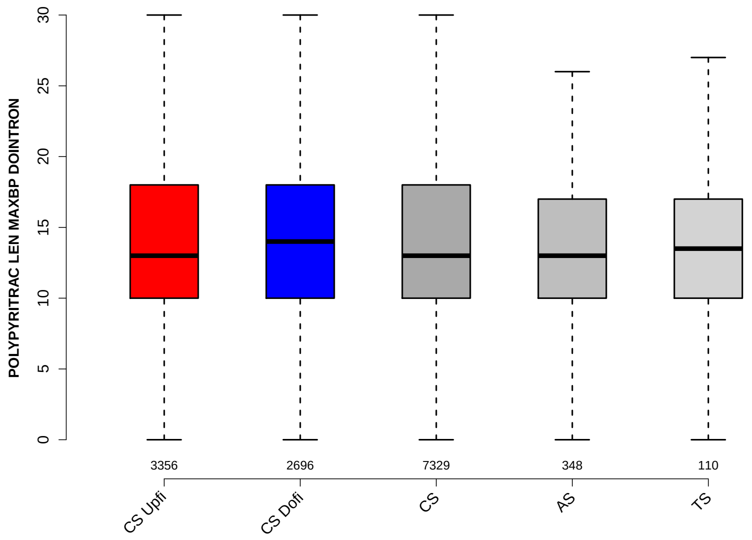

Significant results from Mann-Whitney U test:
- none

#### 6.49 POLYPYRITRAC SCORE MAXBP DOINTRON

Back to: Overview | ToC

Meaning: Polypyrimidine track score for best BP

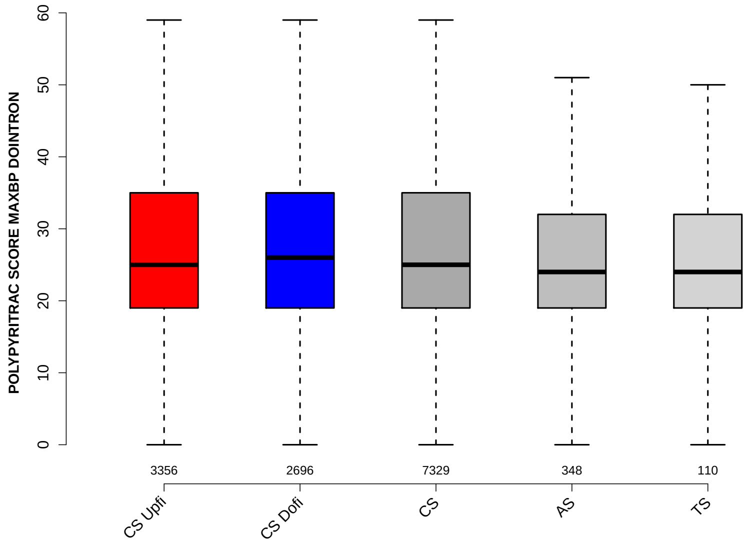

Significant results from Mann-Whitney U test:

- none

#### 6.50 BPSCORE MAXBP DOINTRON

Back to: Overview | ToC

Meaning: SVM classification score of best BP

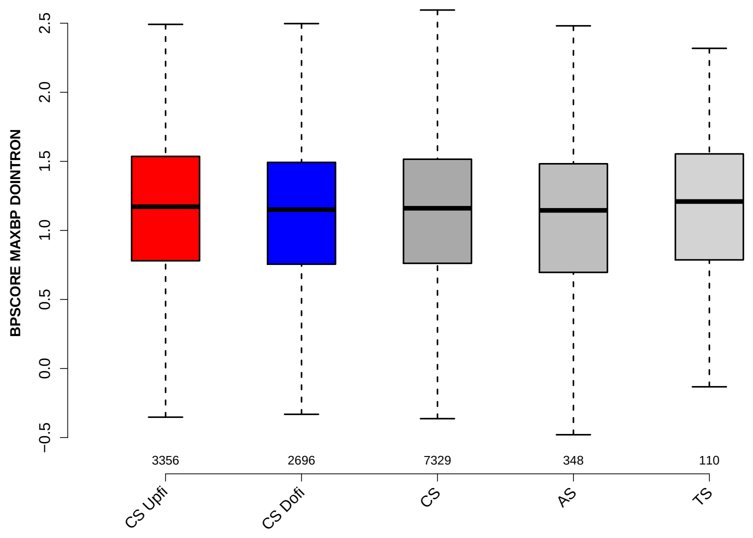

Significant results from Mann-Whitney U test:

- none

#### 6.51 NUM PREDICTED BPS DOINTRON

Back to: Overview | ToC

Meaning: number of all predicted BPs which have a positive BP score

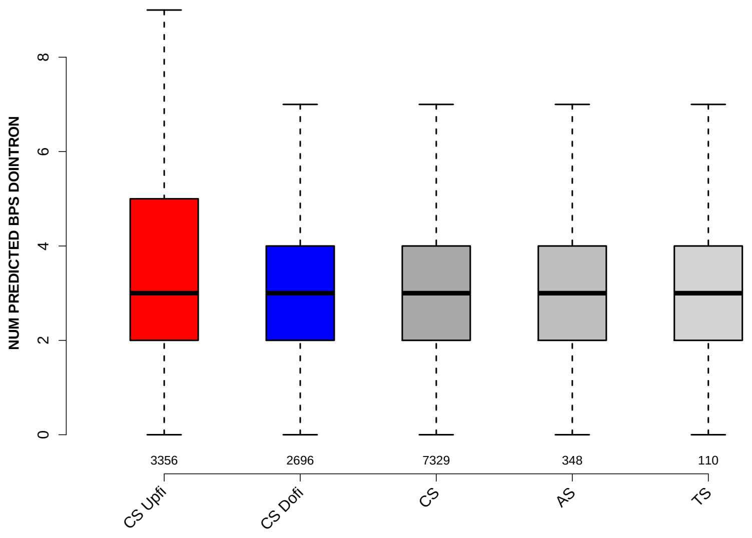

Significant results from Mann-Whitney U test:

- `CS_Upfi` vs `CS_Dofi` : 2.90865e-06
  mean: 3.3167 > 3.1142 , median: 3 = 3
- `CS_Dofi` vs `CS` : 0.000279654
  mean: 3.1142 < 3.2588 , median: 3 = 3
- `CS_Dofi` vs `AS` : 0.0465397
  mean: 3.1142 < 3.2586 , median: 3 = 3

#### 6.52 MEDIAN DIST FROM BP TO 3SS DOINTRON

Back to: Overview | ToC

Meaning: like DIST FROM MAXBP TO 3SS but median over top-3 predicted BPs

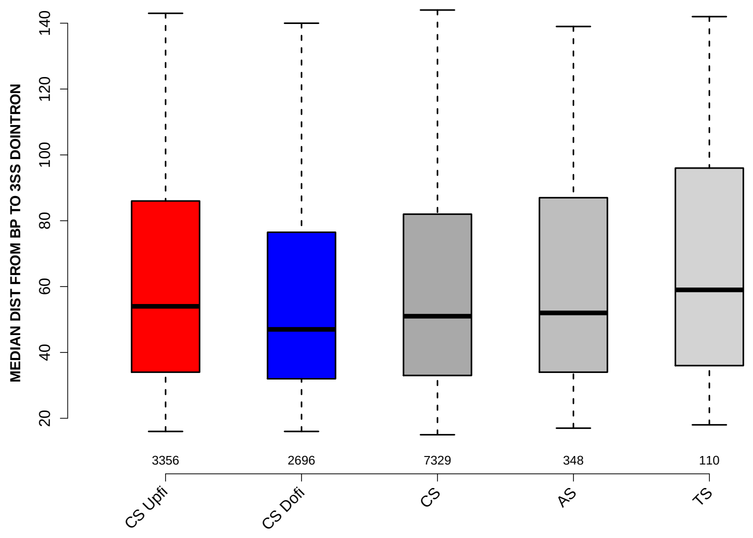

Significant results from Mann-Whitney U test:

- `CS_Upfi` vs `CS_Dofi` : 5.78944e-10
  mean: 61.7901 > 56.5757 , median: 54 > 47
- `CS_Upfi` vs `CS` : 0.000664006
  mean: 61.7901 > 59.4628 , median: 54 > 51
- `CS_Dofi` vs `CS` : 6.89389e-05
  mean: 56.5757 < 59.4628 , median: 47 < 51
- `CS_Dofi` vs `AS` : 0.0363436
  mean: 56.5757 < 60.3175 , median: 47 < 52
- `CS_Dofi` vs `TS` : 0.00654598
  mean: 56.5757 < 66.0773 , median: 47 < 59

#### 6.53 MEDIAN SCORE FOR BPSEQ DOINTRON

Back to: Overview | ToC

Meaning: like SCORE FOR MAXBP SEQ but median over top-3 predicted BPs

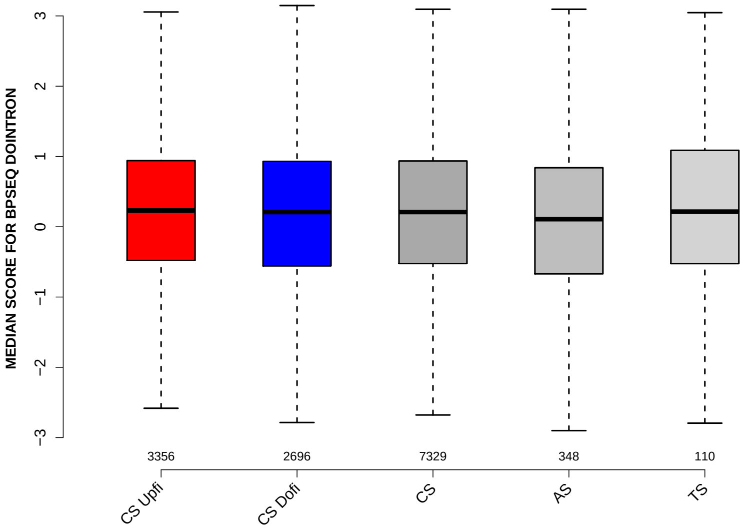

Significant results from Mann-Whitney U test:
- none

#### 6.54 MEDIAN PYRIMIDINECONT DOINTRON

Back to: Overview | ToC

Meaning: like PYRIMIDINECONT MAXBP but median over top-3 predicted BPs

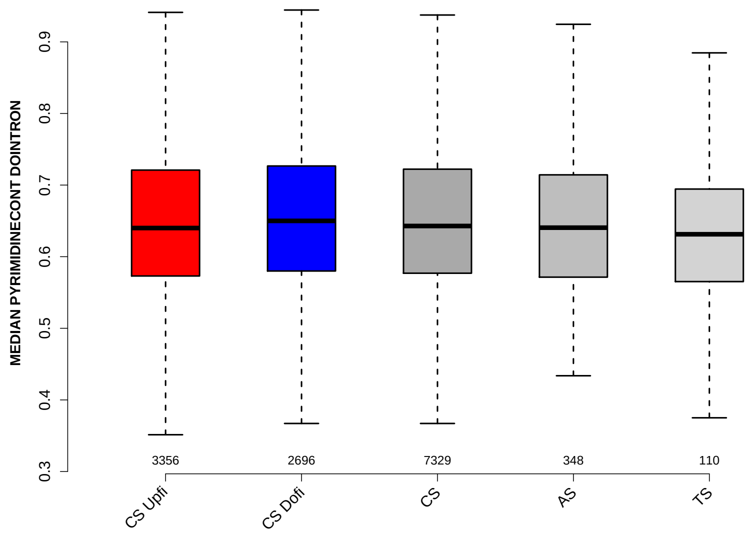

Significant results from Mann-Whitney U test:

- CS_Upfi vs CS_Dofi : 0.0122787
  mean: 0.649231 < 0.65505 , median: 0.64 < 0.65
- CS_Dofi vs AS : 0.0473992
  mean: 0.65505 > 0.642944 , median: 0.65 > 0.640513

#### 6.55 MEDIAN POLYPYRITRAC OFFSET DOINTRON

Back to: Overview | ToC

Meaning: like POLYPYRITRAC OFFSET MAXBP but median over top-3 predicted BPs

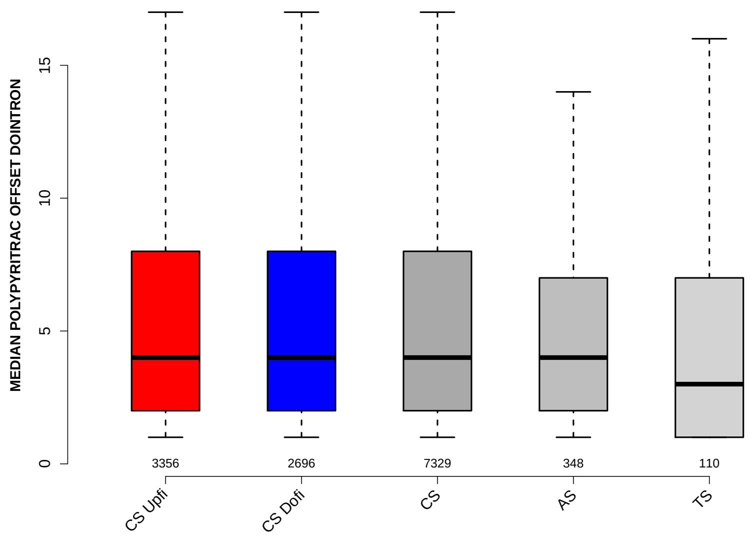

Significant results from Mann-Whitney U test:

- none

#### 6.56 MEDIAN POLYPYRITRAC LEN DOINTRON

Back to: Overview | ToC

Meaning: like POLYPYRITRAC LEN MAXBP but median over top-3 predicted BPs

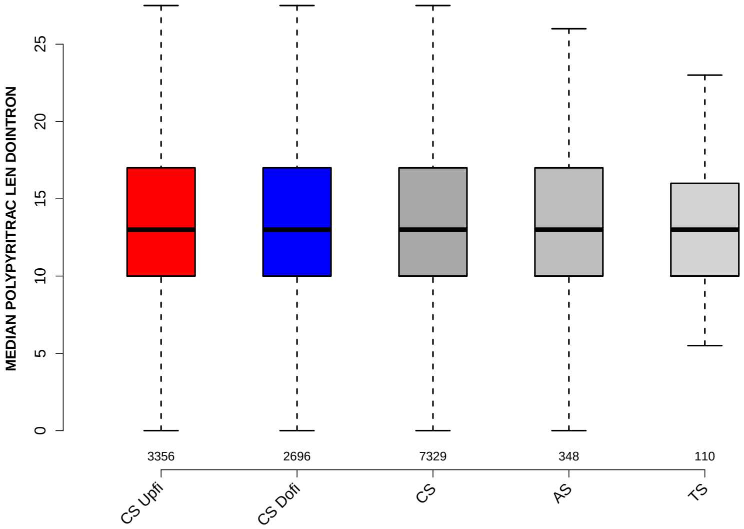

Significant results from Mann-Whitney U test:

- none

#### 6.57 MEDIAN POLYPYRITRAC SCORE DOINTRON

Back to: Overview | ToC

Meaning: like POLYPYRITRAC SCORE MAXBP but median over top-3 predicted BPs

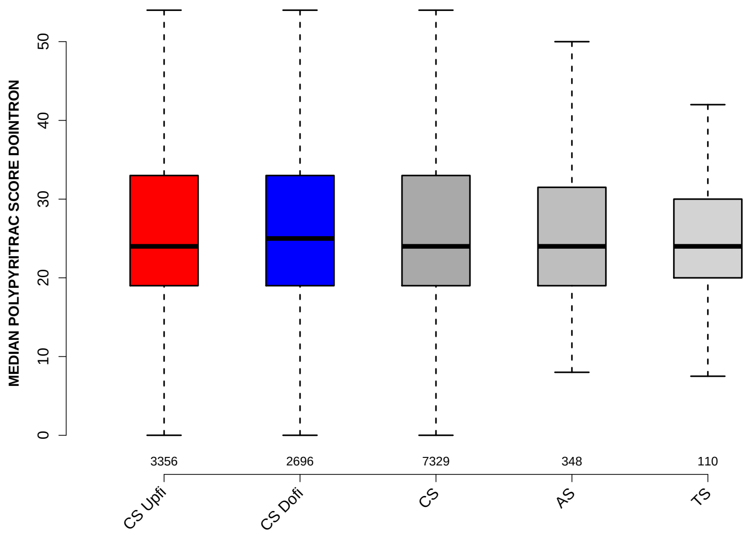

Significant results from Mann-Whitney U test:

- none

#### 6.58 MEDIAN BPSCORE DOINTRON

Back to: Overview | ToC

Meaning: like BPSCORE MAXBP but median over top-3 predicted BPs

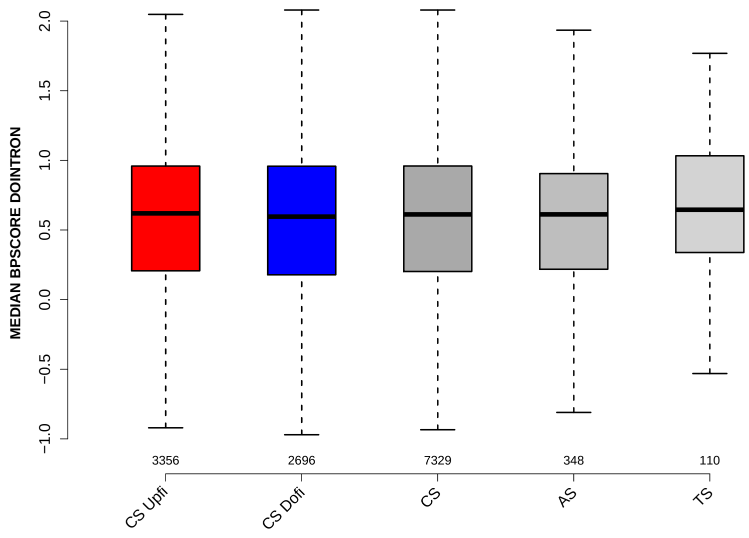

Significant results from Mann-Whitney U test:

- none

#### 6.59 MEDIAN TR LENGTH

Back to: Overview | ToC

Meaning: median length of transcripts the exon occurs in

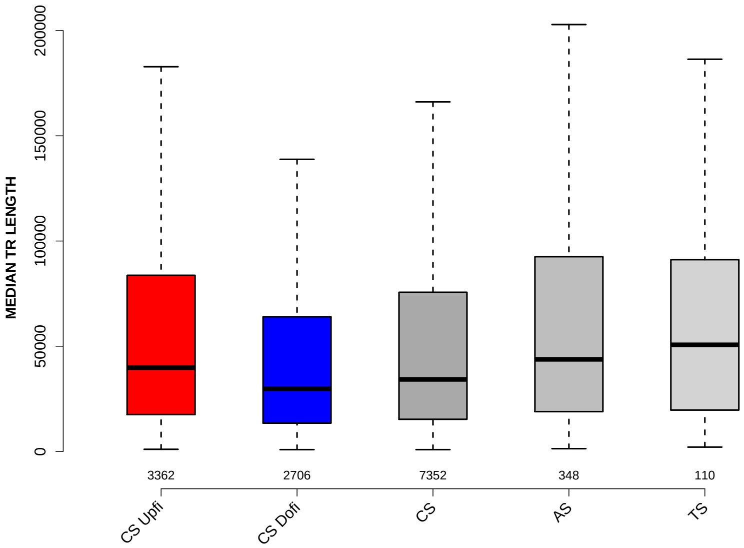

Significant results from Mann-Whitney U test:

- CS_Upfi vs CS_Dofi : 6.27236e-20
  mean: 70108.3841 > 51070.179 , median: 39779.5 > 29721.5
- CS_Upfi vs CS : 1.62006e-07
  mean: 70108.3841 > 60642.9765 , median: 39779.5 > 34234.75
- CS_Dofi vs CS : 1.36596e-08
  mean: 51070.179 < 60642.9765 , median: 29721.5 < 34234.75
- CS_Dofi vs AS : 2.84542e-07
  mean: 51070.179 < 74006.7902 , median: 29721.5 < 43767.25
- CS_Dofi vs TS : 0.00309992
  mean: 51070.179 < 84014.9136 , median: 29721.5 < 50657
- CS vs AS : 0.0023907
  mean: 60642.9765 < 74006.7902 , median: 34234.75 < 43767.25

#### 6.60 MEDIAN EXON NUMBER

Back to: Overview | ToC

Meaning: ... of transcripts where exon was found in

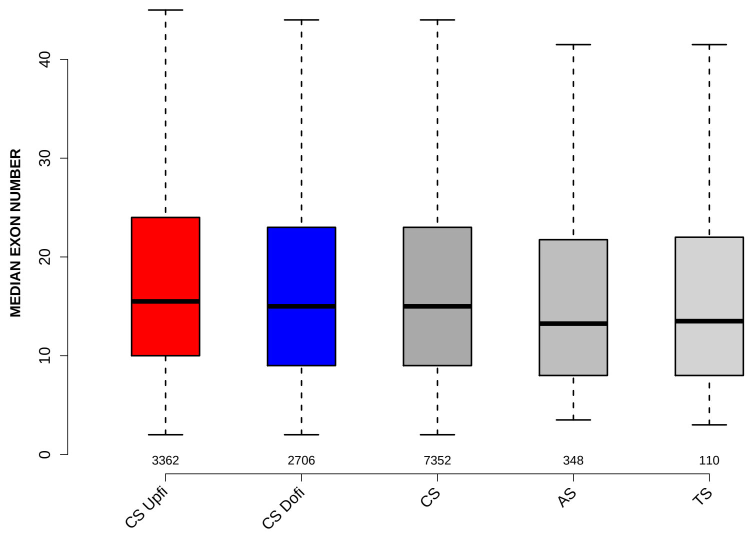

Significant results from Mann-Whitney U test:

- CS_Upfi vs CS_Dofi : 0.0248345
  mean: 19.0257 > 18.4189 , median: 15.5 > 15
- CS_Upfi vs AS : 0.00253283
  mean: 19.0257 > 18.342 , median: 15.5 > 13.25
- CS_Upfi vs TS : 0.0245199
  mean: 19.0257 > 16.0455 , median: 15.5 > 13.5
- CS_Dofi vs AS : 0.0489439
  mean: 18.4189 > 18.342 , median: 15 > 13.25
- CS vs AS : 0.0123761
  mean: 18.6219 > 18.342 , median: 15 > 13.25

#### 6.61 EXON MEDIANRELATIVERANK

Back to: Overview | ToC

Meaning: relative rank = rank / number of all exons in transcript, is between 0 and 1

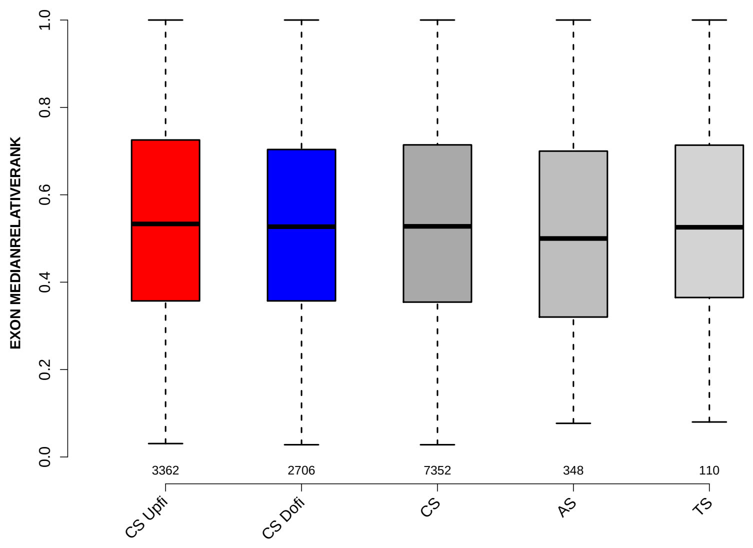

Significant results from Mann-Whitney U test:

- `CS_Upfi` vs `AS` : 0.0203689
  mean: 0.538358 > 0.508445 , median: 0.533333 > 0.5
- `CS` vs `AS` : 0.0421525
  mean: 0.533369 > 0.508445 , median: 0.527778 > 0.5

#### 6.62 EXON MEDIANRELATIVERANK 3BINS

Back to: Overview | ToC

Meaning: median bin into which EXON MEDIANRELATIVERANK falls when binning 0-1 into 3 bins

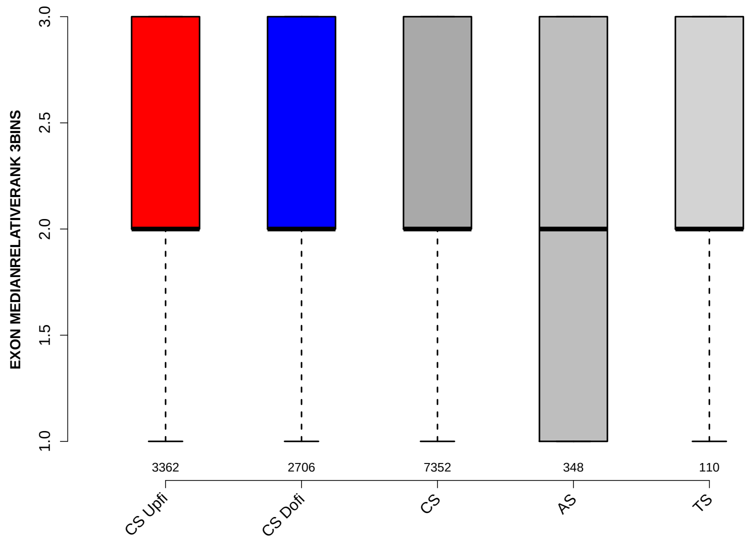

Significant results from Mann-Whitney U test:

- `CS_Upfi` vs `AS` : 0.0242383
  mean: 2.1341 > 2.0402 , median: 2 = 2

#### 6.63 EXON MEDIANRELATIVERANK 5BINS

Back to: Overview | ToC

Meaning: similar to EXON MEDIANRELATIVERANK 3BINS with 5 bins

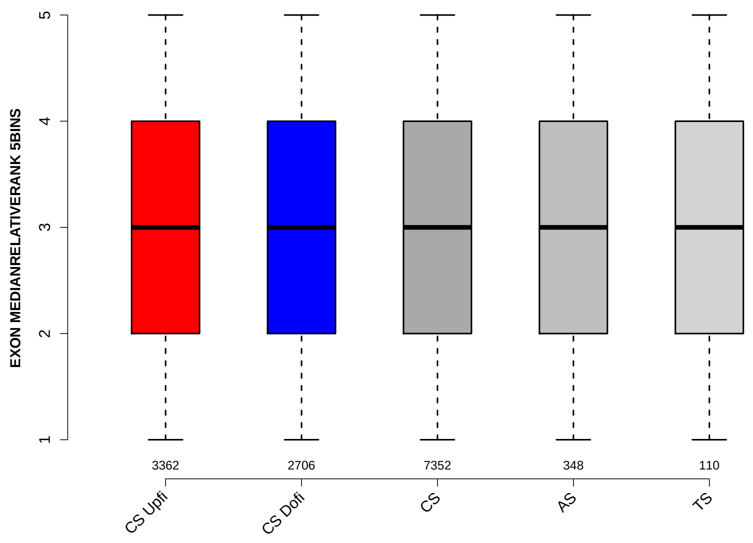

Significant results from Mann-Whitney U test:

- `CS_Upfi` vs `AS` : 0.0186074
  mean: 3.1936 > 3.0374 , median: 3 = 3
- `CS` vs `AS` : 0.0412976
  mean: 3.1708 > 3.0374 , median: 3 = 3

#### 6.64 EXON MEDIANRELATIVERANK 10BINS

Back to: Overview | ToC

Meaning: similar to EXON MEDIANRELATIVERANK 3BINS with 10 bins

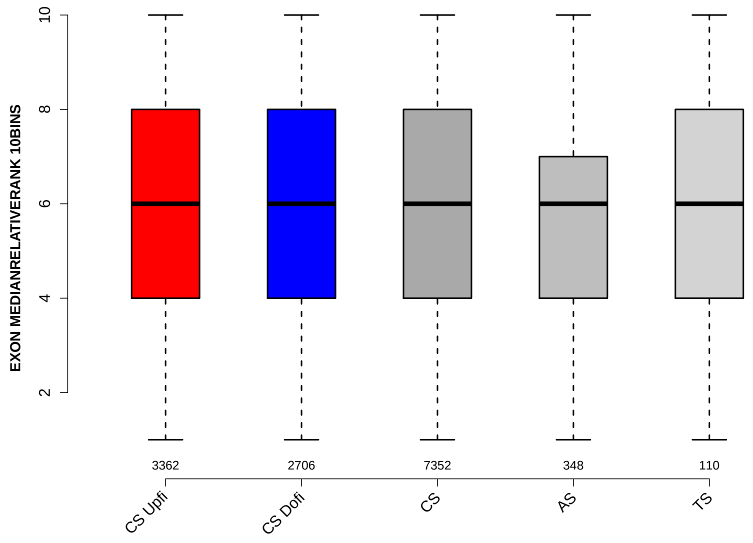

Significant results from Mann-Whitney U test:

- `CS_Upfi` vs `AS` : 0.0271396
  mean: 5.9179 > 5.6236 , median: 6 = 6

#### 6.65 NTRS ALL FOR GENE

Back to: Overview | ToC

Meaning: number of transcripts of gene where the exon was found in

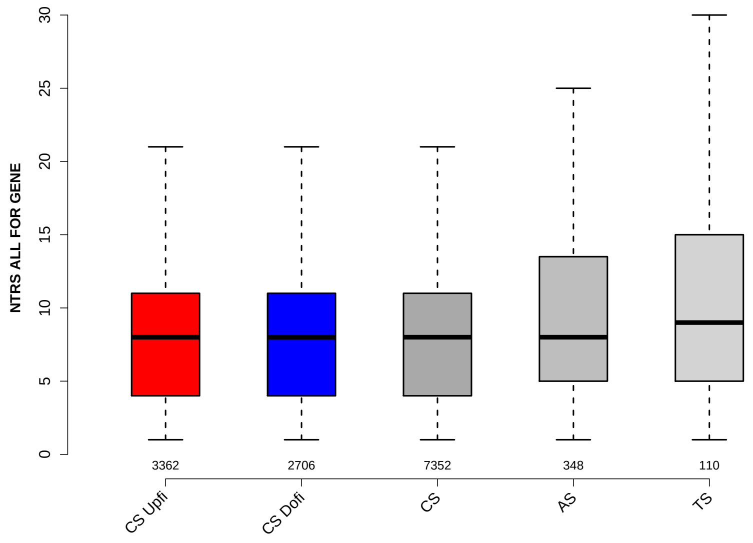

Significant results from Mann-Whitney U test:

- CS_Upfi vs AS : 0.00116423
  mean: 8.4783 < 9.6322 , median: 8 = 8
- CS_Upfi vs TS : 0.0015787
  mean: 8.4783 < 10.8364 , median: 8 < 9
- CS_Dofi vs AS : 0.00138433
  mean: 8.486 < 9.6322 , median: 8 = 8
- CS_Dofi vs TS : 0.00174896
  mean: 8.486 < 10.8364 , median: 8 < 9
- CS vs AS : 0.0013663
  mean: 8.5394 < 9.6322 , median: 8 = 8
- CS vs TS : 0.00183387
  mean: 8.5394 < 10.8364 , median: 8 < 9

#### 6.66 PROP FIRST EXON

Back to: Overview | ToC

Meaning: NTRS WITH EXON AS FIRST EXON / NTRS WITH EXON

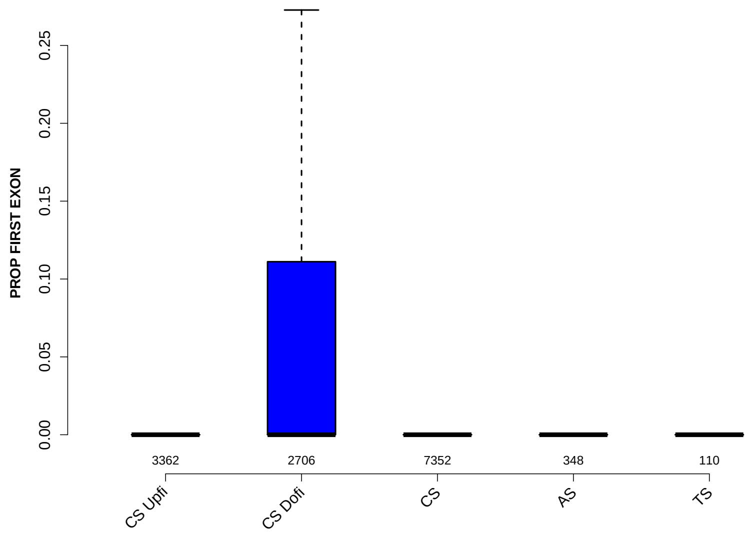

Significant results from Mann-Whitney U test:

- CS_Upfi vs CS_Dofi : 1.0432e-65
  mean: $0.0224109 < 0.0685492$ , median: $0 = 0$
- CS_Upfi vs CS : 2.29962e-24
  mean: $0.0224109 < 0.0429351$ , median: $0 = 0$
- CS_Upfi vs AS : 0.000258802
  mean: $0.0224109 < 0.0423092$ , median: $0 = 0$
- CS_Upfi vs TS : 0.0225434
  mean: $0.0224109 < 0.0273017$ , median: $0 = 0$
- CS_Dofi vs CS : 1.67316e-24
  mean: $0.0685492 > 0.0429351$ , median: $0 = 0$
- CS_Dofi vs AS : 3.44425e-05
  mean: $0.0685492 > 0.0423092$ , median: $0 = 0$
- CS_Dofi vs TS : 0.00887162
  mean: $0.0685492 > 0.0273017$ , median: $0 = 0$

#### 6.67 PROP LAST EXON

Back to: Overview | ToC

Meaning: NTRS WITH EXON AS LAST EXON / NTRS WITH EXON

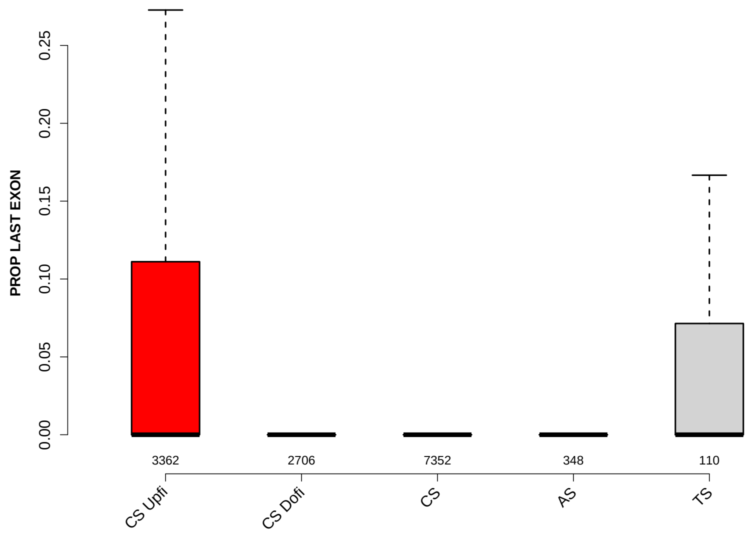

Significant results from Mann-Whitney U test:

- CS_Upfi vs CS_Dofi : 3.52035e-35
  mean: 0.0694444 > 0.0320389 , median: 0 = 0
- CS_Upfi vs CS : 3.54822e-11
  mean: 0.0694444 > 0.0524761 , median: 0 = 0
- CS_Upfi vs AS : 0.000610512
  mean: 0.0694444 > 0.0432578 , median: 0 = 0
- CS_Dofi vs CS : 2.23098e-16
  mean: 0.0320389 < 0.0524761 , median: 0 = 0
- CS_Dofi vs AS : 0.019616
  mean: 0.0320389 < 0.0432578 , median: 0 = 0
- CS_Dofi vs TS : 0.00128866
  mean: 0.0320389 < 0.0566585 , median: 0 = 0

#### 6.68 PROP INTERNAL EXON

Back to: Overview | ToC

Meaning: NTRS WITH EXON AS INTERNAL EXON / NTRS WITH EXON

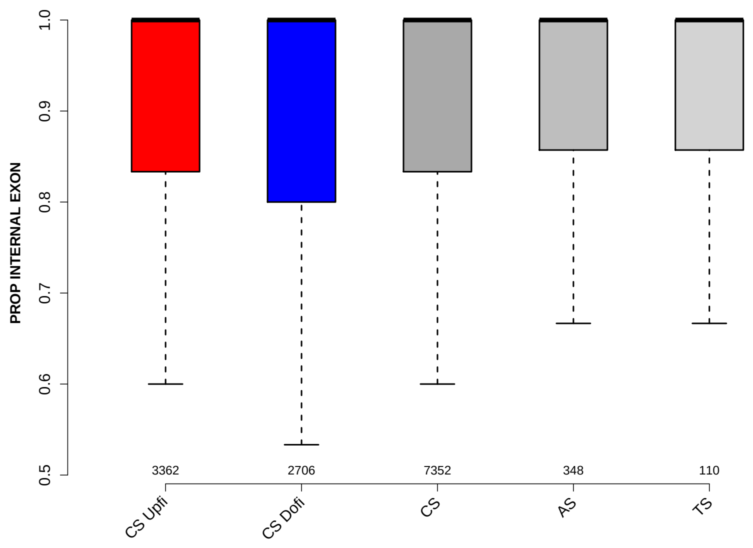

Significant results from Mann-Whitney U test:

- `CS_Upfi` vs `CS_Dofi` : 0.0164461
  mean: 0.910721 > 0.901994 , median: 1 = 1

#### 6.69 PROP EXON IN UTR

Back to: Overview | ToC

Meaning: NTRS WITH EXON IN UTR / NTRS WITH EXON

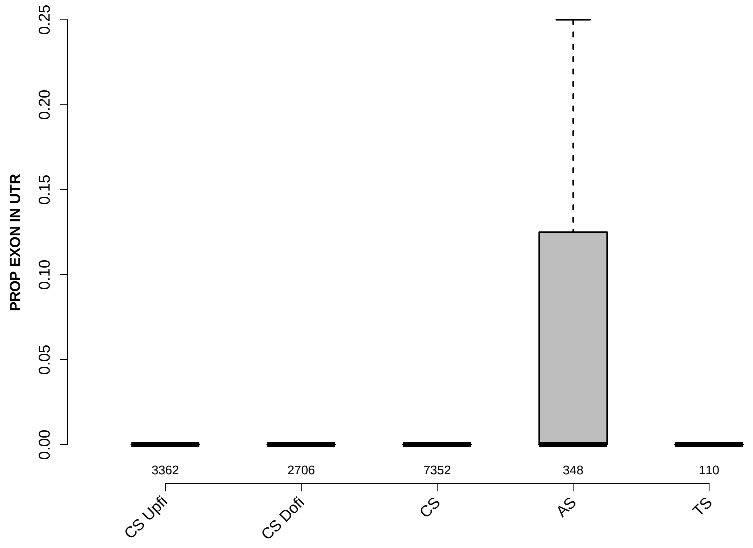

Significant results from Mann-Whitney U test:

- CS_Upfi vs CS_Dofi : 6.17809e-06
  mean: 0.0572499 < 0.0761719 , median: 0 = 0
- CS_Upfi vs CS : 0.0025724
  mean: 0.0572499 < 0.0661572 , median: 0 = 0
- CS_Upfi vs AS : 8.44723e-07
  mean: 0.0572499 < 0.0855124 , median: 0 = 0
- CS_Dofi vs CS : 0.01681
  mean: 0.0761719 > 0.0661572 , median: 0 = 0
- CS_Dofi vs AS : 0.0146454
  mean: 0.0761719 < 0.0855124 , median: 0 = 0
- CS vs AS : 0.000209062
  mean: 0.0661572 < 0.0855124 , median: 0 = 0
