## Supplementary File 3 for "*Insplico*: Effective computational tool for studying intron splicing order genome-wide with short and long RNA-seq reads"

### Comparison of exons grouped into: ctr, srrm4-upfi, srrm4-dofi

June 10, 2022
Matt version 1.3.0

#### Contents

1 Infos 4

2 Warning: Please read this note carefully 4

3 Notes for publishing results 4

4 Data sets 5

5 Overview: Features with statistically significant differences (p-val $\leq 0.05$) 6

### 4 Data sets

Input file:
`srrm4_exons_dpsi15-MI-v2.tab`
Selection criteria for defining exons groups:
`ctr` : having value `ctr` in column `GROUP`
`srrm4_upfi` : having value `srrm4_upfi` in column `GROUP`
`srrm4_dofi` : having value `srrm4_dofi` in column `GROUP`
Exon duplicates removal: yes
Numbers of exons per group before / after neglecting exons which were not found in GTF file (gene annotation). For the comparisons only exons which were found in the gene annotation are used. These numbers might change slightly for each feature if NAs occur.
`ctr`: 120 / 115
`srrm4_upfi`: 24 / 24
`srrm4_dofi`: 55 / 55

### 5 Overview: Features with statistically significant differences (p-val $\leq 0.05$)

#### MAXENTSCR HSAMODEL 3SS

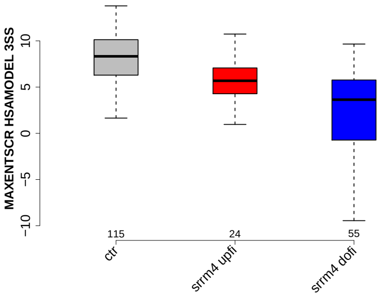

#### MEDIAN TR LENGTH

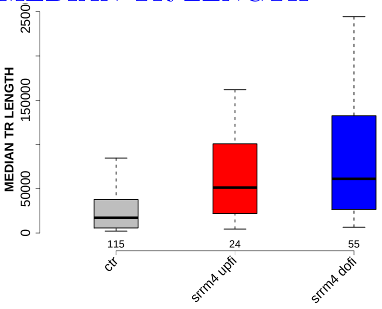

#### EXON LENGTH

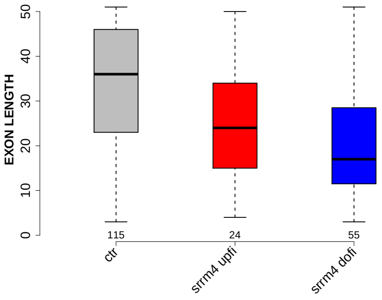

#### RATIO UPINTRON EXON LENGTH

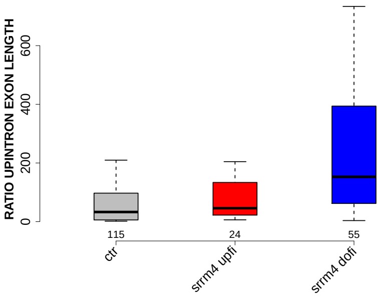

#### UPINTRON GCC

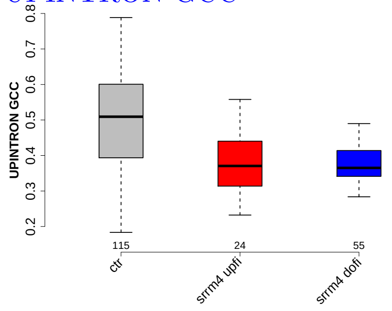

#### RATIO DOEXON EXON LENGTH

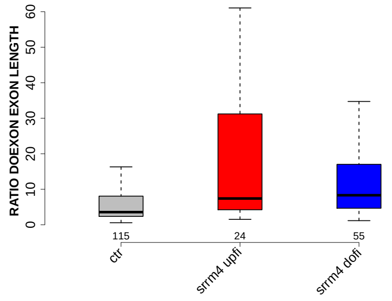

#### SF1 HIGHESTSCORE 3SS UPINTRON

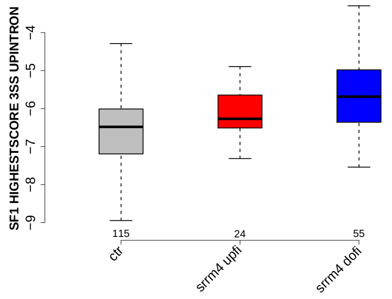

#### RATIO UPEXON EXON LENGTH

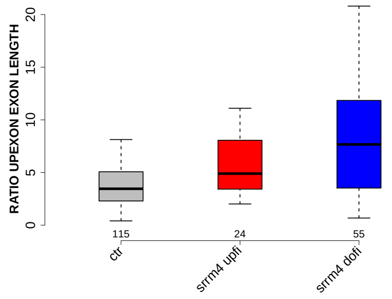

#### PROP INTERNAL EXON

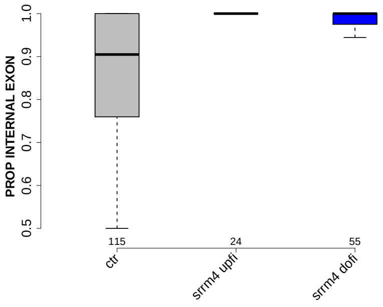

#### PROP EXON IN UTR

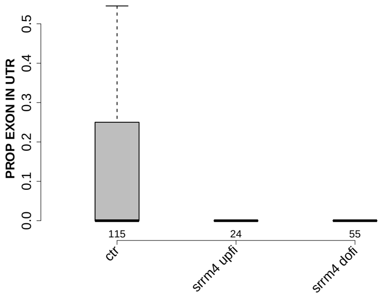

#### MEDIAN EXON NUMBER

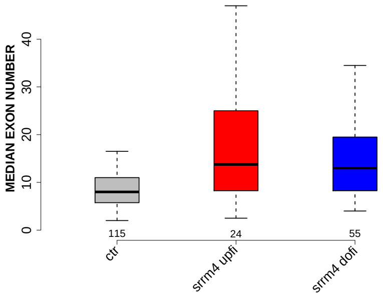

#### DOINTRON GCC

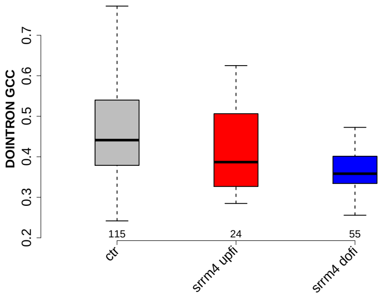

#### UPEXON GCC

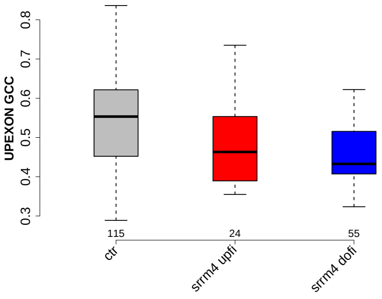

#### MEDIAN POLYPYRITRAC SCORE UPINTRON

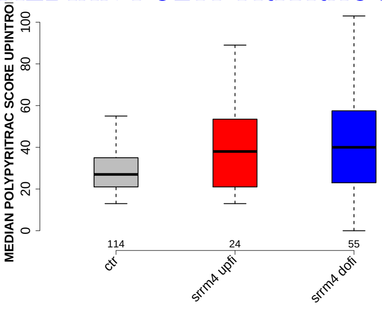

#### UPINTRON MEDIANLENGTH

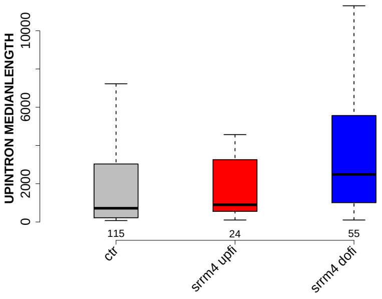

#### UP 5SS 20INT10EX GCC

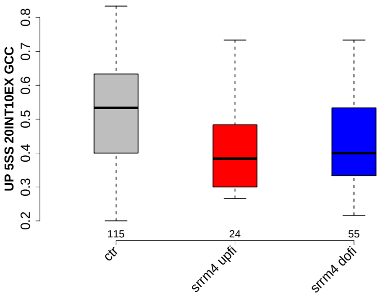

#### MEDIAN POLYPYRITRAC OFFSET UPINTRON

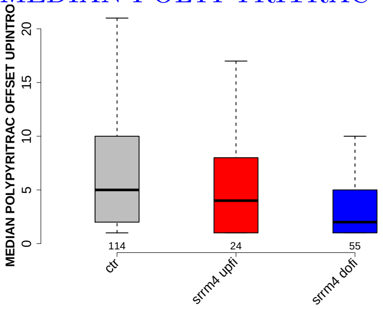

#### PROP LAST EXON

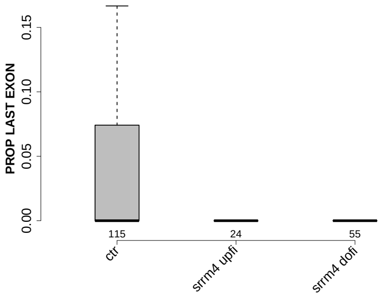

#### NUM PREDICTED BPS UPINTRON

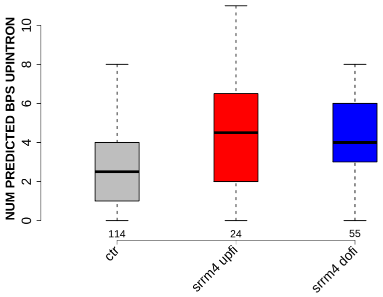

#### GCC 5SS 20INT10EX

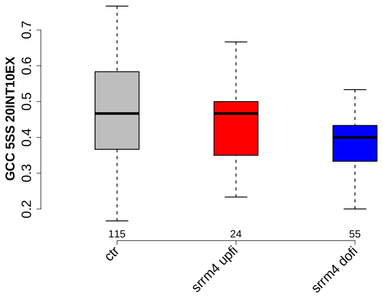

#### MEDIAN POLYPYRITRAC LEN UPINTRON

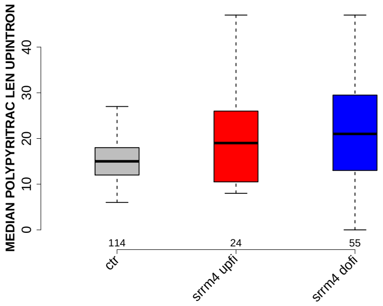

#### DOEXON GCC

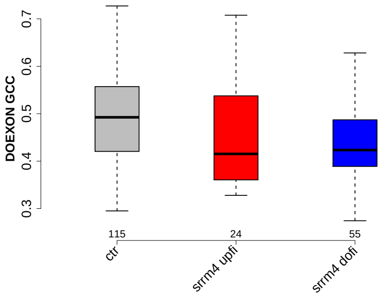

#### PROP FIRST EXON

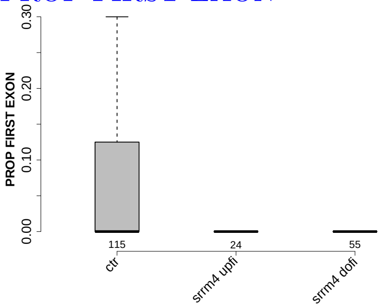

#### RATIO UPINTRON EXON GCC

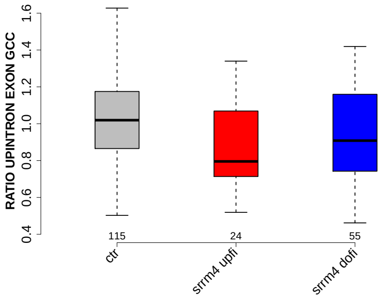

#### EXON MEDIANRELATIVERANK 5BINS

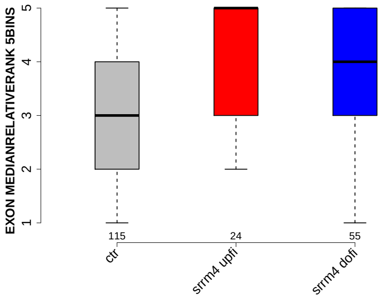

#### EXON MEDIANRELATIVERANK 10BINS

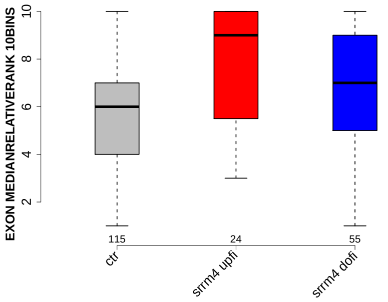

#### EXON MEDIANRELATIVERANK

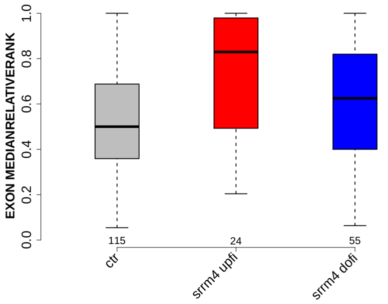

#### POLYPYRITRAC LEN MAXBP UPINTRON

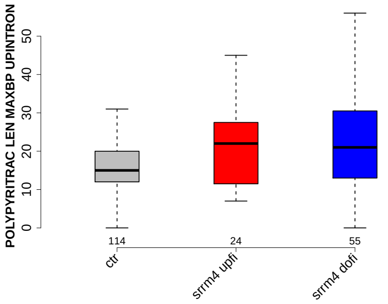

#### POLYPYRITRAC SCORE MAXBP UPINTRON

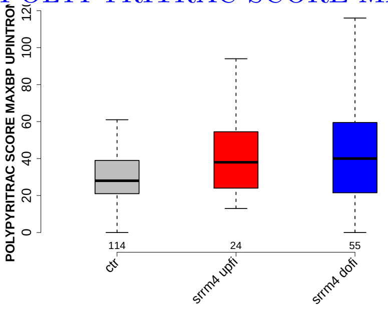

#### MAXENTSCR HSAMODEL 5SS

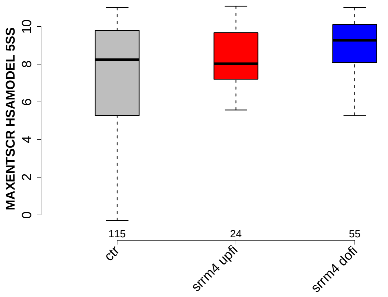

#### MAXENTSCR HSAMODEL UPSTRM 5SS

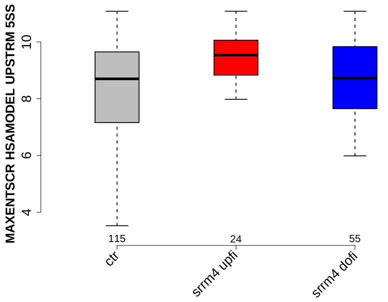

#### MEDIAN PYRIMIDINECONT UPINTRON

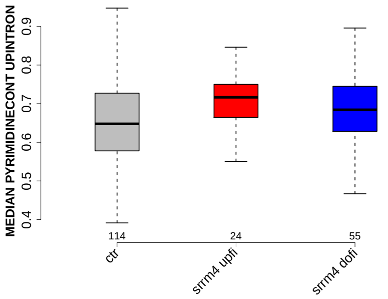

#### RATIO DOINTRON EXON LENGTH

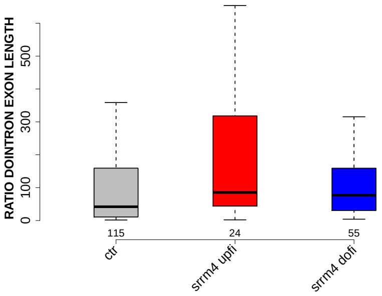

#### POLYPYRITRAC OFFSET MAXBP UPINTRON

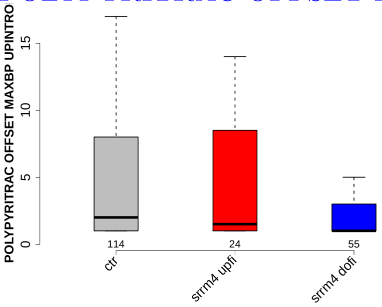

#### EXON GCC

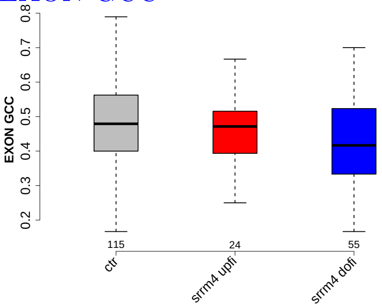

#### NUM PREDICTED BPS DOINTRON

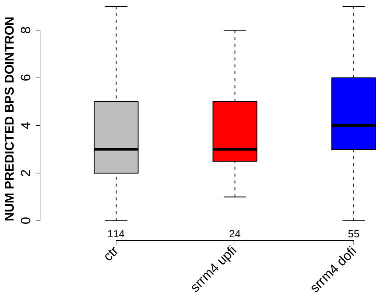

#### BPSCORE MAXBP UPINTRON

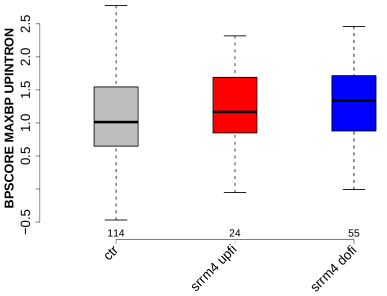

#### EXON MEDIANRELATIVERANK 3BINS

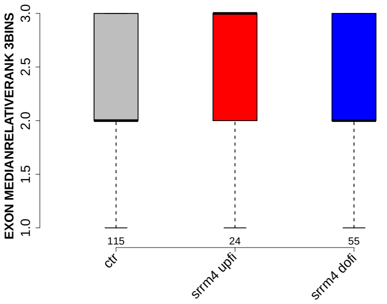

#### MEDIAN BPSCORE UPINTRON

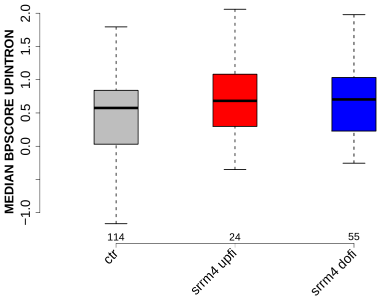

#### DOEXON MEDIANLENGTH

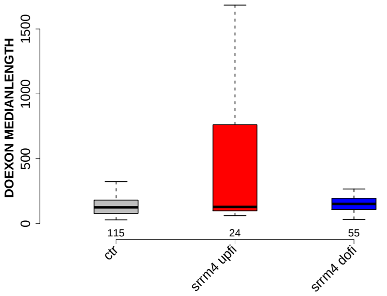

### DOINTRON MEDIANLENGTH

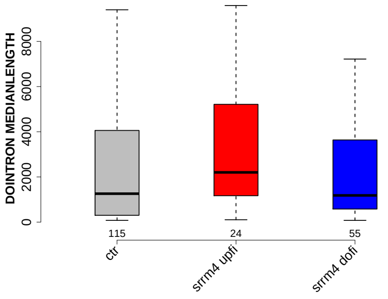

### MEDIAN BPSCORE DOINTRON

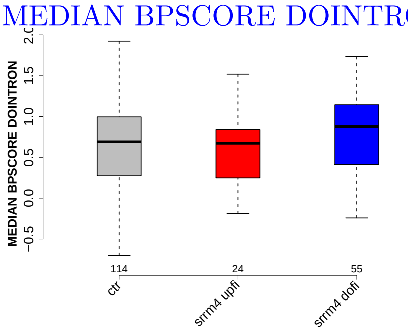

### 6 Details: Box plots and statistical assessments for all features

#### 6.1 EXON LENGTH

Back to: Overview | ToC

Meaning:

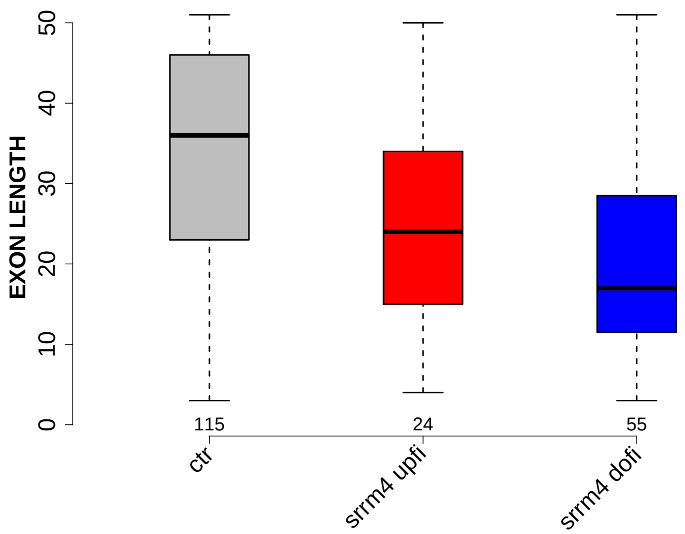

Significant results from Mann-Whitney U test:

- `ctr` vs `srrm4_upfi` : 0.0159894
  mean: 33.1304 > 25.3333 , median: 36 > 24
- `ctr` vs `srrm4_dofi` : 2.8587e-07
  mean: 33.1304 > 20.5091 , median: 36 > 17

#### 6.2 UPEXON MEDIANLENGTH

Back to: Overview | ToC

Meaning: median length of up-stream exon

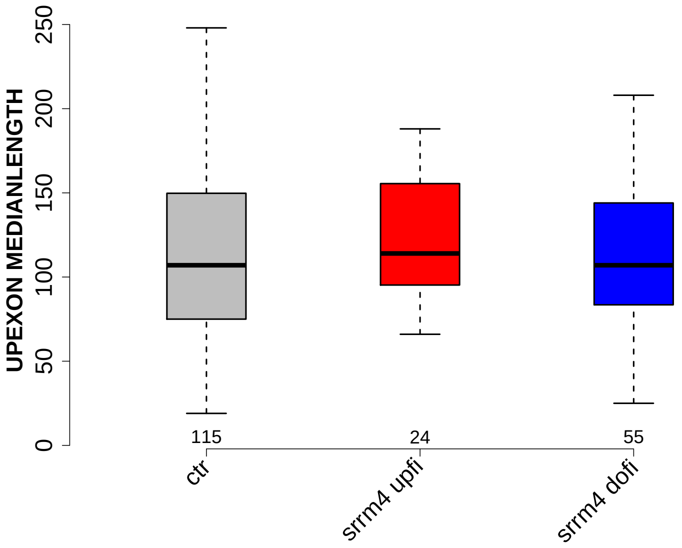

Significant results from Mann-Whitney U test:

- none

##### 6.3 DOEXON MEDIANLENGTH

Back to: Overview | ToC

Meaning: median length of down-stream exon

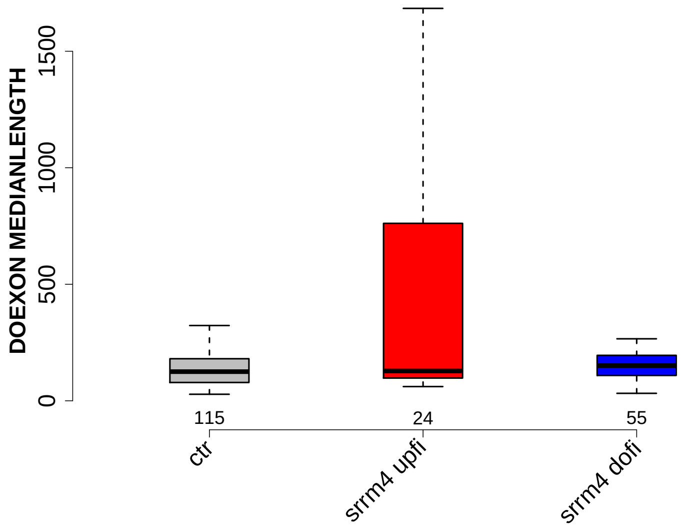

Significant results from Mann-Whitney U test:

- `ctr` vs `srrm4_dofi` : 0.0334315
  mean: 221.1174 > 209.1636 , median: 125 < 151

#### 6.4 RATIO UPEXON EXON LENGTH

Back to: Overview | ToC

Meaning: median up-stream exon length / exon length

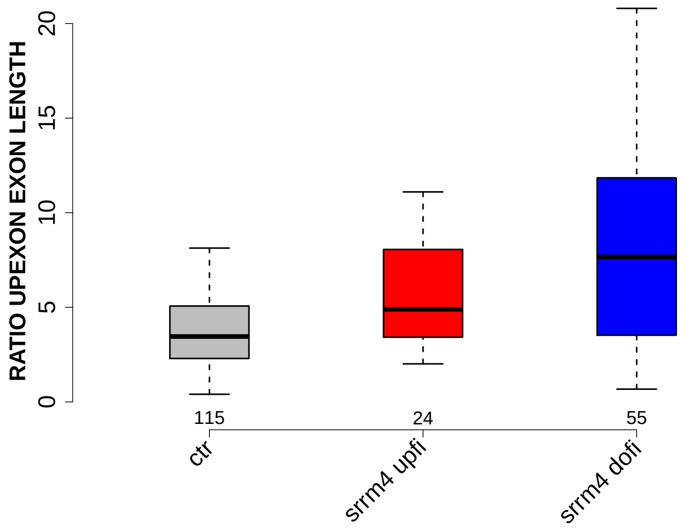

Significant results from Mann-Whitney U test:

- `ctr` vs `srrm4_upfi` : 0.00699159
  mean: 5.6095 < 7.4116 , median: 3.4565 < 4.8958
- `ctr` vs `srrm4_dofi` : 8.51637e-06
  mean: 5.6095 < 10.0838 , median: 3.4565 < 7.6667

#### 6.5 RATIO DOEXON EXON LENGTH

Back to: Overview | ToC

Meaning: median down-stream exon length / exon length

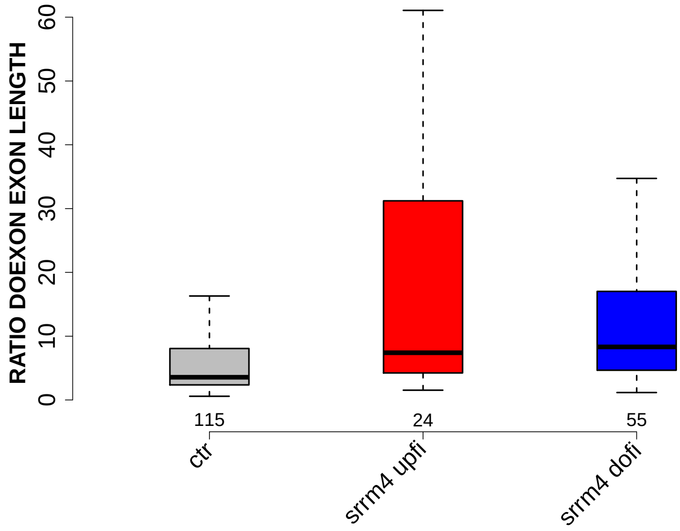

Significant results from Mann-Whitney U test:

- `ctr` vs `srrm4_upfi` : 0.00182094
  mean: 9.3354 < 39.2443 , median: 3.5625 < 7.4028
- `ctr` vs `srrm4_dofi` : 4.46799e-06
  mean: 9.3354 < 15.9763 , median: 3.5625 < 8.32

#### 6.6 UPINTRON MEDIANLENGTH

Back to: Overview | ToC

Meaning: median length of up-stream introns

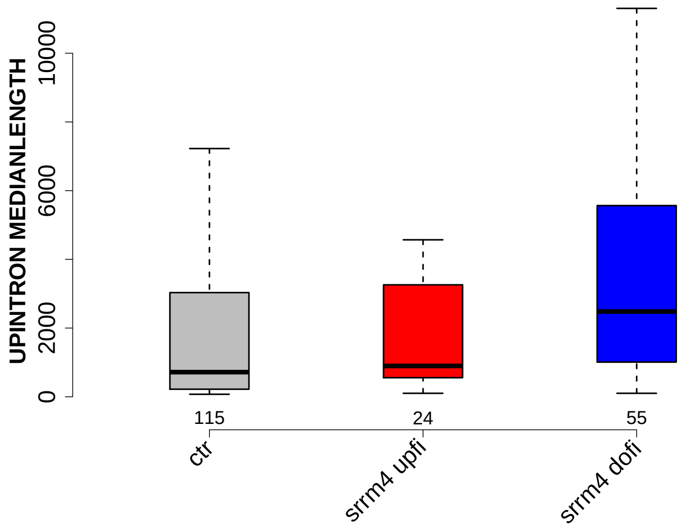

Significant results from Mann-Whitney U test:

- `ctr` vs `srrm4_dofi` : 0.000299529
  mean: 4293.1478 < 4827.0091 , median: 717 < 2482

#### 6.7 DOINTRON MEDIANLENGTH

Back to: Overview | ToC

Meaning: median length of down-stream introns

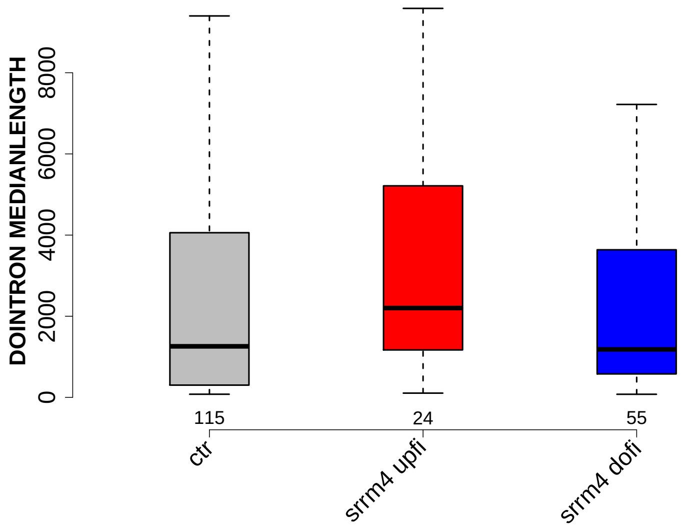

Significant results from Mann-Whitney U test:

- `ctr` vs `srrm4_upfi` : 0.0436588
  mean: 4657.3217 < 5738.625 , median: 1258 < 2201.5
- `srrm4_upfi` vs `srrm4_dofi` : 0.0479923
  mean: 5738.625 > 2441.4727 , median: 2201.5 > 1182

#### 6.8 RATIO UPINTRON EXON LENGTH

Back to: Overview | ToC

Meaning: median up-stream intron length / exon length

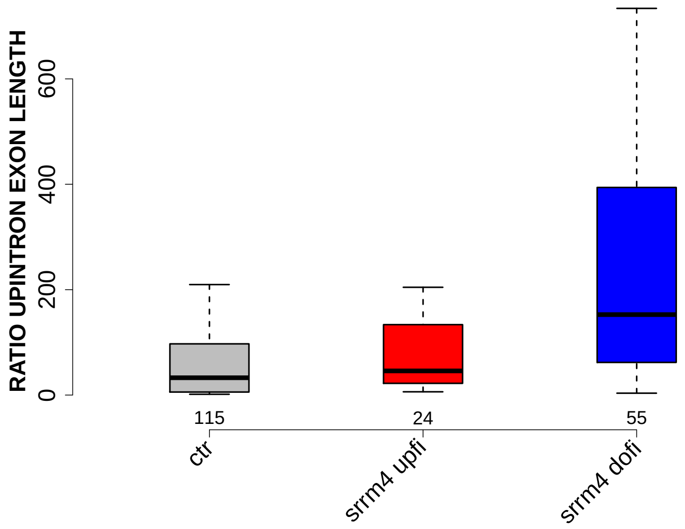

Significant results from Mann-Whitney U test:

- `ctr` vs `srrm4_dofi` : 7.1816e-07
  mean: 160.6581 < 289.6254 , median: 32.875 < 152.6667
- `srrm4_upfi` vs `srrm4_dofi` : 0.0116989
  mean: 217.7701 < 289.6254 , median: 45.8611 < 152.6667

#### 6.9 RATIO DOINTRON EXON LENGTH

Back to: Overview | ToC

Meaning: median down-stream intron length / exon length

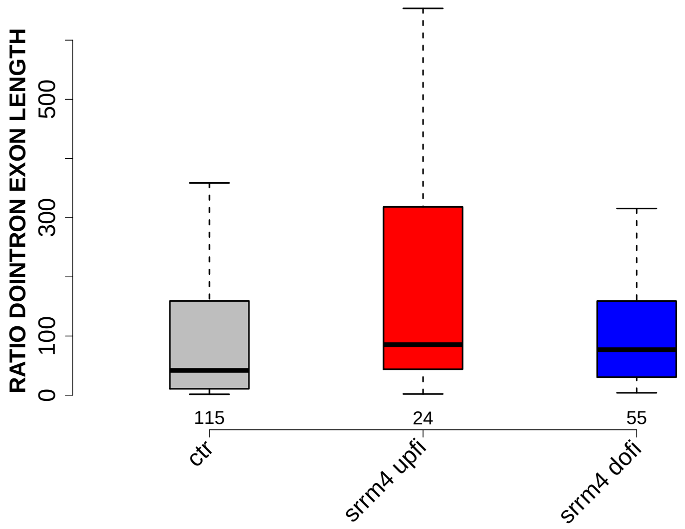

Significant results from Mann-Whitney U test:

- `ctr` vs `srrm4_upfi` : 0.011136
  mean: 233.2837 < 453.3498 , median: 41.9333 < 85.4111
- `ctr` vs `srrm4_dofi` : 0.0241319
  mean: 233.2837 > 170.4348 , median: 41.9333 < 76.9167

#### 6.10 EXON GCC

Back to: Overview | ToC

Meaning: GC content of entire exon sequence

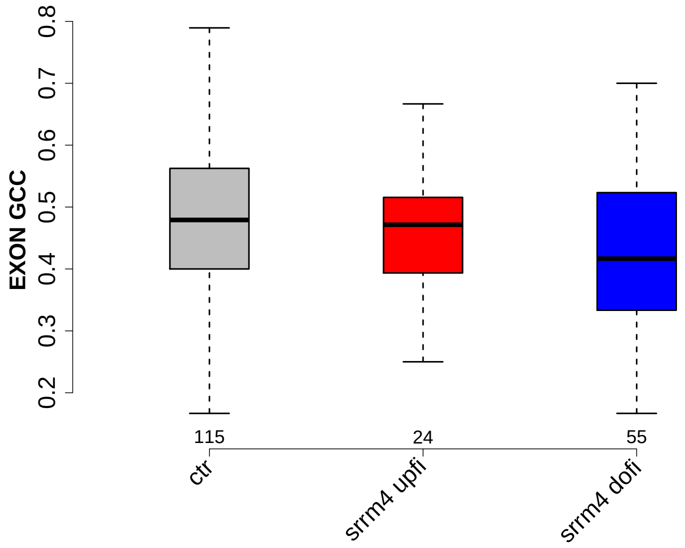

Significant results from Mann-Whitney U test:

- `ctr` vs `srrm4_dofi` : 0.0133733
  mean: 0.482338 > 0.433147 , median: 0.479167 > 0.416667

#### 6.11 UPINTRON GCC

Back to: Overview | ToC

Meaning: GC content of entire up-stream intron sequence

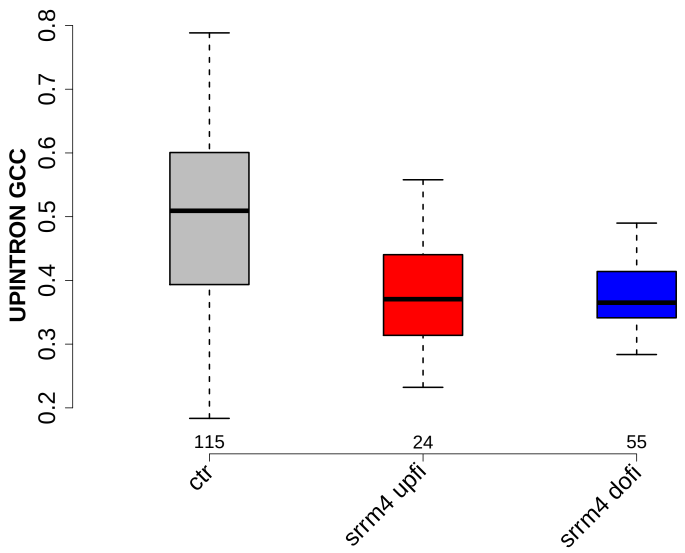

Significant results from Mann-Whitney U test:

- `ctr` vs `srrm4_upfi` : 0.000350309
  mean: 0.49662 > 0.393058 , median: 0.509219 > 0.370542
- `ctr` vs `srrm4_dofi` : 1.00954e-06
  mean: 0.49662 > 0.400146 , median: 0.509219 > 0.365022

#### 6.12 UPEXON GCC

Back to: Overview | ToC

Meaning: GC content of entire up-stream exon sequence

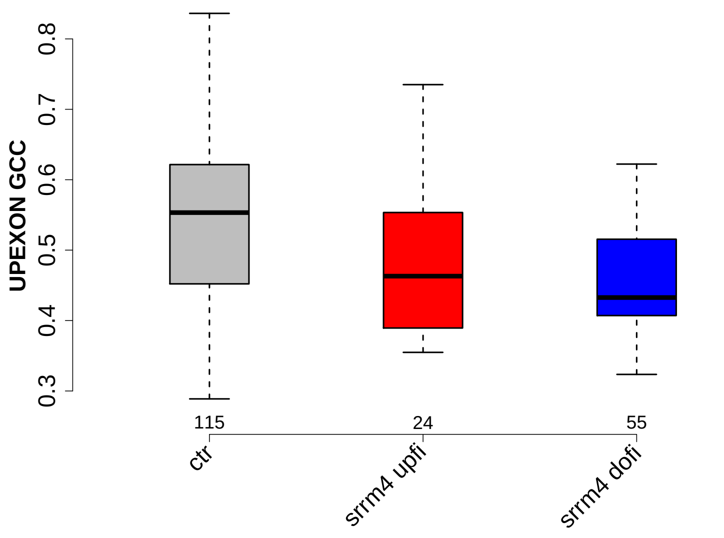

Significant results from Mann-Whitney U test:

- `ctr` vs `srrm4_upfi` : 0.0145383
  mean: 0.544426 > 0.48744 , median: 0.553299 > 0.463063
- `ctr` vs `srrm4_dofi` : 4.59343e-05
  mean: 0.544426 > 0.467907 , median: 0.553299 > 0.432692

#### 6.13 DOINTRON GCC

Back to: Overview | ToC

Meaning: GC content of entire down-stream intron sequence

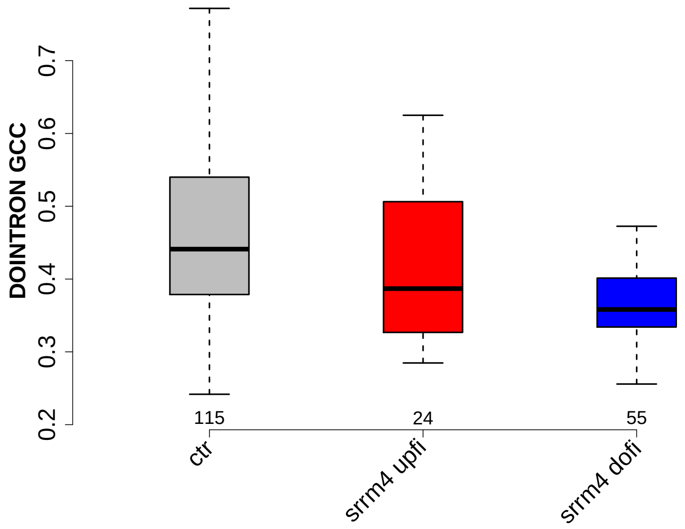

Significant results from Mann-Whitney U test:

- `ctr` vs `srrm4_upfi` : 0.0287194
  mean: 0.460786 > 0.410773 , median: 0.44111 > 0.386832
- `ctr` vs `srrm4_dofi` : 1.73228e-05
  mean: 0.460786 > 0.39575 , median: 0.44111 > 0.358217

#### 6.14 DOEXON GCC

Back to: Overview | ToC

Meaning: GC content of entire down-stream exon sequence

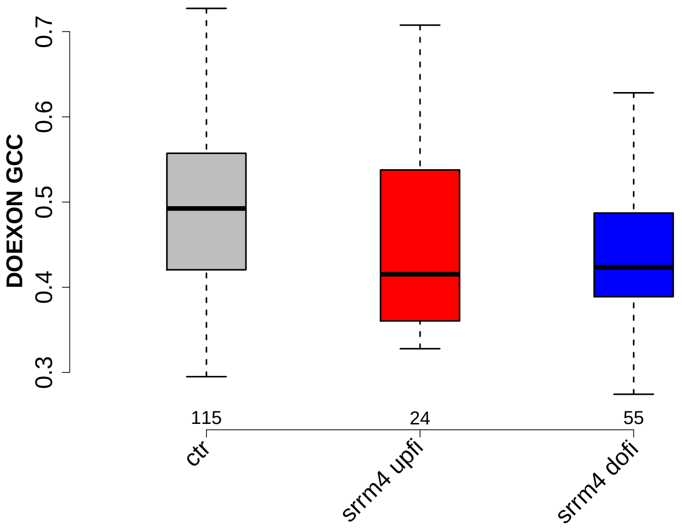

Significant results from Mann-Whitney U test:

- `ctr` vs `srrm4_dofi` : 0.00138524
  mean: 0.492421 > 0.443651 , median: 0.492453 > 0.423423

#### 6.15 RATIO UPEXON EXON GCC

Back to: Overview | ToC

Meaning: UPEXON GCC / EXON GCC

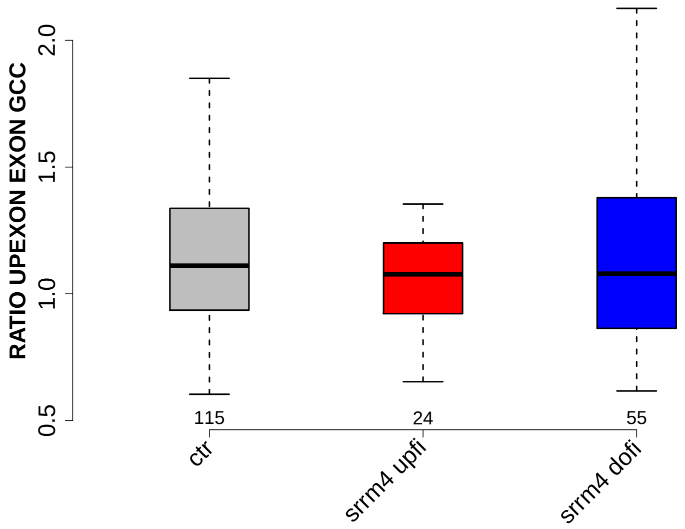

Significant results from Mann-Whitney U test:

- none

#### 6.16 RATIO UPINTRON EXON GCC

Back to: Overview | ToC

Meaning: UPINTRON GCC / EXON GCC

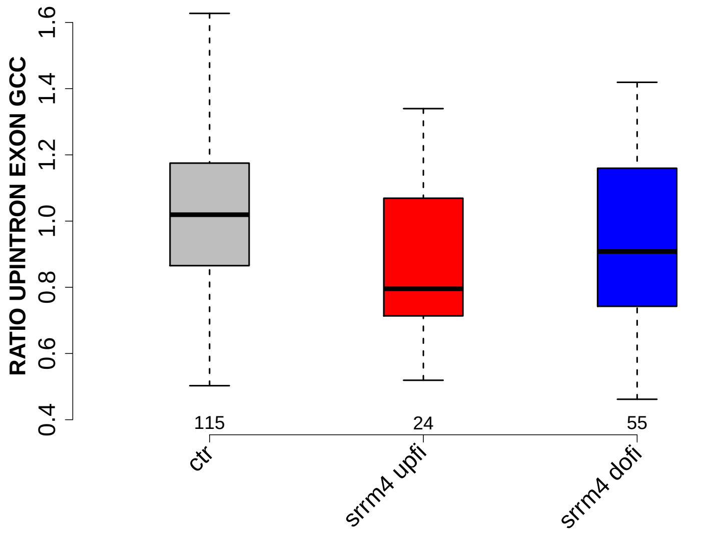

Significant results from Mann-Whitney U test:

- `ctr` vs `srrm4_upfi` : 0.00168796
  mean: 1.0708 > 0.870257 , median: 1.0193 > 0.79553

#### 6.17 RATIO DOINTRON EXON GCC

Back to: Overview | ToC

Meaning: DOINTRON GCC / EXON GCC

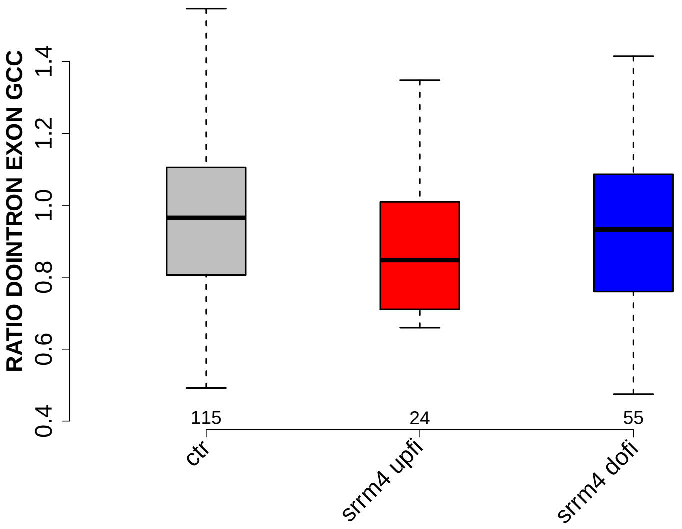

Significant results from Mann-Whitney U test:

- none

#### 6.18 RATIO DOEXON EXON GCC

Back to: Overview | ToC

Meaning: DOEXON GCC / EXON GCC

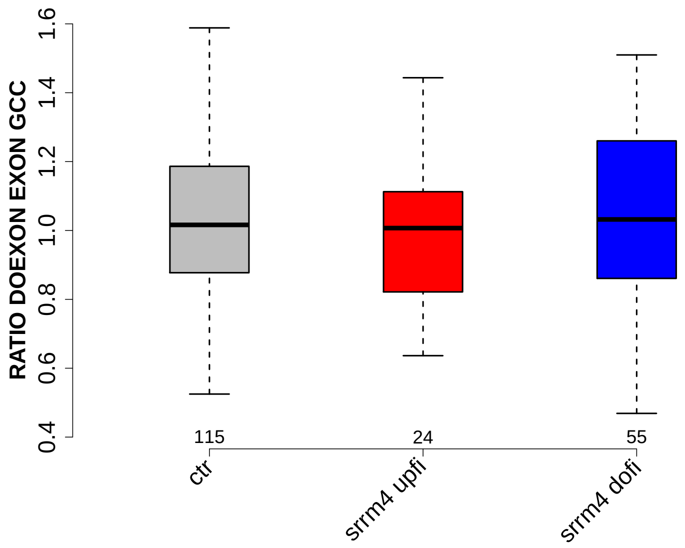

Significant results from Mann-Whitney U test:

- none

#### 6.19 SF1 HIGHESTSCORE 3SS UPINTRON

Back to: Overview | ToC

Meaning: highest score of a SF1 positon weight matrix trained with human data in the last 150 nt 3 prime intron positons of up-stream intron

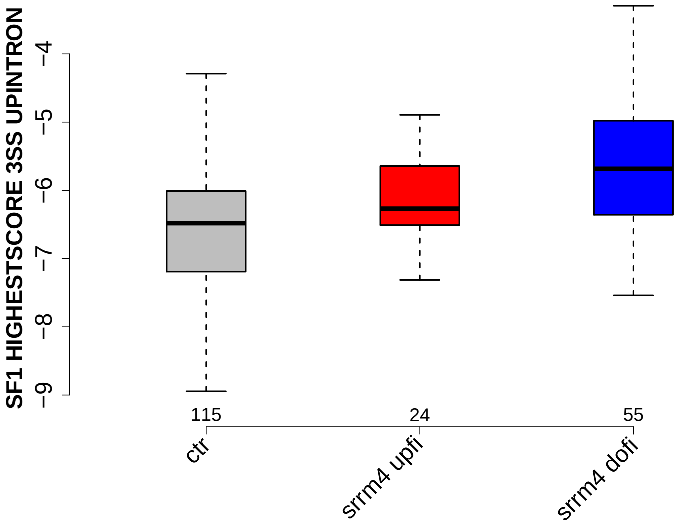

Significant results from Mann-Whitney U test:

- `ctr` vs `srrm4_upfi` : 0.0325769
  mean: -6.52951 < -6.13069 , median: -6.48026 < -6.26872
- `ctr` vs `srrm4_dofi` : 5.8465e-06
  mean: -6.52951 < -5.7077 , median: -6.48026 < -5.6844

#### 6.20 SF1 HIGHESTSCORE 3SS DOINTRON

Back to: Overview | ToC

Meaning: highest score of a SF1 positon weight matrix trained with human data in the last 150 nt 3 prime intron positons of down-stream intron

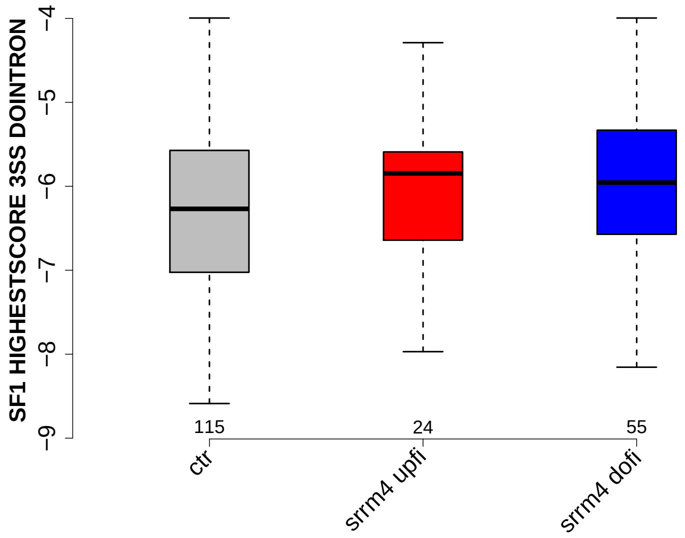

Significant results from Mann-Whitney U test:

- none

#### 6.21 UP 5SS 20INT10EX GCC

Back to: Overview | ToC

Meaning: GC content of up-stream 5ss sequence (20int+10ex positions)

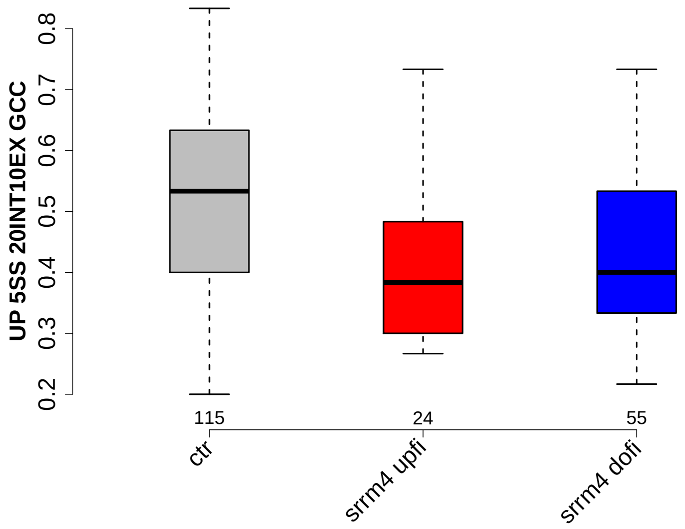

Significant results from Mann-Whitney U test:

- `ctr` vs `srrm4_upfi` : 0.00276321
  mean: 0.521884 > 0.425 , median: 0.533333 > 0.383333
- `ctr` vs `srrm4_dofi` : 0.000410572
  mean: 0.521884 > 0.433636 , median: 0.533333 > 0.4

##### 6.22 GCC 3SS 20INT10EX

Back to: Overview | ToC

Meaning: GC content of 3ss sequence (20int+10ex positions)

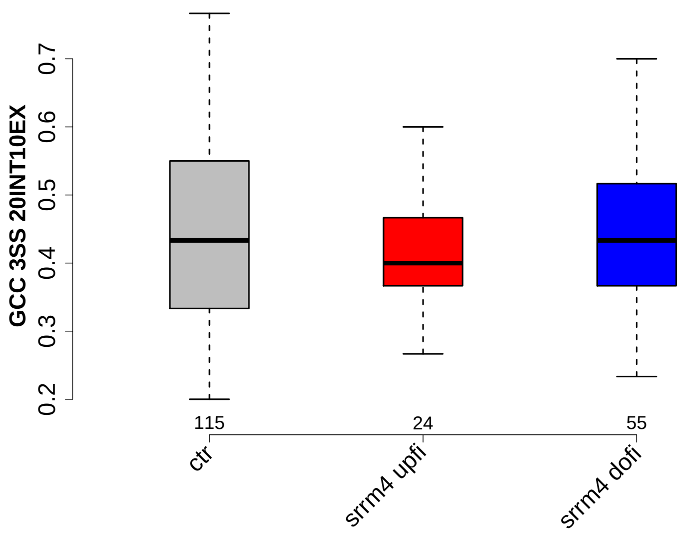

Significant results from Mann-Whitney U test:

- none

#### 6.23 GCC 5SS 20INT10EX

Back to: Overview | ToC

Meaning: GC content of 5ss sequence (20int+10ex positions)

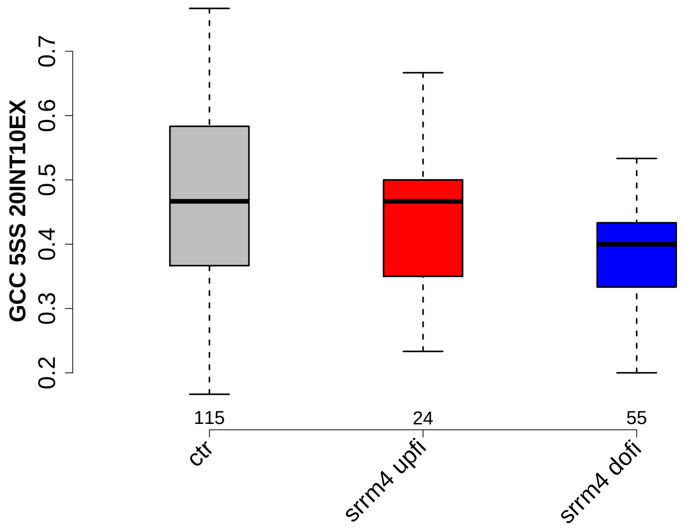

Significant results from Mann-Whitney U test:

- `ctr` vs `srrm4_dofi` : 0.000680785
  mean: 0.469855 > 0.39697 , median: 0.466667 > 0.4

#### 6.24 DO 3SS 20INT10EX GCC

Back to: Overview | ToC

Meaning: GC content of down-stream 3ss sequence (20int+10ex positions)

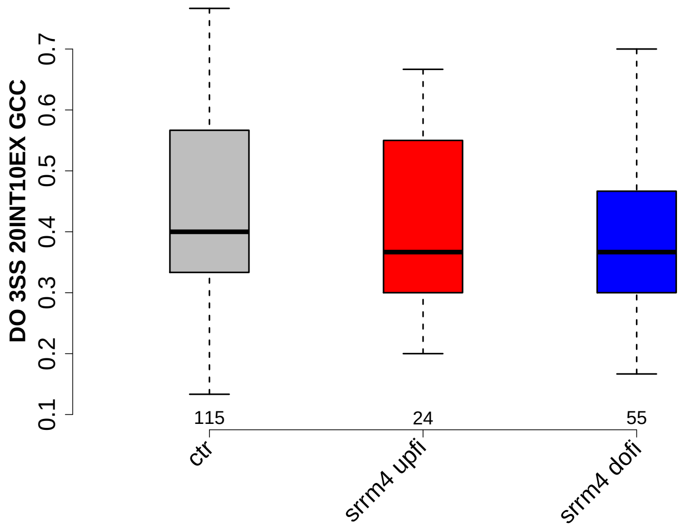

Significant results from Mann-Whitney U test:

- none

#### 6.25 MAXENTSCR HSAMODEL UPSTRM 5SS

Back to: Overview | ToC

Meaning: maximum entropy score of 5ss of up-stream exon using a model trained with human splice sites

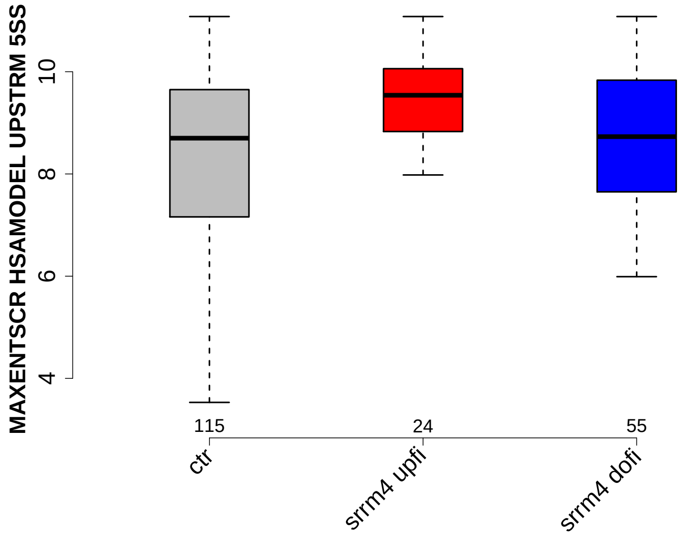

Significant results from Mann-Whitney U test:

- `ctr` vs `srrm4_upfi` : 0.00947595
  mean: 7.734 < 9.0471 , median: 8.7 < 9.54

#### 6.26 MAXENTSCR HSAMODEL 3SS

Back to: Overview | ToC

Meaning: maximum entropy score of 3ss using a model trained with human splice sites

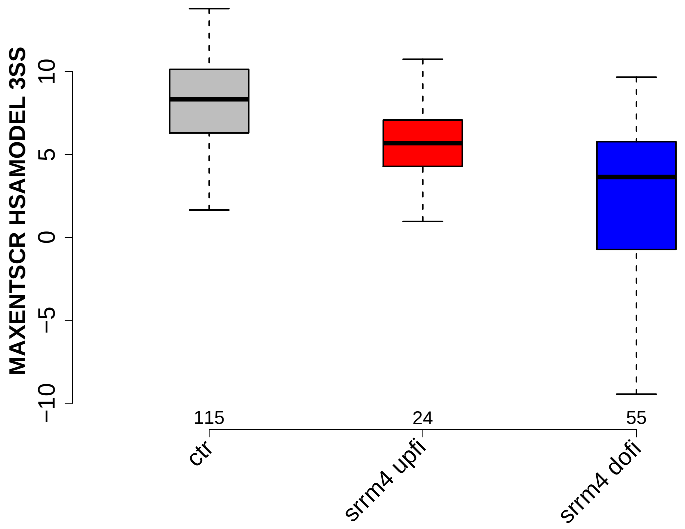

Significant results from Mann-Whitney U test:

- ctr vs srrm4_upfi : 0.000648238
  mean: 7.2993 > 4.3788 , median: 8.33 > 5.685
- ctr vs srrm4_dofi : 2.03641e-13
  mean: 7.2993 > 1.7489 , median: 8.33 > 3.64
- srrm4_upfi vs srrm4_dofi : 0.0103551
  mean: 4.3788 > 1.7489 , median: 5.685 > 3.64

#### 6.27 MAXENTSCR HSAMODEL 5SS

Back to: Overview | ToC

Meaning: maximum entropy score of 5ss using a model trained with human splice sites

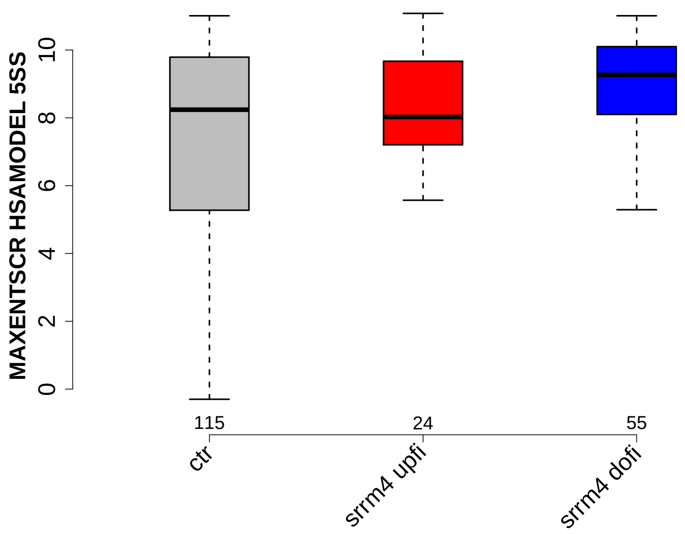

Significant results from Mann-Whitney U test:
- `ctr` vs `srrm4_dofi` : 0.00475608
  mean: 6.5228 < 8.834 , median: 8.24 < 9.27

#### 6.28 MAXENTSCR HSAMODEL DOWNSTRM 3SS

Back to: Overview | ToC

Meaning: maximum entropy score of 3ss of down-stream exon using a model trained with human splice sites

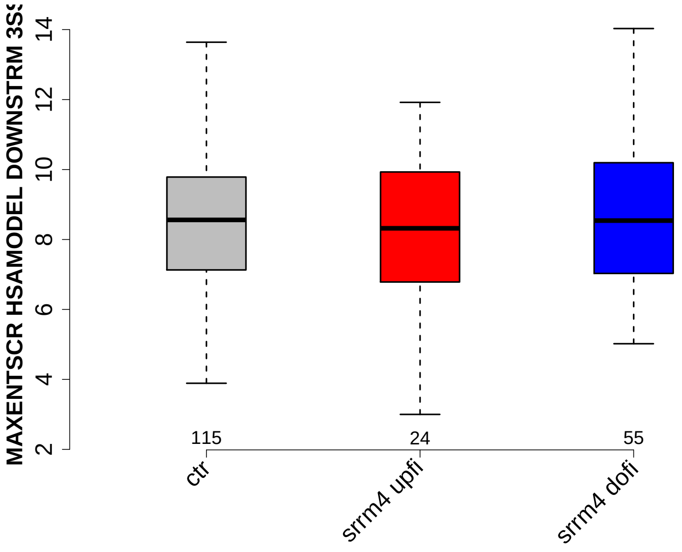

Significant results from Mann-Whitney U test:

- none

#### 6.29 DIST FROM MAXBP TO 3SS UPINTRON

Back to: Overview | ToC

Meaning: distance to 3ss of best precited BP

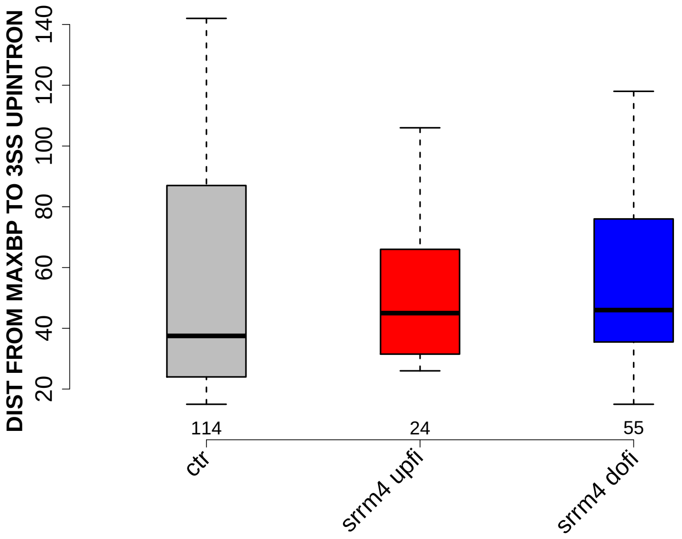

Significant results from Mann-Whitney U test:

- none

#### 6.30 SCORE FOR MAXBP SEQ UPINTRON

Back to: Overview | ToC

Meaning: BP sequence score of best predicted BP

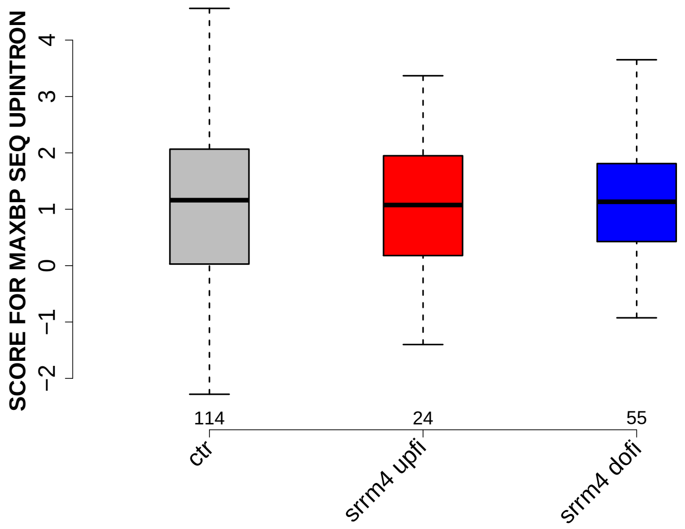

Significant results from Mann-Whitney U test:

- none

#### 6.31 PYRIMIDINECONT MAXBP UPINTRON

Back to: Overview | ToC

Meaning: Pyrimidine content between the BP adenine and the 3 prime splice site for best BP

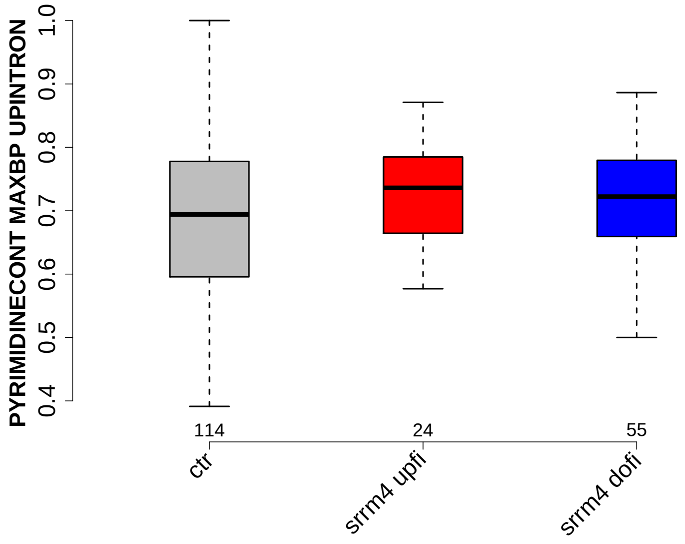

Significant results from Mann-Whitney U test:

- none

#### 6.32 POLYPYRITRAC OFFSET MAXBP UPINTRON

Back to: Overview | ToC

Meaning: Polypyrimidine track offset relative to the BP adenine for best BP

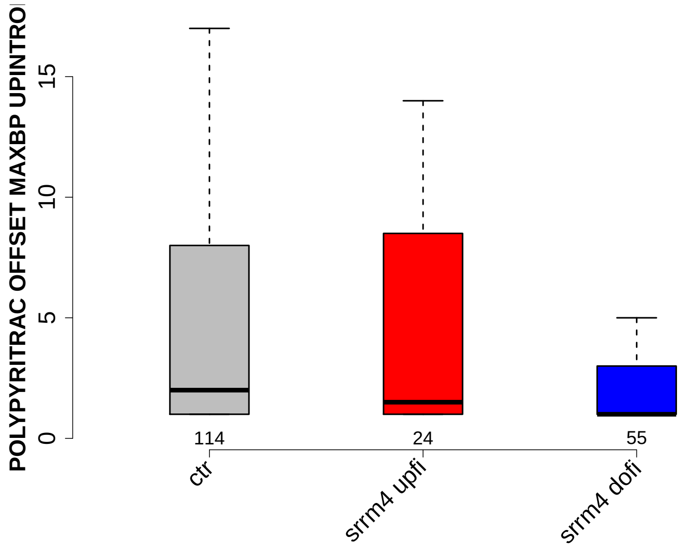

Significant results from Mann-Whitney U test:

- `ctr` vs `srrm4_dofi` : 0.0128239
  mean: 5.2807 > 3.0364 , median: 2 > 1

#### 6.33 POLYPYRITRAC LEN MAXBP UPINTRON

Back to: Overview | ToC

Meaning: Polypyrimidine track length for best BP

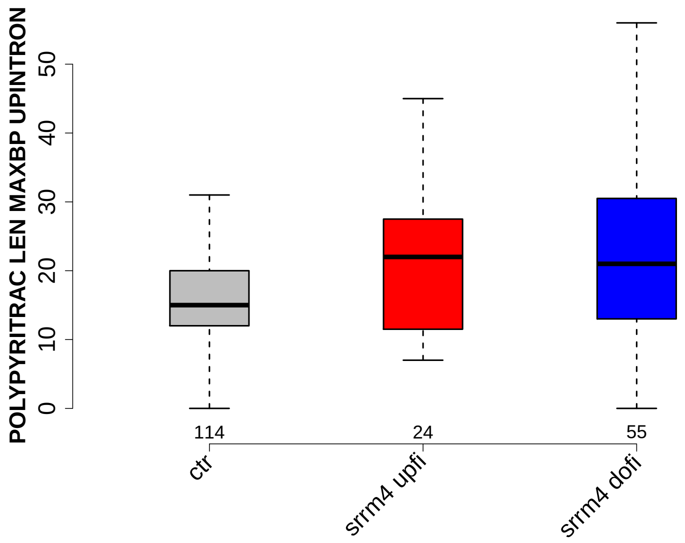

Significant results from Mann-Whitney U test:

- `ctr` vs `srrm4_upfi` : 0.0476083
  mean: 16.2193 < 21.7083 , median: 15 < 22
- `ctr` vs `srrm4_dofi` : 0.00348289
  mean: 16.2193 < 22.0182 , median: 15 < 21

#### 6.34 POLYPYRITRAC SCORE MAXBP UPINTRON

Back to: Overview | ToC

Meaning: Polypyrimidine track score for best BP

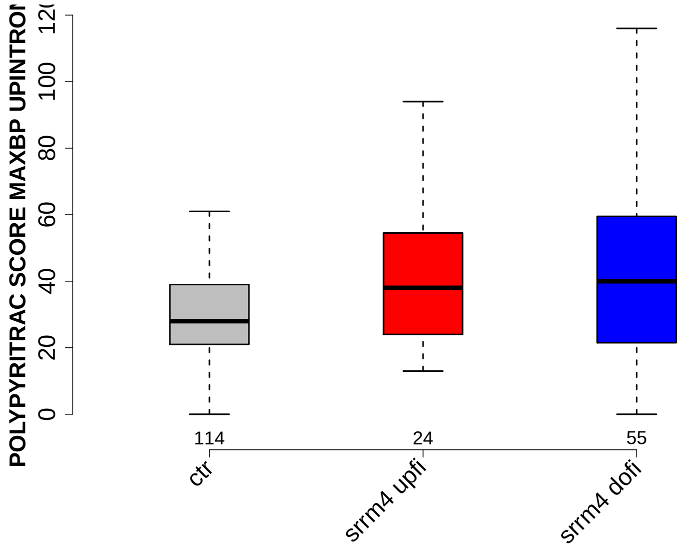

Significant results from Mann-Whitney U test:

- `ctr` vs `srrm4_upfi` : 0.0205878
  mean: 30.6667 < 43.8333 , median: 28 < 38
- `ctr` vs `srrm4_dofi` : 0.0036507
  mean: 30.6667 < 42.6 , median: 28 < 40

#### 6.35 BPSCORE MAXBP UPINTRON

Back to: Overview | ToC

Meaning: SVM classification score of best BP

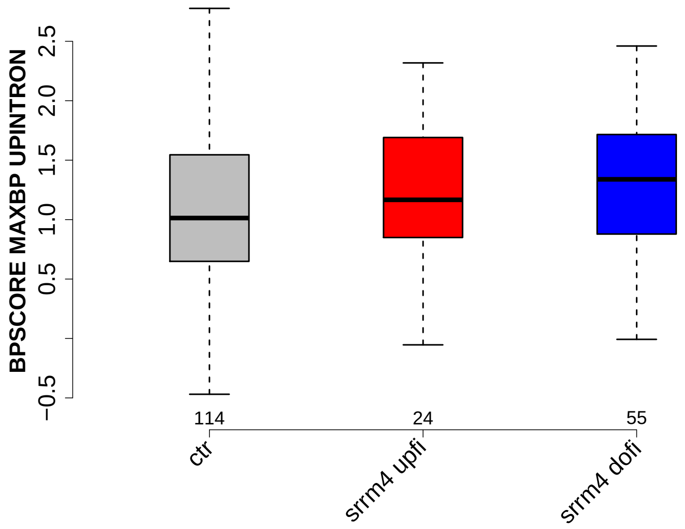

Significant results from Mann-Whitney U test:

- `ctr` vs `srrm4_dofi` : 0.020881
  mean: 1.0135 < 1.2932 , median: 1.0134 < 1.3389

#### 6.36 NUM PREDICTED BPS UPINTRON

Back to: Overview | ToC

Meaning: number of all predicted BPs which have a positive BP score

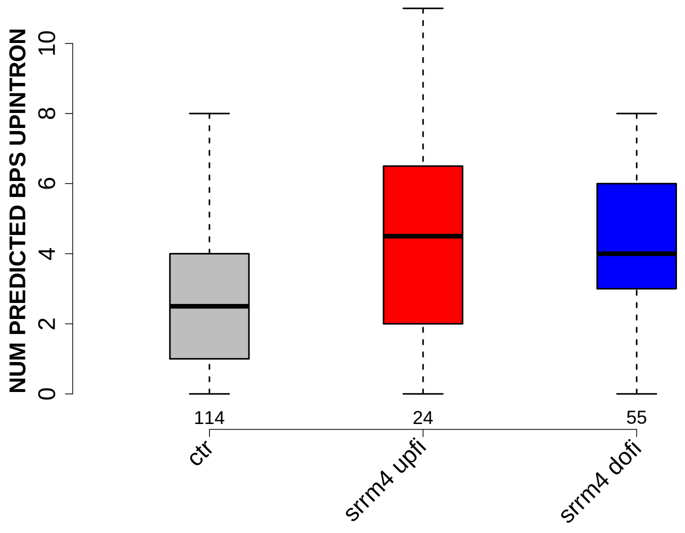

Significant results from Mann-Whitney U test:

- `ctr` vs `srrm4_upfi` : 0.00736109
  mean: 2.9474 < 4.625 , median: 2.5 < 4.5
- `ctr` vs `srrm4_dofi` : 0.000610198
  mean: 2.9474 < 4.0909 , median: 2.5 < 4

#### 6.37 MEDIAN DIST FROM BP TO 3SS UPINTRON

Back to: Overview | ToC

Meaning: like DIST FROM MAXBP TO 3SS but median over top-3 predicted BPs

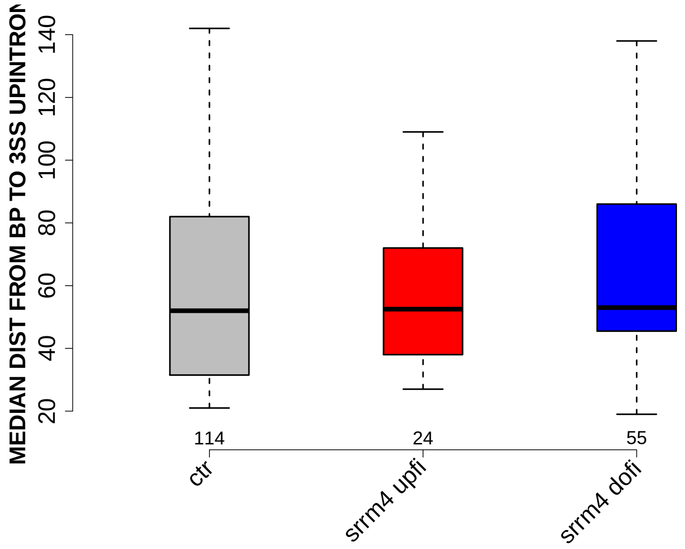

Significant results from Mann-Whitney U test:

- none

#### 6.38 MEDIAN SCORE FOR BPSEQ UPINTRON

Back to: Overview | ToC

Meaning: like SCORE FOR MAXBP SEQ but median over top-3 predicted BPs

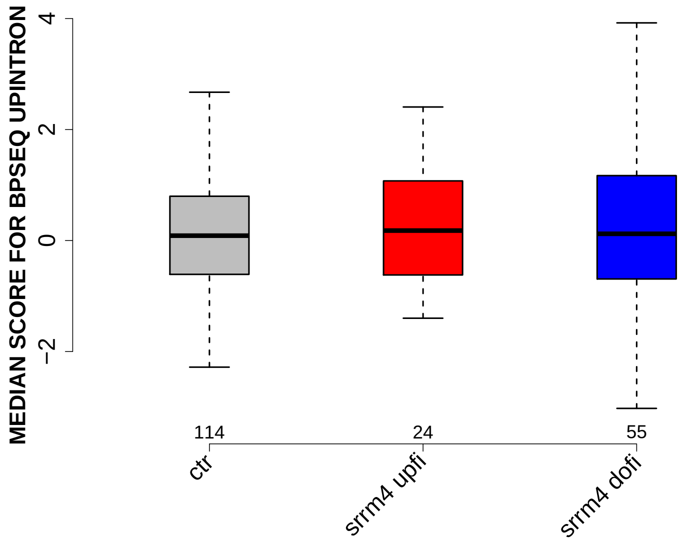

Significant results from Mann-Whitney U test:

- none

#### 6.39 MEDIAN PYRIMIDINECONT UPINTRON

Back to: Overview | ToC

Meaning: like PYRIMIDINECONT MAXBP but median over top-3 predicted BPs

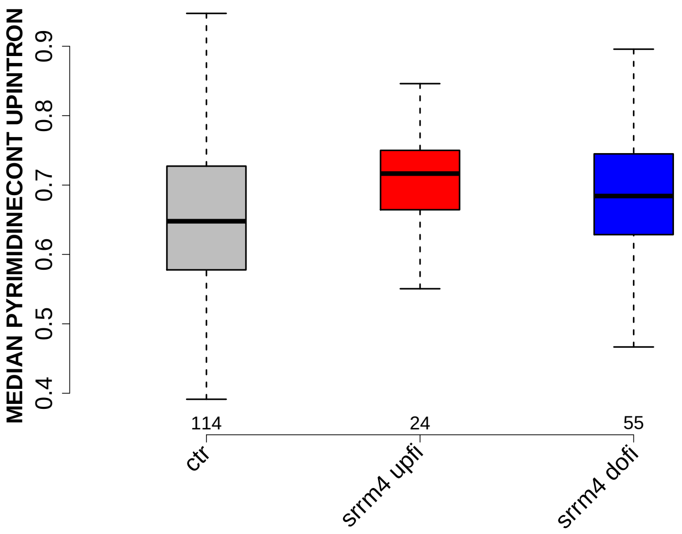

Significant results from Mann-Whitney U test:

- `ctr` vs `srrm4_upfi` : 0.0102522
  mean: 0.653347 < 0.707328 , median: 0.647854 < 0.716518

#### 6.40 MEDIAN POLYPYRITRAC OFFSET UPINTRON

Back to: Overview | ToC

Meaning: like POLYPYRITRAC OFFSET MAXBP but median over top-3 predicted BPs

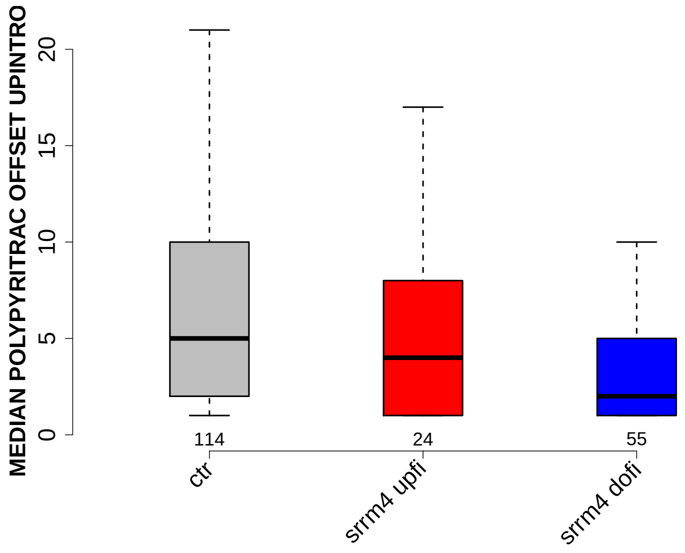

Significant results from Mann-Whitney U test:

- `ctr` vs `srrm4_dofi` : 0.000455268
  mean: 9.1404 > 4.0182 , median: 5 > 2

#### 6.41 MEDIAN POLYPYRITRAC LEN UPINTRON

Back to: Overview | ToC

Meaning: like POLYPYRITRAC LEN MAXBP but median over top-3 predicted BPs

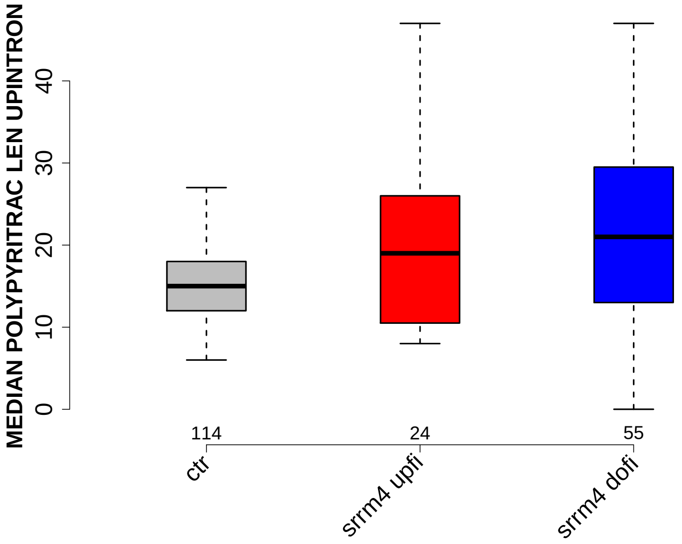

Significant results from Mann-Whitney U test:

- `ctr` vs `srrm4_dofi` : 0.000859434
  mean: 15.8772 < 21.9273 , median: 15 < 21

#### 6.42 MEDIAN POLYPYRITRAC SCORE UPINTRON

Back to: Overview | ToC

Meaning: like POLYPYRITRAC SCORE MAXBP but median over top-3 predicted BPs

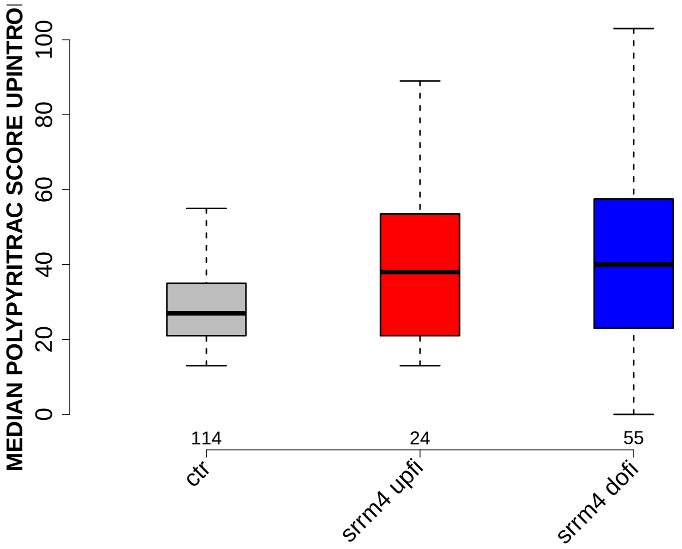

Significant results from Mann-Whitney U test:

- `ctr` vs `srrm4_dofi` : 0.000198875
  mean: 29.5702 < 42.5091 , median: 27 < 40

#### 6.43 MEDIAN BPSCORE UPINTRON

Back to: Overview | ToC

Meaning: like BPSCORE MAXBP but median over top-3 predicted BPs

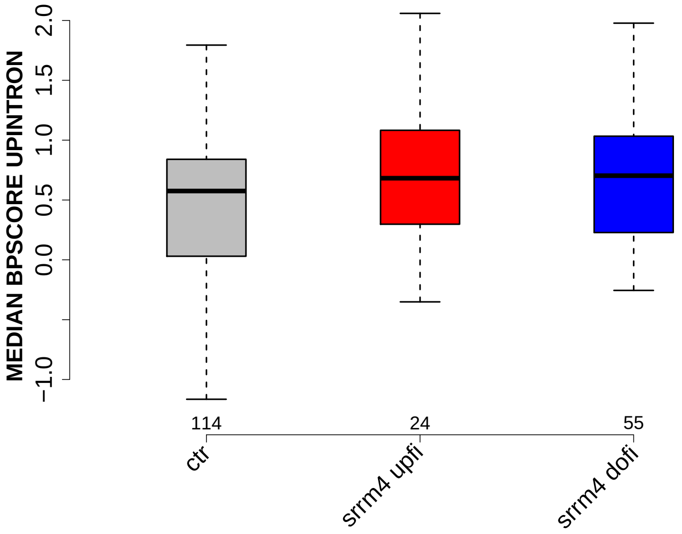

Significant results from Mann-Whitney U test:

- `ctr` vs `srrm4_dofi` : 0.0288168
  mean: 0.303545 < 0.680841 , median: 0.575111 < 0.703222

#### 6.44 DIST FROM MAXBP TO 3SS DOINTRON

Back to: Overview | ToC

Meaning: distance to 3ss of best precited BP

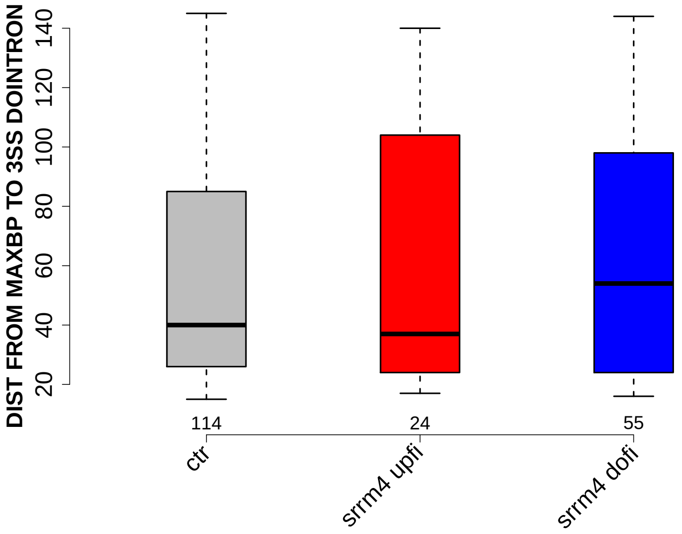

Significant results from Mann-Whitney U test:

- none

#### 6.45 SCORE FOR MAXBP SEQ DOINTRON

Back to: Overview | ToC

Meaning: BP sequence score of best predicted BP

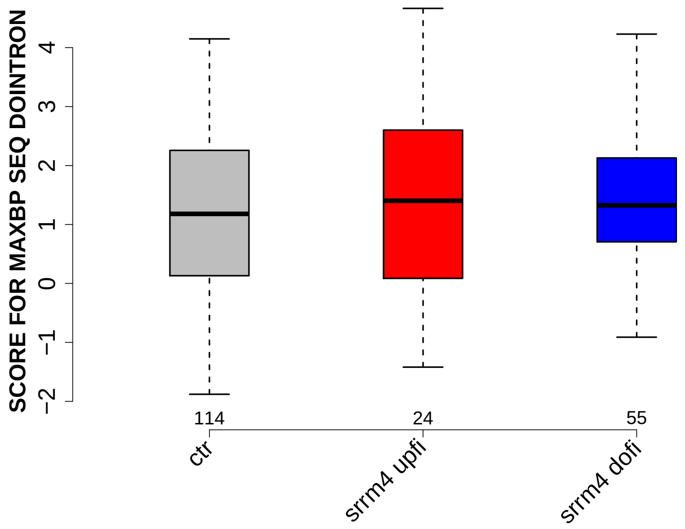

Significant results from Mann-Whitney U test:

- none

#### 6.46 PYRIMIDINECONT MAXBP DOINTRON

Back to: Overview | ToC

Meaning: Pyrimidine content between the BP adenine and the 3 prime splice site for best BP

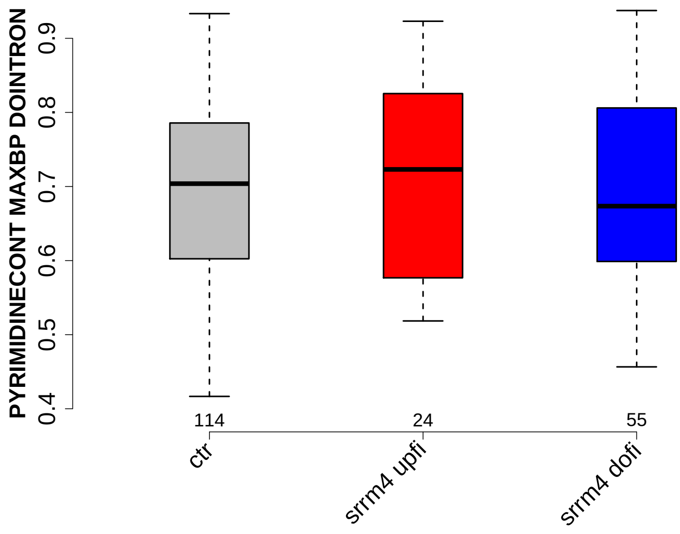

Significant results from Mann-Whitney U test:

- none

#### 6.47 POLYPYRITRAC OFFSET MAXBP DOINTRON

Back to: Overview | ToC

Meaning: Polypyrimidine track offset relative to the BP adenine for best BP

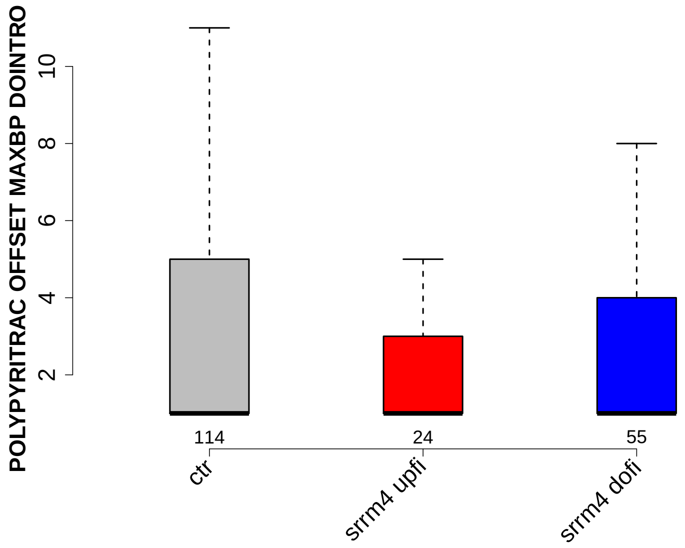

Significant results from Mann-Whitney U test:

- none

#### 6.48 POLYPYRITRAC LEN MAXBP DOINTRON

Back to: Overview | ToC

Meaning: Polypyrimidine track length for best BP

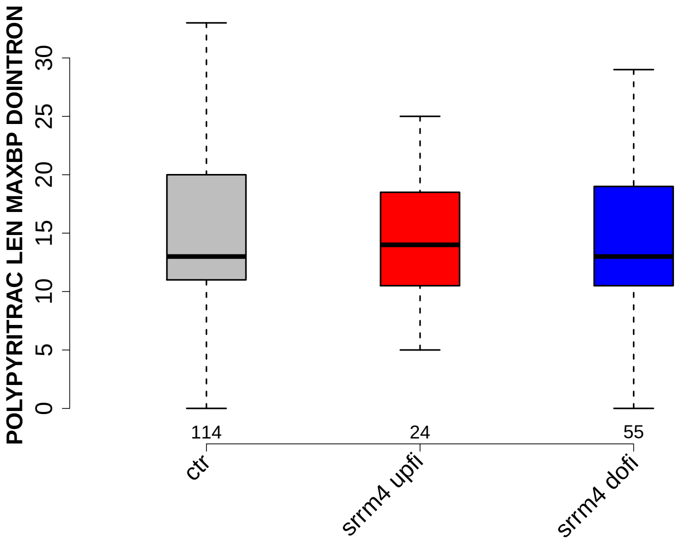

Significant results from Mann-Whitney U test:

- none

#### 6.49 POLYPYRITRAC SCORE MAXBP DOINTRON

Back to: Overview | ToC

Meaning: Polypyrimidine track score for best BP

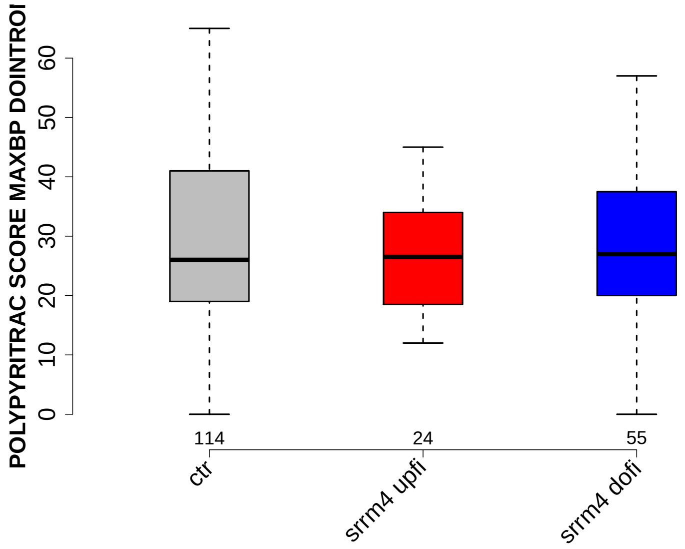

Significant results from Mann-Whitney U test:

- none

#### 6.50 BPSCORE MAXBP DOINTRON

Back to: Overview | ToC

Meaning: SVM classification score of best BP

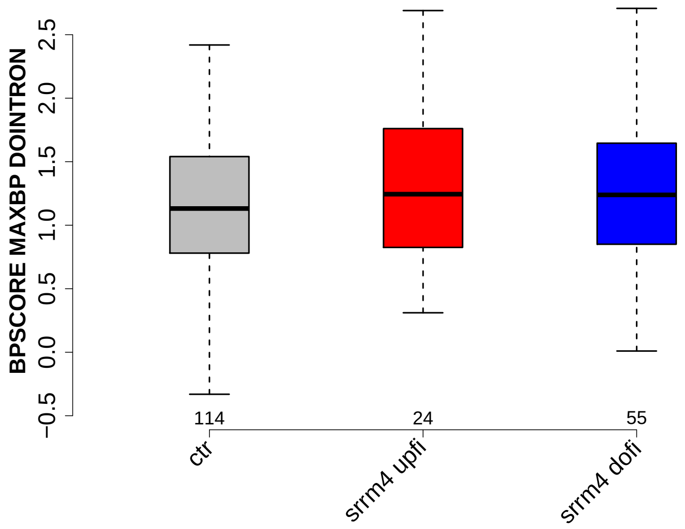

Significant results from Mann-Whitney U test:

- none

#### 6.51 NUM PREDICTED BPS DOINTRON

Back to: Overview | ToC

Meaning: number of all predicted BPs which have a positive BP score

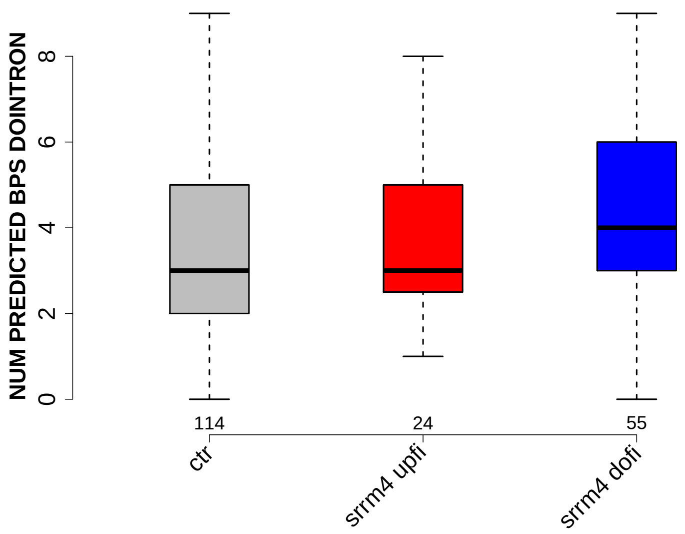

Significant results from Mann-Whitney U test:

- `ctr` vs `srrm4_dofi` : 0.0176817
  mean: 3.4211 < 4.2727 , median: 3 < 4

#### 6.52 MEDIAN DIST FROM BP TO 3SS DOINTRON

Back to: Overview | ToC

Meaning: like DIST FROM MAXBP TO 3SS but median over top-3 predicted BPs

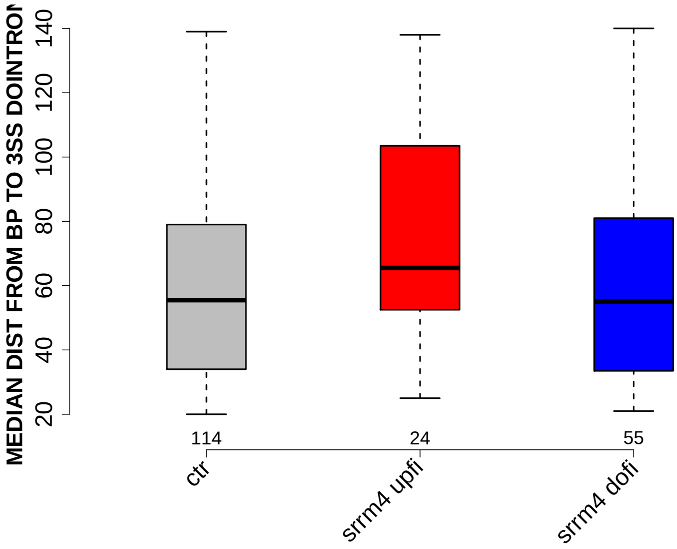

Significant results from Mann-Whitney U test:

- none

#### 6.53 MEDIAN SCORE FOR BPSEQ DOINTRON

Back to: Overview | ToC

Meaning: like SCORE FOR MAXBP SEQ but median over top-3 predicted BPs

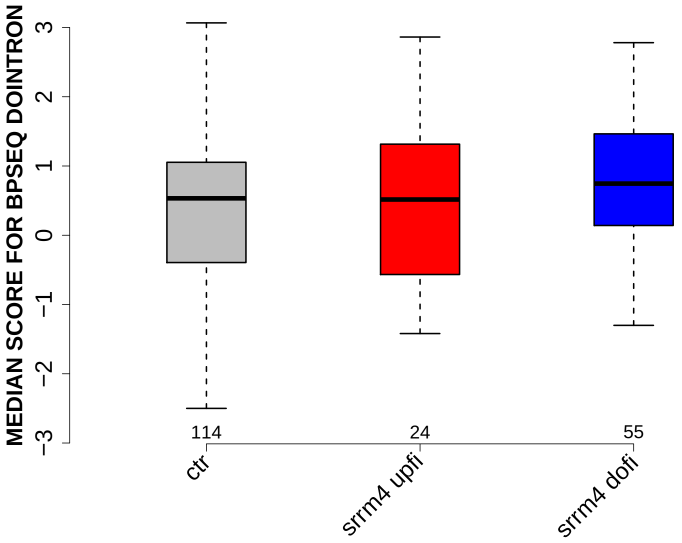

Significant results from Mann-Whitney U test:

- none

#### 6.54 MEDIAN PYRIMIDINECONT DOINTRON

Back to: Overview | ToC

Meaning: like PYRIMIDINECONT MAXBP but median over top-3 predicted BPs

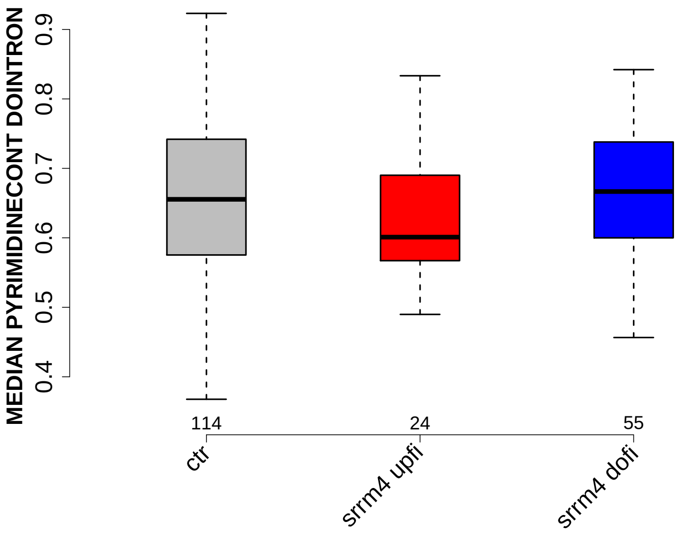

Significant results from Mann-Whitney U test:

- none

#### 6.55 MEDIAN POLYPYRITRAC OFFSET DOINTRON

Back to: Overview | ToC

Meaning: like POLYPYRITRAC OFFSET MAXBP but median over top-3 predicted BPs

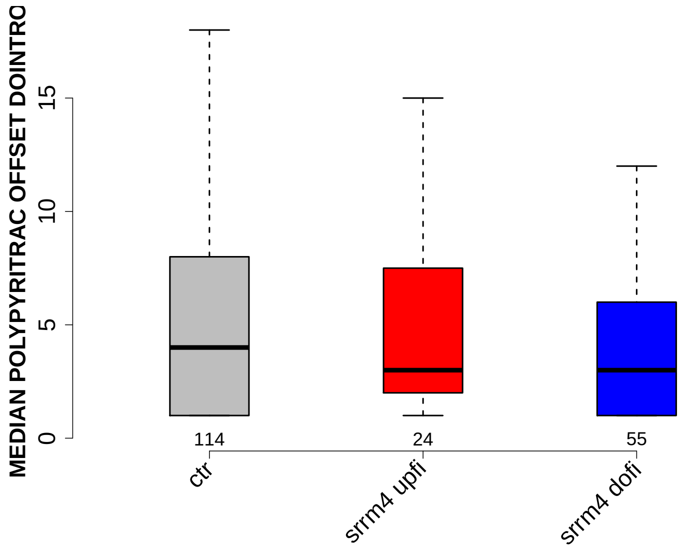

Significant results from Mann-Whitney U test:
- none

#### 6.56 MEDIAN POLYPYRITRAC LEN DOINTRON

Back to: Overview | ToC

Meaning: like POLYPYRITRAC LEN MAXBP but median over top-3 predicted BPs

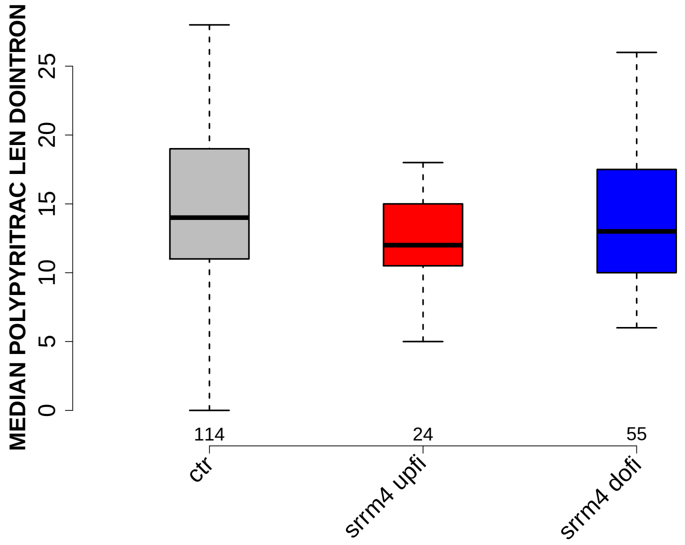

Significant results from Mann-Whitney U test:

- none

#### 6.57 MEDIAN POLYPYRITRAC SCORE DOINTRON

Back to: Overview | ToC

Meaning: like POLYPYRITRAC SCORE MAXBP but median over top-3 predicted BPs

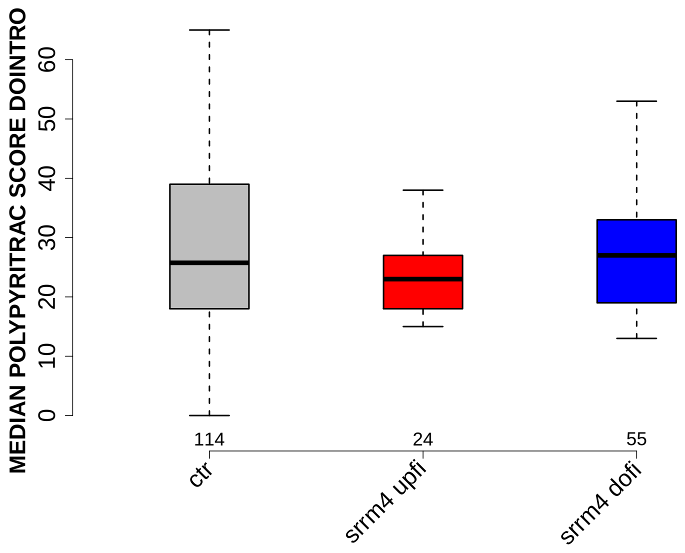

Significant results from Mann-Whitney U test:

- none

#### 6.58 MEDIAN BPSCORE DOINTRON

Back to: Overview | ToC

Meaning: like BPSCORE MAXBP but median over top-3 predicted BPs

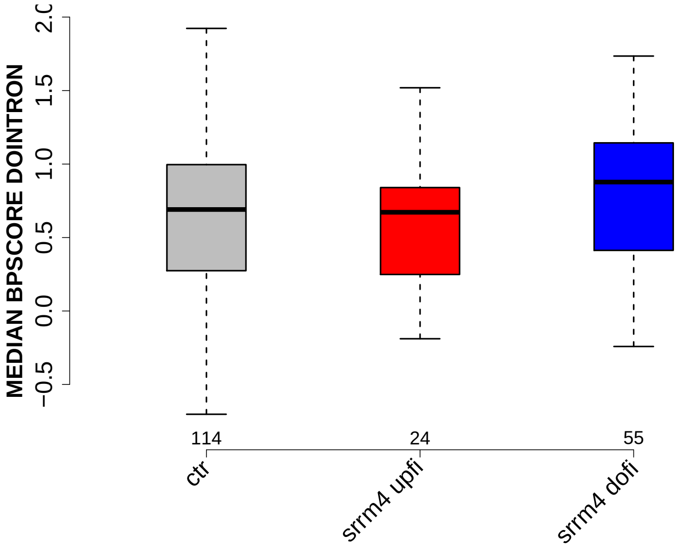

Significant results from Mann-Whitney U test:

- `ctr` vs `srrm4_dofi` : 0.0471817
  mean: 0.464177 < 0.768531 , median: 0.690675 < 0.877449

#### 6.59 MEDIAN TR LENGTH

Back to: Overview | ToC

Meaning: median length of transcripts the exon occurs in

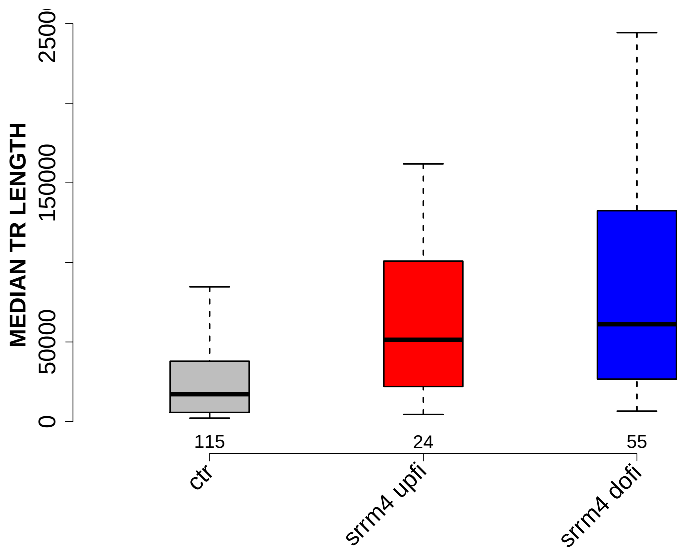

Significant results from Mann-Whitney U test:

- `ctr` vs `srrm4_upfi` : 0.00136682
  mean: 34533.1217 < 85219.8333 , median: 17235 < 51337.75
- `ctr` vs `srrm4_dofi` : 3.80683e-09
  mean: 34533.1217 < 103036.1818 , median: 17235 < 61218

#### 6.60 MEDIAN EXON NUMBER

Back to: Overview | ToC

Meaning: ... of transcripts where exon was found in

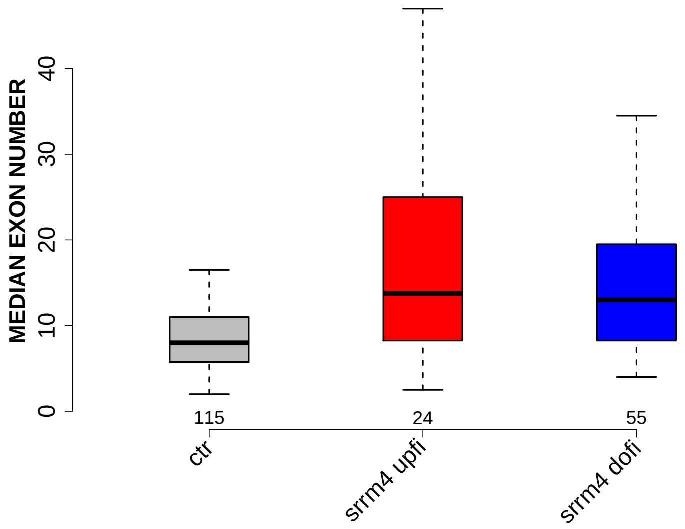

Significant results from Mann-Whitney U test:

- `ctr` vs `srrm4_upfi` : 0.00111351
  mean: 9.5522 < 19.1042 , median: 8 < 13.75
- `ctr` vs `srrm4_dofi` : 1.29223e-05
  mean: 9.5522 < 14.7364 , median: 8 < 13

#### 6.61 EXON MEDIANRELATIVERANK

Back to: Overview | ToC

Meaning: relative rank = rank / number of all exons in transcript, is between 0 and 1

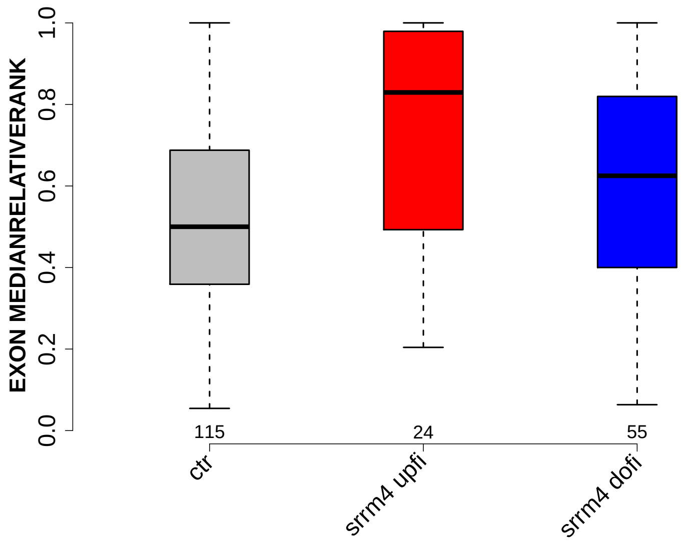

Significant results from Mann-Whitney U test:

- ctr vs srrm4_upfi : 0.00329206
  mean: 0.531743 < 0.712914 , median: 0.5 < 0.82938

#### 6.62 EXON MEDIANRELATIVERANK 3BINS

Back to: Overview | ToC

Meaning: median bin into which EXON MEDIANRELATIVERANK falls when binning 0-1 into 3 bins

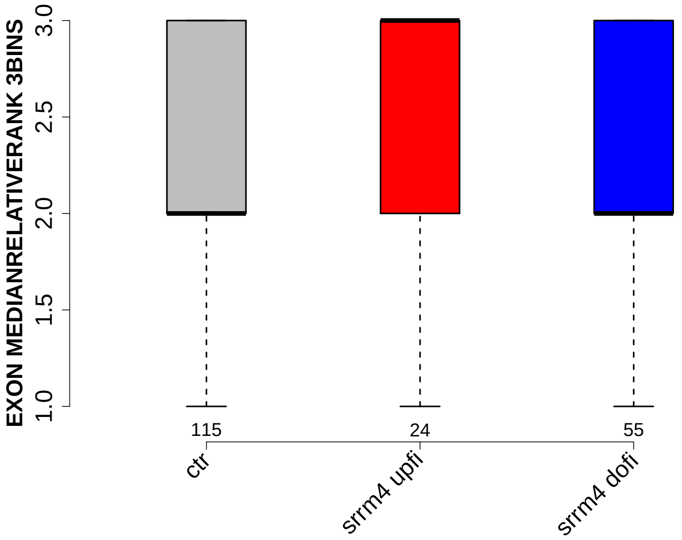

Significant results from Mann-Whitney U test:

- `ctr` vs `srrm4_upfi` : 0.0266448
  mean: 2.1304 < 2.4583 , median: 2 < 3

#### 6.63 EXON MEDIANRELATIVERANK 5BINS

Back to: Overview | ToC

Meaning: similar to EXON MEDIANRELATIVERANK 3BINS with 5 bins

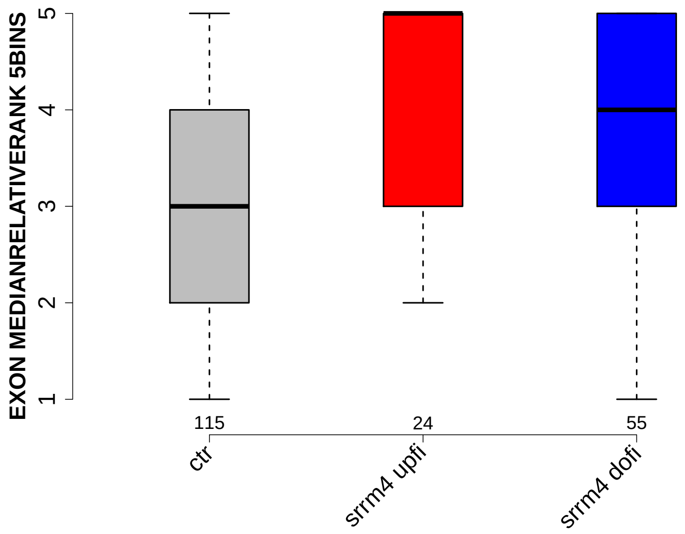

Significant results from Mann-Whitney U test:

- `ctr` vs `srrm4_upfi` : 0.00242098
  mean: 3.1565 < 4 , median: 3 < 5

#### 6.64 EXON MEDIANRELATIVERANK 10BINS

Back to: Overview | ToC

Meaning: similar to EXON MEDIANRELATIVERANK 3BINS with 10 bins

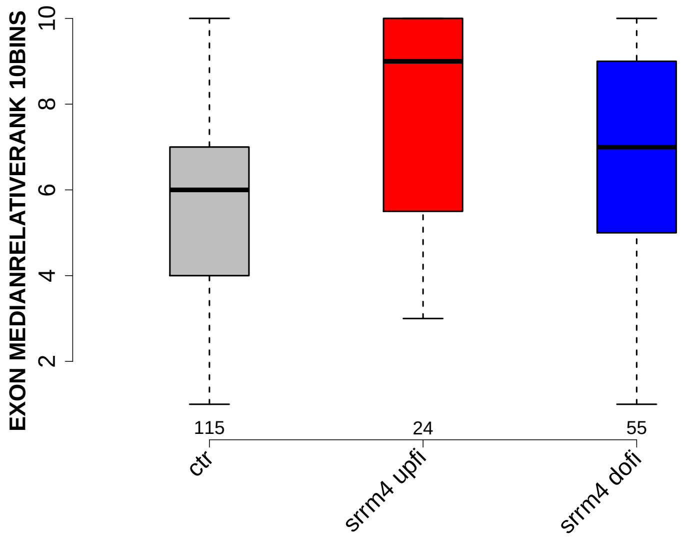

Significant results from Mann-Whitney U test:

- `ctr` vs `srrm4_upfi` : 0.00266538
  mean: 5.8348 $<$ 7.5833 , median: 6 $<$ 9

#### 6.65 NTRS ALL FOR GENE

Back to: Overview | ToC

Meaning: number of transcripts of gene where the exon was found in

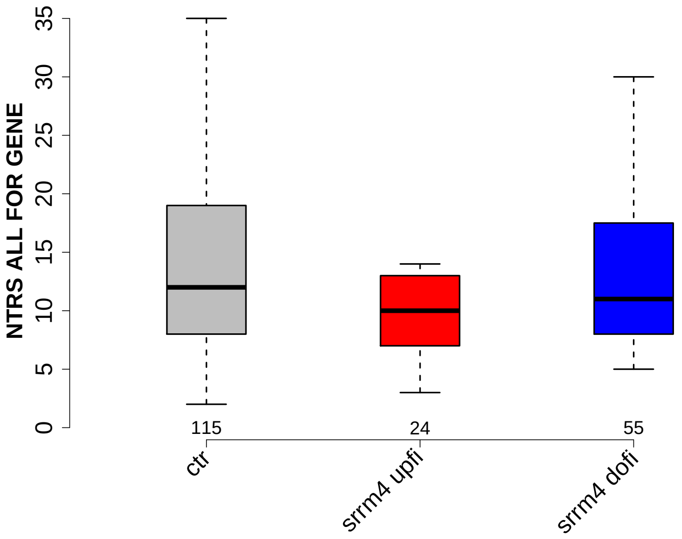

Significant results from Mann-Whitney U test:

- none

#### 6.66 PROP FIRST EXON

Back to: Overview | ToC

Meaning: NTRS WITH EXON AS FIRST EXON / NTRS WITH EXON

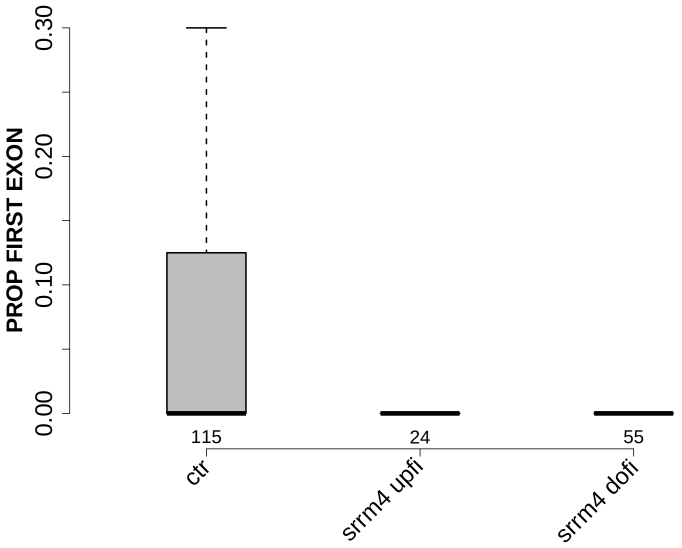

Significant results from Mann-Whitney U test:

- `ctr` vs `srrm4_upfi` : 0.00163247
  mean: 0.0848067 > 0.0104167 , median: 0 = 0
- `ctr` vs `srrm4_dofi` : 0.00587701
  mean: 0.0848067 > 0.0286305 , median: 0 = 0

#### 6.67 PROP LAST EXON

Back to: Overview | ToC

Meaning: NTRS WITH EXON AS LAST EXON / NTRS WITH EXON

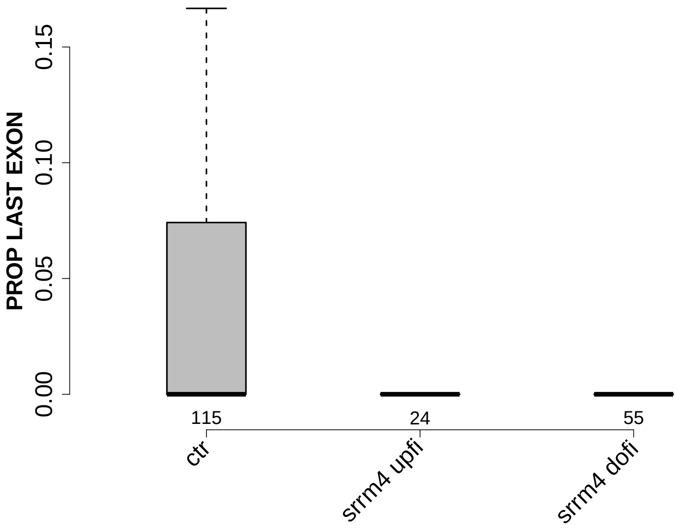

Significant results from Mann-Whitney U test:

- `ctr` vs `srrm4_dofi` : 0.000537025
  mean: 0.0558943 > 0.00854257 , median: 0 = 0

#### 6.68 PROP INTERNAL EXON

Back to: Overview | ToC

Meaning: NTRS WITH EXON AS INTERNAL EXON / NTRS WITH EXON

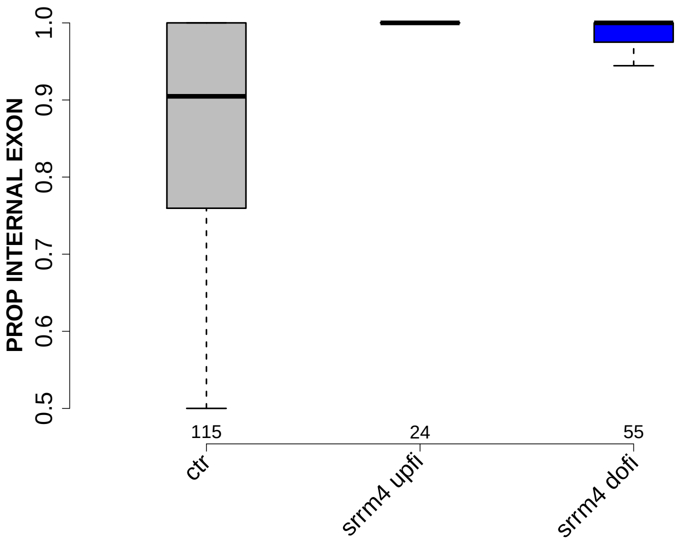

Significant results from Mann-Whitney U test:

- `ctr` vs `srrm4_upfi` : 0.000390521
  mean: 0.860024 $<$ 0.96875 , median: 0.904762 $<$ 1
- `ctr` vs `srrm4_dofi` : 9.09738e-06
  mean: 0.860024 $<$ 0.962827 , median: 0.904762 $<$ 1

#### 6.69 PROP EXON IN UTR

Back to: Overview | ToC

Meaning: NTRS WITH EXON IN UTR / NTRS WITH EXON

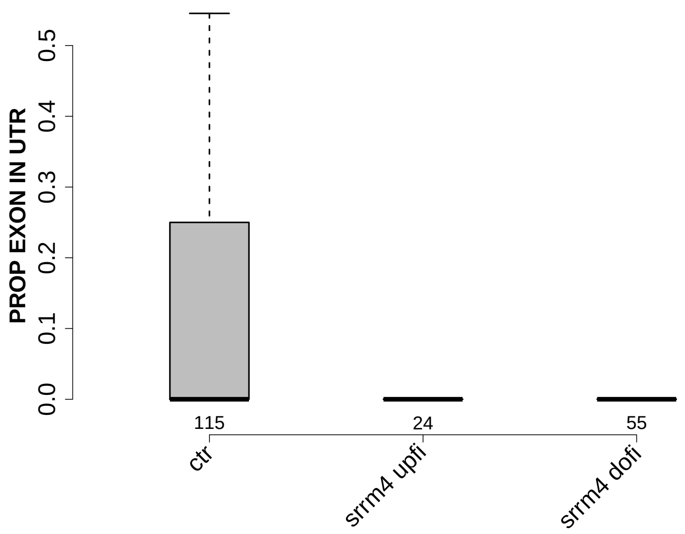

Significant results from Mann-Whitney U test:

- `ctr` vs `srrm4_upfi` : 0.019758
  mean: 0.149222 > 0.0590278 , median: 0 = 0
- `ctr` vs `srrm4_dofi` : 1.06979e-05
  mean: 0.149222 > 0.0251787 , median: 0 = 0
